# The fitness landscape of the *E.coli lac* operator is highly rugged in two different environments

**DOI:** 10.1101/2025.07.23.666252

**Authors:** Gopinath Chattopadhyay, Andrei Papkou, Andreas Wagner

**Affiliations:** Institute for Evolutionary Biology and Environmental Studies, University of Zurich, Zurich, Switzerland; Swiss Institute of Bioinformatics (SIB), University of Lausanne, Switzerland; The Santa Fe Institute, Santa Fe, New Mexico

## Abstract

We know little about the fitness landscapes of bacterial operators, regulatory DNA elements that are crucial to regulate metabolic genes like those of the *lac* operon for lactose utilization. For example, we do not know whether adaptive evolution could easily create strong operators from weak ones or from non-regulatory DNA. To find out, we used CRISPR-Cas-assisted genome editing, bulk competition, and high-throughput sequencing to map the fitness landscape of more than 140,000 *lac* operator variants in two chemical environments that harbor lactose or glycerol as sole carbon sources. Both landscapes are highly rugged and contain thousands of fitness peaks, which allow only 2 percent of evolving populations to reach a high fitness peak. The landscapes share only 15 percent of fitness peaks. Our work illustrates that landscape ruggedness caused by epistasis can represent an important obstacle to adaptive evolution of regulatory sequences. It also shows that a simple environmental change can substantially affect fitness landscape topography.

## Introduction

A fitness landscape is an analogue of a physical landscape, in which each location corresponds to a genotype. The elevation at each location corresponds to the fitness of organisms with this genotype (1–3). While experimental studies have mapped fitness landscapes across environments, they have done so mainly for few genotypes of proteins (4–10). Such studies have provided insights into how proteins adapt to shifting selection pressures. Much less is known about the fitness landscapes of regulatory DNA (11–15).

Regulation of gene expression is fundamental to all cellular life. Gene expression carries a cost, because it consumes resources for mRNA and protein production (16–19). In microbes, this cost can reduce growth rate and fitness (16,19,20). For a gene to persist in a genome, its benefits must outweigh its costs in at least some environments (18,21,22). Both benefits and costs depend on a gene’s expression (23–26). To attain high fitness, an organism must regulate gene expression in an environment-specific manner. This regulation is primarily exerted by transcription factors (TFs), proteins that bind specific DNA sequences called transcription factor binding sites (TFBSs) near a gene (27,28).

TFBSs are subject to adaptive evolution, because DNA mutations can change their affinity to a cognate transcription factor, and thus gene regulation and fitness. Mapping the fitness landscape of a TFBS can help to understand both its adaptive evolution and the emergence of new binding sites (de novo evolution) from non-regulatory DNA (12,13,29). Here we map the fitness landscape of a prominent prokaryotic TFBS and study adaptive evolution on this landscape. To find out whether landscape topography depends on the environment, we map the fitness landscape of this TFBS in two distinct environments: one where a gene needs to be repressed because it conveys no benefit, and another where this same gene needs to be activated (de-repressed).

We chose the catabolic lactose (*lac*) operon of *Escherichia coli* as a model system for studying the fitness landscape of prokaryotic gene regulation. The *lac* operon comprises the genes *lacZ*, *lacY*, and *lacA*, whose protein products are essential to metabolize lactose. The operon is thus important in environments where lactose is the sole carbon source (30–33). The operon is regulated by the constitutively expressed LacI repressor protein, encoded by the *lacI* gene (Figure 1a). In the absence of lactose, LacI binds tightly to the *lac* operator, inhibits RNA polymerase binding, and represses the transcription of the *lac* genes (Figure 1a). In the presence of lactose LacI undergoes an allosteric shift, and detaches from the operator (Figure 1a), enabling RNA polymerase to transcribe the *lac* genes, which allows cells to extract energy and carbon from lactose (34–37). The *lac* operon is also regulated by the CAP/CRP (Catabolite activator protein/ cAMP receptor protein) activator, which binds a CAP binding site to enhance transcription when lactose is available and glucose absent (Figure 1a).

**Figure 1.**
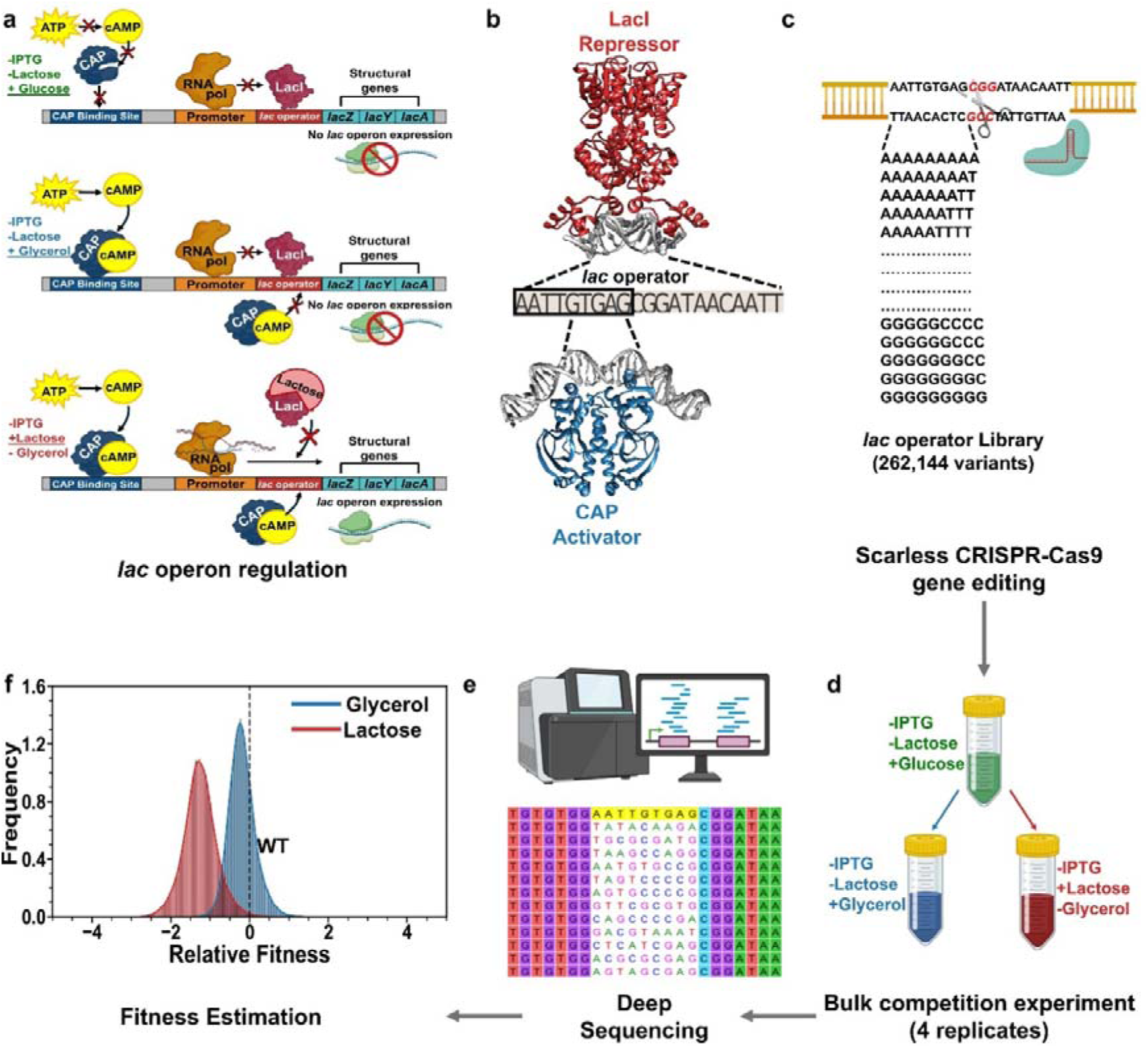
A minimally invasive approach to study the adaptive landscapes of prokaryotic gene regulation. (a) Regulation of the *lac* operon. In glucose, the CAP activator (blue) is inactive, and the Lac repressor (red) blocks transcription. In glycerol, the cAMP-CAP complex activates CAP, but transcription remains blocked by the repressor. In lactose, the repressor is released, and the cAMP-CAP complex mostly enhances RNA synthesis, but may also bind the operator directly (31,38). (b) Illustration of the *lac* operator site, with the positions chosen for mutagenesis marked by a black rectangle. The LacI repressor (red) binds to this 21 bp operator sequence in the absence of lactose (Protein Database [PDB] ID: 1EFA). The CAP activator (blue) can bind to the GTGA sequence of the *lac* operator when lactose is present, and glucose is absent (PDB ID: 1CGP). (c) We used CRISPR-Cas9 gene editing to create a library of *lac* operator site. We targeted a 9-nt segment of the operator site (21 bp) on the *E. coli* chromosome, resulting in 2.62×10^5^ *E. coli* genotypes that differ only in their *lac* operator sites. (d) We used these mutagenized cells to perform a selection experiment. In this experiment, we initially grew four replicate populations of all genotypes in a minimal environment containing glucose, and then split these populations to perform bulk competition of the populations in a minimal environment with either glycerol or lactose. (e) We subjected genomic DNA from the populations to high-throughput sequencing of the *lac* operator site both before and after selection, and estimated the fitness of each genotype from the resulting data. (f) Distribution of fitness relative to WT (zero, dashed black line) in glycerol (blue) or lactose (red), based on 187,485 genotypes in glycerol and 141,149 genotypes in lactose that met a read cutoff standard of 50 reads across all replicates (Methods). *[Created with BioRender.com]*

We focus on the *lac* operator because it plays a key role in regulating *lac* operon expression in response to different carbon sources. The *lac* operator is the primary binding site for the LacI repressor (Figure 1a-b). It also serves as a secondary binding site for the CAP activator (Figure 1a-b), although its role in CAP-mediated regulation is poorly understood (31,38). Because both repressors and activators may interact with the *lac* operator, mutations in this site can influence gene expression differently across environments. These mutations can affect how efficiently *E. coli* adapts physiologically to changing carbon sources.

To characterize the fitness landscape of the *lac* operator in *E.coli*, we used the Scarless CRISPR-Cas9 assisted recombineering (no-SCAR) system to systematically mutate nine nucleotide positions within the *lac* operator site, resulting in 2.62×10^5^ (≈0.25 million) operator site variants (39–41). We then performed bulk competition experiments in two different environments where glycerol and lactose are the only carbon sources, and estimated the fitness of these variants by high-throughput sequencing.

In glycerol the wild-type *lac* operon is repressed by LacI. However, in mutated operons with variants of the *lac* operator that have reduced affinity to LacI, the operon may be partially or fully expressed. The resulting expression cost may reduce fitness because the *lac* operon genes are not needed in a glycerol environment. Conversely, in lactose, detachment of the LacI repressor allows full expression of the *lac* operon, regardless of LacI’s affinity to *lac* operator site variants. This expression increases fitness, because *lac* operon genes are essential in lactose. However, the effect of lactose is made more complicated by the CAP, because in the absence of glucose, the CAP can bind to the *lac* operator site (39–41). Thus, the expression of the *lac* operon may also depend on CAP’s affinity for the *lac* operator site variants.

We found that the lactose operator is well-adapted to lactose environments but poorly adapted to glycerol environments, because many more operator variants have higher fitness than the wild-type in glycerol than in lactose. The fitness landscape is highly rugged in both environments and would only allow a small fraction of evolving populations to reach the highest fitness peaks. The landscape also shows topological differences that highlight how a change in a single carbon source can alter landscape topography substantially.

## Results

### Experimental design

We performed CRISPR-Cas9 deep mutagenesis to edit the *lac* operator site on the bacterial chromosome. Specifically, we mutated nine nucleotide positions within the operator site (Figure 1b), which resulted in a combinatorically complete library of 4^9^≈262,000 unique nucleotide sequences (*genotypes*, *variants*) (Figure 1c). Mutated positions had to be near a genomic PAM (protospacer adjacent motif) site, required for Cas9 recognition and mutagenesis (42). The nine mutated positions comprise sites with both high and intermediate importance for LacI binding (Figure S1), according to limited prior empirical information from genomic binding sites and other experimental data (42,43). Since no previous mutational data is available for the operator’s interaction with the CAP, we have no knowledge how variation at these sites may affect CAP binding.

After growing four replicates of the mutagenized library in glucose-supplemented M9 minimal medium (Figure 1d), we deep-sequenced genomic DNA to estimate pre-selection frequencies of wild-type (WT) and variant *lac* operator sites in our population. In glucose, the absence of cyclic AMP keeps the CAP activator inactive (Figure 1a). As a result, all *lac* operator variants are expected to convey similar (low) operon expression and thus similar fitness in this pre-selection environment. Additionally, in the absence of lactose, the LacI repressor also blocks transcription when binding to a functional operator.

We then performed bulk competition in two minimal environments with glycerol or lactose as sole carbon sources (Figure 1d). In the glycerol environment, cAMP accumulates due to the absence of glucose, activating CAP to promote *lac* operon expression (Figure 1a). However, transcription remains blocked if the LacI repressor is bound to the operator. Operator variants with reduced LacI binding may allow partial or full expression of the *lac* operon, which can reduce fitness since the operon is unnecessary in this environment.

In the lactose environment, the LacI repressor no longer binds DNA, which allows transcription of the *lac* operon (Figure 1a). All *lac* operator variants are thus expected to have similar fitness due to efficient lactose metabolism. However, fitness may also depend on CAP binding (Figure 1a). In the absence of glucose, the activated CAP can bind to the *lac* operator, and the strength of this binding may affect operon expression, leading to different fitness outcomes in lactose (Figure 1a).

After growing four replicates of the library for 16 hours in each of the two environments, we isolated DNA and sequenced more than 15 million reads of *lac* operator DNA per replicates to determine the final frequencies of WT and variant *lac* operators in each environment (Figure 1e). Variant frequencies were highly correlated across all replicates, with average Pearson’s pairwise correlation coefficient (r) of 0.971 in glucose, r=0.945 in glycerol, and r=0.825 in lactose (Figure S2-S4). We retained only those genotypes with sufficient read counts across replicates, and those present in before- and after-selection replicates for further analysis (see Methods).

We used variant frequencies across all replicates to calculate the fitness of each *lac* operator variant relative to the wild-type (WT) (Figure 1f, Methods). We also fitted a generalized linear model (GLM) to the data, which allowed us to estimate fitness measurement errors from read count variation across replicates. After filtering for sufficient read counts (Methods), we obtained fitness data for 71.52% (187,485/262,144) of all possible variants in glycerol and 53.84% (141,149/262,144) in lactose.

As prescribed by population genetic theory, we represent all fitness values on a natural logarithmic scale relative to the WT (44). On this scale, the WT has a fitness value of 0, and a variant *i* with a relative fitness (*f*_i_ = *r*_i_ - *r_WT_*) of 1 has an exponential growth rate that is e≈2.718 times greater. Approximately 25.4% of the *lac* operator variants (47,558/187,485) were fitter than the WT in glycerol, suggesting that these variants bind the LacI repressor more strongly. In contrast, only 0.45% of variants (630/141,149) showed higher fitness than the WT in lactose. If fitness in lactose depended solely only on LacI binding, all variants would show similar fitness and thus show a much narrower distribution than in glycerol. However, that is not the case (Figure 1f). The distribution of fitness is actually wider in lactose than in glycerol (Lactose: Mean Absolute Deviation [MAD]: 0.320; Glycerol: MAD: 0.256). Levene’s test (45) rejects the null hypothesis that fitness variances in the two environments are equal (Levene’s Statistic: 6052.72, P < 2×10^-308^). These observations show that fitness is not just influenced by LacI binding but also by other factors, such as CAP binding to the operator.

To validate our fitness estimates with a different method, we isolated 83 *lac* operator variants and measured their growth rates in glycerol and lactose in single cultures (Methods). We found that relative fitness from bulk competition strongly correlates with single-culture growth rates (Figure S5-S13; glycerol: Pearson’s r= 0.88, P= 4.11×10^-28^; lactose: Pearson’s r= 0.95, P= 2.00×10^-42^, N= 84).

### Four major categories of *lac* operator variant fitness

We next compared the fitness of each variant in the glycerol and the lactose environment (Figure 2a). Variants with high fitness in one environment tend to have high fitness in the other (Pearson’s r= 0.473, P<2.0×10^-308^, Spearman’s ρ= 0.433, P<2.0×10^-308^, N= 133888; *H_O_:r* = 0 for Pearson, *H_O_*: ρ =0 for Spearman). In other words, no trade-off exists between fitness in the two environments. This positive relationship is also supported by individual growth assays (Pearson’s r= 0.75, *P*= 2.27×10⁻¹², *N*= 61; Figure S14).

**Figure 2.**
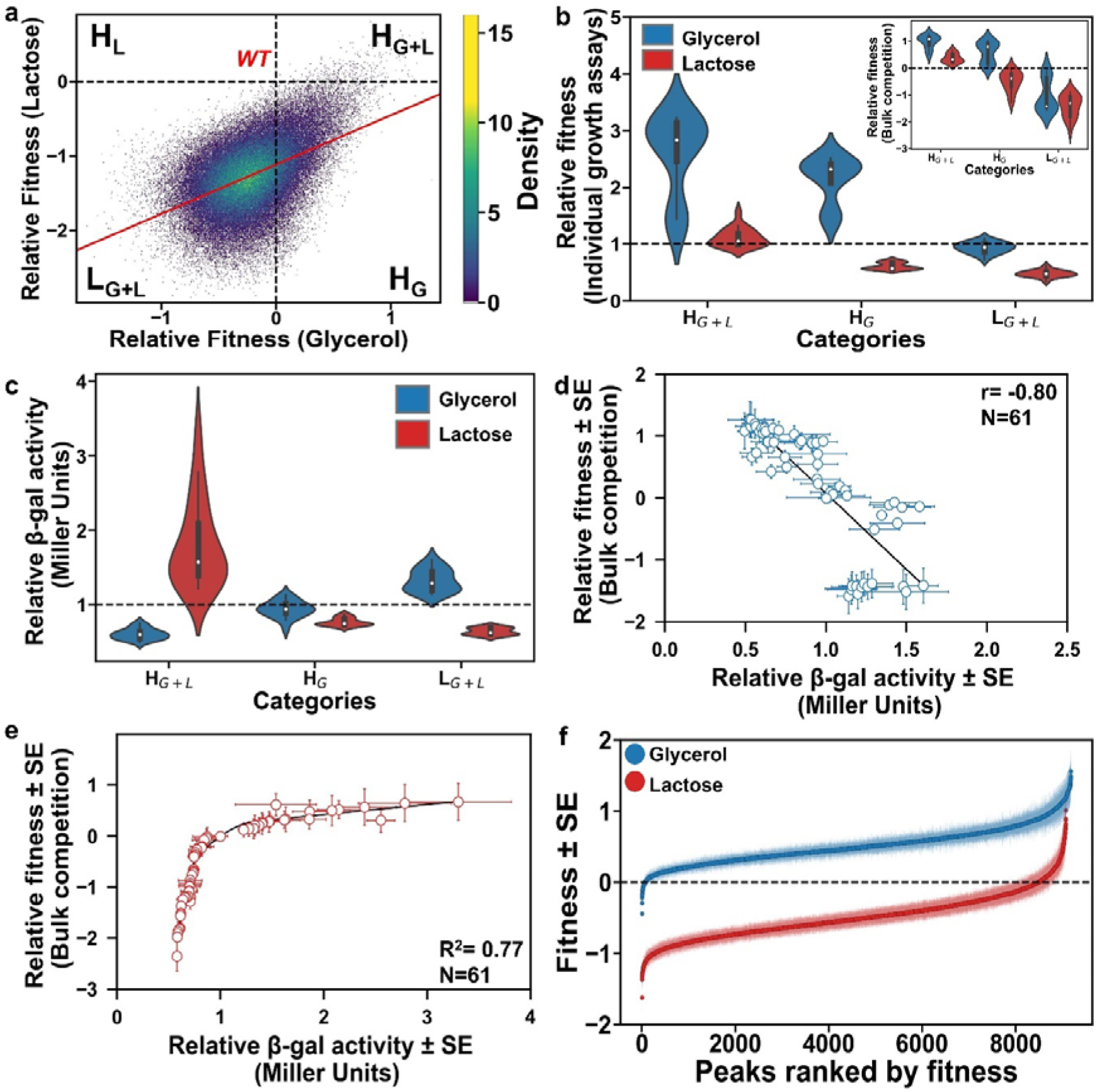
Relative fitness and enzyme activity of *lac* operator variants, and the distribution of fitness peaks in the *lac* operator landscape across environments. (a) Density scatter plot of relative fitness (WT fitness: 0, dashed lines) of all *lac* operator variants in lactose and glycerol. Color (see legend) indicates number of data points. Regression line (red) is based on a Gaussian GLM model fitted to the data. Pearson’s r=0.473, P<2.0×10^-308^, Spearman’s ρ=0.433, P<2.0×10^-308^, n= 133888. Variants are categorized into four groups, H_G+L_ (high fitness in glycerol and lactose), H_G_ (high fitness in glycerol), H_L_ (high fitness in lactose) and L_G+L_ (low fitness in glycerol and lactose). (b) Relative fitness from individual growth assays of 20 selected variants each from three categories (H_G+L_, H_G_ and L_G+L_) in glycerol and lactose. (dashed line: WT fitness in individual growth assays). The inset shows the relative fitness of the same variants measured from bulk competition experiments in glycerol and lactose. (dashed line: WT fitness from bulk competition). (c) Relative β-galactosidase activity of 20 selected variants in glycerol and lactose from the same three categories as (b). (Dashed line: WT activity). (d) Relative β-gal enzyme activity (horizontal axis) and its relationship to relative fitness estimated from bulk competition (vertical axis) in glycerol. (e) like (d), but in lactose. (f) Ranked distribution of fitness and corresponding standard errors for ∼9000 peak genotypes in the *lac* operator landscape in either glycerol (blue) or lactose (red). (Dashed black line: WT fitness).

The variants fall into four main categories (Figure 2a, S15). The first category (H_G+L_) comprises those 0.24% of variants (321/133888) with higher fitness than WT in both environments. These variants likely repress the operon more strongly than WT in glycerol, where expression is unnecessary, and activate (de-repress) the operon more strongly than WT in lactose, where expression improves fitness. The second category (H_G_) comprises 19.8% (26567/133888) of variants with high fitness in glycerol but low fitness in lactose. These variants likely repress the operon strongly in glycerol, increasing fitness, but activate it weakly in lactose, lowering fitness. The third category (H_L_) consists of only two variants (0.0015%). They have low fitness in glycerol but high fitness in lactose, possibly because they repress the operon less strongly in glycerol while activating it strongly in lactose. The fourth category (L_G+L_) comprises 79% of all variants (106998/133888). They have lower fitness than the WT in both environments. The most likely reason is that they repress the operon less strongly in glycerol and activate it less strongly in lactose.

Categories H_G+L_, H_G_, and L_G+L_ comprise enough variants to test this hypothesis by exploring how mutations in the *lac* operator site influence gene expression, enzyme activity, and thus fitness in glycerol and lactose. To this end we generated three small oligopool libraries, each consisting of 50 *lac* operator site variants from category H_G+L_, H_G_, and L_G+L_ using CRISPR-Cas9 mutagenesis. We then selected 20 variants from each category and measured their growth rates (Figure 2b, S7-S12). We also quantified their β-galactosidase production, as a measure of operon (*lacZ*) expression and its relationship to fitness (Figure 2c-e, S16-S23). The wild-type’s growth rate equals 0.542±0.127 h^-1^ in glycerol (mean±SD), and 2.048±1.087 in lactose h^-1^ (mean±SD, Figures S5-S6), corresponding to a doubling time of 8.48 h and 7.31 h, respectively. The wild-type’s β-galactosidase activity equals 712.0±32.7 (mean±SD) Miller units in glycerol and 2006.6±138.0 (mean±SD) Miller units in lactose (Figure S16). We report measurements for other variants as the ratio of their β-galactosidase activity and growth rate relative to the WT, both set to one.

Fitness measurements from bulk competition agree with individual growth assays (Figure S13, S17-22). β-galactosidase measurements show that H_G+L_ variants repress the *lac* operon strongly in glycerol and activate it strongly in lactose, leading to higher fitness in both environments (Figure S7-S8, S17-S18). H_G_ variants, while efficient at repressing the *lac* operon in glycerol, are less effective at activating the operon in lactose, which corresponds to their lower fitness in lactose (Figure S9-S10, S19-S20). Conversely, L_G+L_ variants show poor repression in glycerol and impaired activation in lactose, explaining their low fitness in both environments (Figure S11-S12, S21-S22). (See Supplementary Results 1 for a more detailed analysis.)

Further supporting this analysis, we found that in glycerol high fitness variants express less β-galactosidase (Figure 2d; bulk competition: r= −0.80, P= 1.19×10^-14^, N=61; Figure S23a; individual assays: r= −0.87, P= 4.37×10^-20^, N=61), just as expected if β-galactosidase incurs a cost but no benefit in glycerol. Conversely, in lactose high fitness variants express more β-galactosidase (Figure 2e; bulk competition: R^2^=0.774, F-statistic = 47.87, F-test P= 4.01×10^−9^, N=61; Figure S23b; individual assays: R^2^=0.935, F-statistic = 67.00, F-test P= 2.99×10^−11^, N=61). We also note that the relationship between activity and fitness is highly non-linear in lactose. When the *lac* operon is already highly expressed, increasing its expression further provides less and less benefit, likely because further increases in β-galactosidase activity do not proportionally enhance metabolic flux (46). Similar nonlinear expression-fitness relationships have been observed in other systems (6,19,47–49).

### The *lac* operator fitness landscape is rugged in both glycerol and lactose

To study the fitness landscape of the *lac* operator site, we represented our data as a graph, where each node represents an operator variant and its associated fitness. Edges connect nodes that differ by one nucleotide. In both glycerol and lactose, 99.9% of all variants belong to the largest connected subgraph, also known as the giant component (50,51). We focus our analysis on this subgraph.

The topology of a fitness landscape affects whether a population evolving by natural selection can reach its highest fitness peaks (11,52). A smooth landscape with one or few peaks facilitates such adaptive evolution, while a rugged landscape with many peaks hinders evolution (52). To assess ruggedness, we counted the number of peaks, which are variants whose fitness is higher than all their one-mutant neighbors (41,53–55).

The *lac* operator landscape is highly rugged in both environments. It has 9,183 peaks (4.9% of all genotypes) in glycerol, while it has 9,074 peaks (6.4% of all genotypes) in lactose (Figure 2f). Notably, in glycerol, most peaks (99.25 percent) show higher fitness than the WT (with fitness 0). Among high fitness variants (f_l_ > 0) in glycerol, approximately 19.2% (9114/ 47,558) are peaks. In contrast, in lactose most peaks (93.41 percent) have lower fitness than the WT, and 95% of the high fitness (f_l_ > 0) variants in lactose (598/630) are peaks. A likely reason for these differences between the two environments is that the WT *lac* operator is better adapted to lactose environments, which may have been more frequent than glycerol environments in *E.coli’s* evolutionary history.

The glycerol and lactose landscapes share only a minority of peaks (1423, 15% and 16% of all peaks in glycerol and lactose, respectively). Shared peaks with high fitness in one environment are likely to have high fitness in the other (Pearson’s r = 0.620, P=1.022×10^-151^, Spearman’s ρ = 0.559, P=1.577×10^-117^, n=1423, Figure S24).

Since the spatial arrangement of peaks in a landscape can influence their evolutionary accessibility, i.e., the number of evolutionary paths along which every single mutational step increases fitness, we also examined how close the peaks are to one another. To this end, we calculated the genetic distance between each pair of all peaks, i.e., the minimum number of single-nucleotide changes required to reach one peak from another (regardless of how each change affects fitness). We compared this distance distribution with the same distribution but for an equal number of randomly selected variants (Figure S25). The mean distances differ by less than 1% in both glycerol (d = 6.75 for peaks and d = 6.74 for random variants) and lactose (d = 6.75 for peaks vs. d = 6.73 for random variants). The overall shapes of the distributions are highly similar (Figure S25).

The same holds for the 100 peaks with highest fitness – the top 100 peaks (fitness>1.207 in glycerol and fitness>0.389 in lactose). That is, the distance distributions of the top 100 peaks and random variants are highly congruent, and average peak distances differ by less than one percent in both glycerol (Figure 3a, d = 6.73 for top 100 peaks and d = 6.77 for random variants) and lactose (Figure 3b, d = 6.75 for top 100 peaks vs. d = 6.71 for random variants). In summary, fitness peaks in general and the top 100 peaks are not clustered but scattered throughout the landscape in both environments (Figure S26-27). Figure S28 indicates which nucleotides occur preferably at high fitness sequences (Supplementary Results 2).

**Figure 3.**
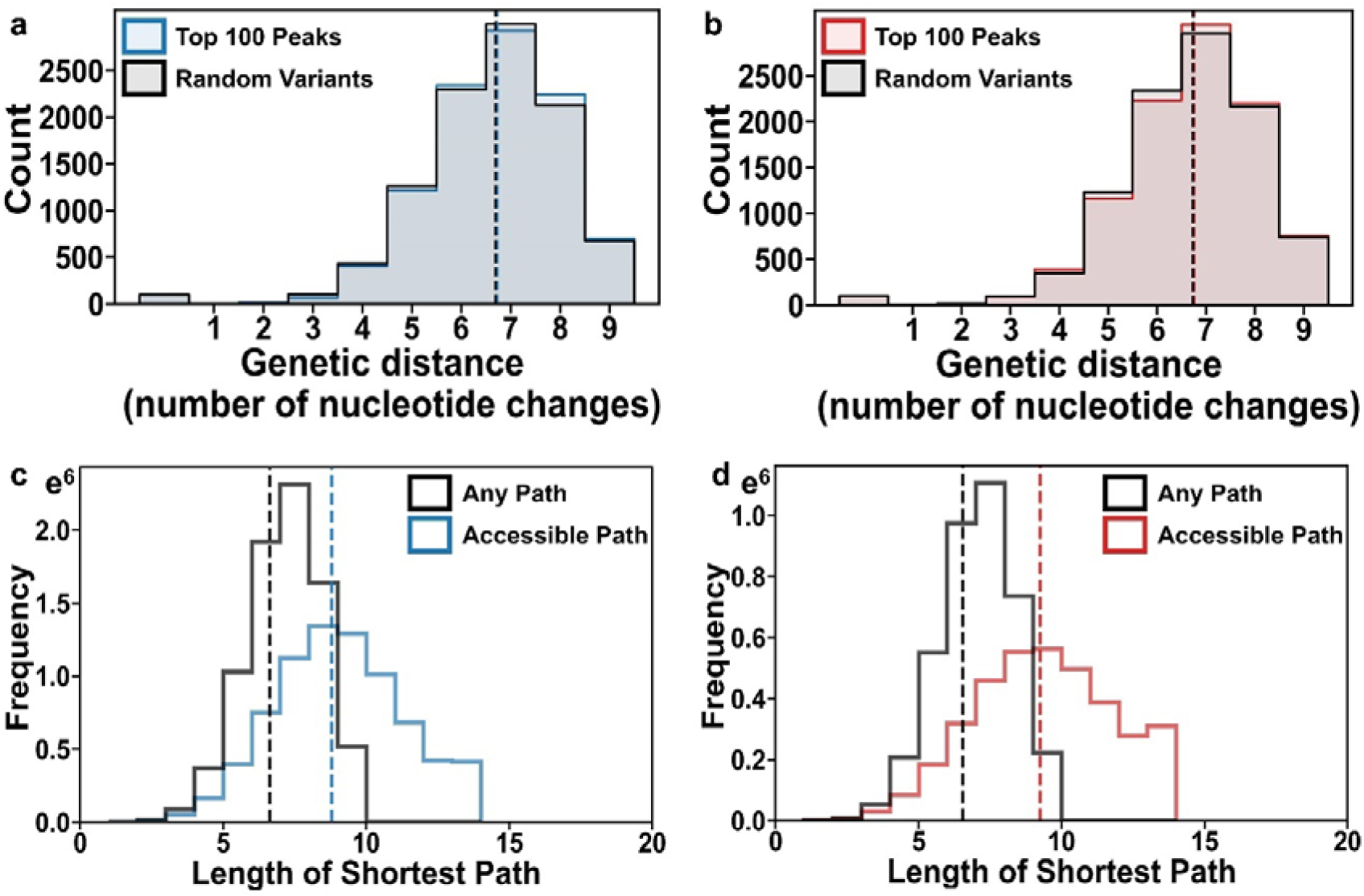
Genetic distance of fitness peaks and their accessibility in *lac* operator landscape across different environmental conditions. (a) Genetic distance distribution of the top 100 peaks (blue) in glycerol and 100 randomly selected non-peak variants (black), with mean values marked by vertical lines (d=6.73 for peaks [in blue] and d=6.77 for the random variants [in black]). (b) like (a), but for peaks in lactose (d=6.75 for peaks [red] and d=6.71 for random variants [black]). (c-d) Accessible paths to the top 100 peaks are longer than the shortest distance in genotype space. (c) The black histogram shows the distribution of genetic distances for all pairs of variants and the peaks accessible from them, while the blue histogram depicts the number of mutational steps on the shortest accessible paths between variants and the peaks attainable from them in glycerol. (d) like (c) but for lactose.

### The top 100 peaks are moderately accessible in either environment

To investigate whether the ruggedness of our landscapes may hinder adaptive evolution, we first examined the evolutionary accessibility of all peaks. We began by assessing whether accessible paths exist from each non-peak variant to each peak, i.e., paths along which every single mutational path increases fitness. Specifically, we quantified the size of each peak’s basin of attraction, that is, the total number of variants from which a peak is accessible (Figure S29a-b).

We found that the basin sizes of peaks vary widely. In glycerol, it ranges from 0.02% to 58.2% (32 and 103,752/178,301) of all variants. In lactose, it ranges from a single variant to 50.3% (66,393/132,073) of variants (Figure S29a-b). The average basin of attraction comprises 39.80±9.74% (mean±SD) percent of variants in glycerol and 22.77±10.96% percent of variants in lactose, a difference that is statistically significant (Mann-Whitney U statistic: 9,621,941.000, P<2.0×10^-308^, Figure S29c).

Notably, high fitness peaks tend to have larger basins of attraction than intermediate or low fitness peaks (Figure S30-S31) both in glycerol (Spearman’s ρ = 0.41, P < 2×10^-308^, N=9183; Figure S31a) and in lactose (Spearman’s ρ = 0.46, P < 2×10^-308^, N=9074; Figure S31b). Supplementary Results 3 provides more detailed analysis.

While we have analyzed the accessibility of all peaks in the landscape, from here on we will focus on the top 100 peaks, which are most important for understanding how ruggedness affects adaptive evolution. In a smooth landscape, the length of the shortest accessible path between each non-peak variant and each peak equals the genetic distance (number of nucleotide changes) between the two. In a rugged landscape like ours, accessible paths may be longer. Indeed, the shortest accessible path between a variant and one of the top 100 peaks in glycerol, is on average more than two mutational steps longer (mean±SD = 8.79±2.59 mutations) than the genetic distance (6.64±1.33 mutations; Mann-Whitney Test, U = 48026880456473.5, P< 2×10^-308^; Welch’s t-test, t=2079.24, P< 2×10^-308^; N = 7910243; Figure 3c). The same holds in lactose (mean±SD = 9.25±2.92 mutations in an accessible path vs. a genetic distance of 6.54±1.35 mutations; Mann-Whitney Test, U = 11981608114124.5, P < 2×10^-308^; Welch’s t-test, t=1662.23, P< 2×10^-308^; N = 3870382; Figure 3d). Therefore, in both glycerol and lactose environments, accessible paths to the top 100 fitness peaks are more circuitous than the shortest distance in genotype space. A population evolving along such a path is thus also more likely to be diverted from such a path, and not reach a high fitness peak.

Given that the top 100 peaks have large basins of attraction, we hypothesized that the basins of different peaks may share genotypes. If so, adaptive evolution starting from one genotype might be able to reach either peak. In other words, genotype sharing between basins of attraction may lead to evolutionary contingency — the dependence of a historical process on chance events. To quantify the potential for such contingency, we first computed the overlap between basins of attraction, the fraction of genotypes they share (Figure S32). For computational feasibility, we focused this analysis on all pairs of the top 100 peaks (Figure S32a-b).

This overlap between basins is substantial. On average, the basins of the top 100 peaks in glycerol share 73.8% ± 12.2% (67347.2 ± 13820.6 variants, mean±SD, N=4950) of genotypes, ranging from 13.33% (10548) to 88.36% (93779). In lactose, the average overlap is lower at 57.1% ± 14.4% of variants (28429.1 ± 9732.0 variants, mean± SD, N=4950) ranging from 10.17% (2199 variants) to 87.76% (59176 variants). The overlap is significantly larger in glycerol (Figure S33, two-sided Mann–Whitney UO= 2,925,859; P<O2.0O×O10^−308^; NO=O4950). In other words, any one non-peak genotype is more likely to be able to access more than one peak in the glycerol environment. Analogous patterns hold for peaks of internediate and low fitness (Figure S32c-f).

Our next analysis underscores that the two landscapes differ in their potential for contingency. In this analysis we quantified the number of the top 100 peaks that are accessible from each non-peak variant. In glycerol, 138,553 variants (77.7%) have access to one or more of these peaks (Figure 4a), and 10,191 variants (5.7%), can access all 100 top peaks. In contrast, peak accessibility is more limited in lactose (Figure 4b), where 94,292 variants (71.4%) have access to at least one top 100 fitness peak, and only 1,178 (0.89%) can access all 100 peaks. In sum, starting from any one non-peak variant an evolving population can typically access multiple top 100 peaks, and this number of accessible peaks is greater in the glycerol environment.

**Figure 4.**
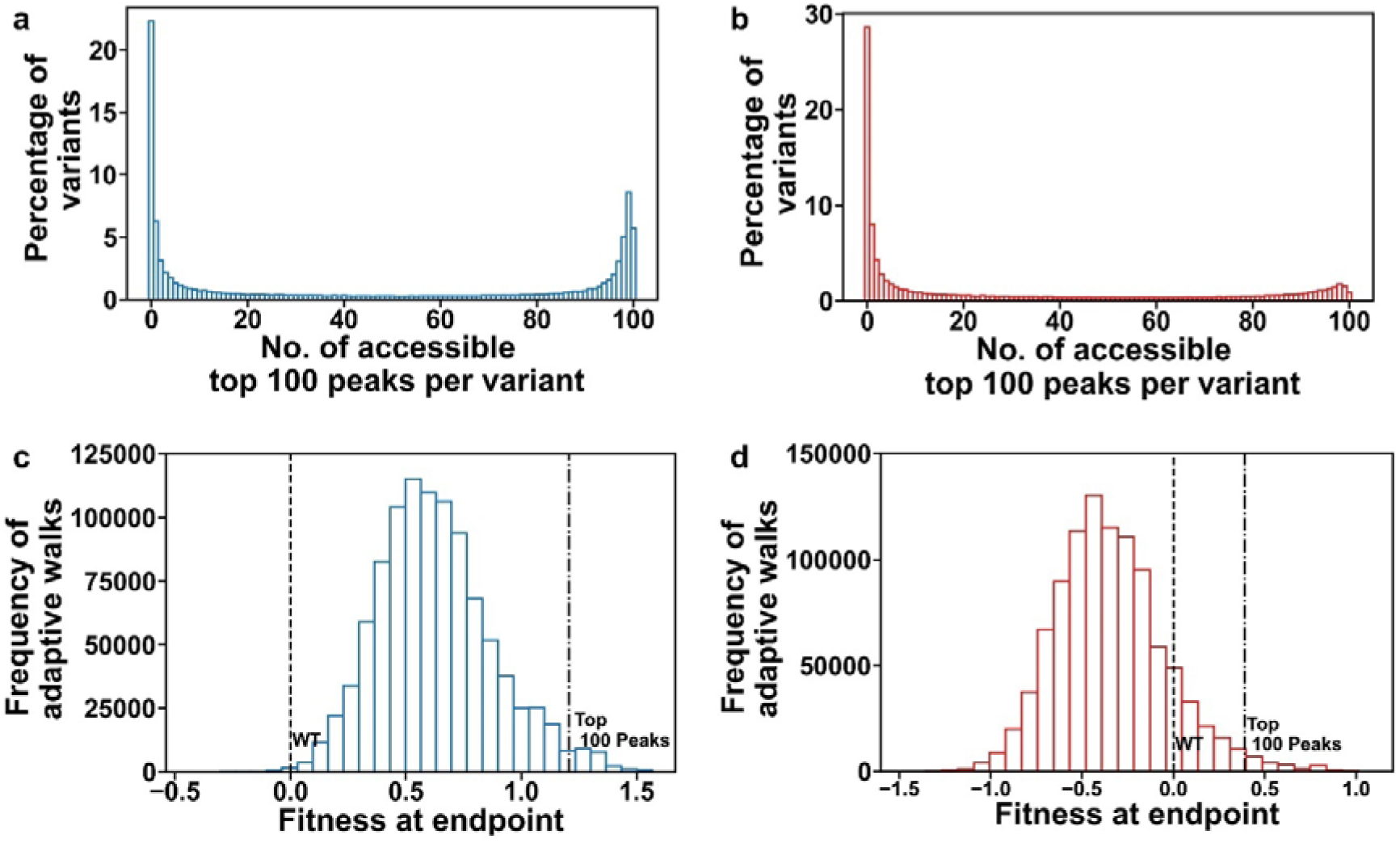
The top 100 peaks are moderately accessible in both environments. (a-b) Multiple top 100 peaks are accessible from many non-peak *lac* operator variants. The panels show the distribution of the number of the top 100 peaks that are accessible from each non-peak variant in glycerol (a) and in lactose (b). (c) Frequency distribution of fitness values attained by 10^3^ adaptive walks starting from each of the 10^3^ lowest-fitness non-peak variants, i.e,10^6^ adaptive walks in total (Methods) in glycerol. The left dashed vertical line at x=0 indicates the fitness of the WT in glycerol. 99.85% of adaptive walks achieved a fitness value higher than the WT. The right dotted vertical line marks the fitness of the lowest of the top 100 peaks. Only 2.32% of adaptive walks reached one of these peaks. (d) Like (c) but in lactose. 12.27% of adaptive walks reach a fitness higher than the WT, and 2.04% of adaptive walks reach one of the top 100 peaks.

### Few evolving populations reach a top 100 fitness peak

Thus far, we have only analyzed the static topography of each landscape, which neglects that adaptive evolution on a landscape is a dynamic process. For example, even though a peak may be accessible by a fitness-increasing path, if accessible paths are rare among all paths, an evolving population may be unlikely to reach the peak. To study evolutionary dynamics, we simulated adaptive evolution in the weak-mutation strong-selection regime (SSWM, Methods) (56–58) which is justfied by the low mutation rate at our nine-nucleotide mutational target in the *E. coli* genome (9×2.2×10^−10^ = 1.98×10^−9^ mutations per binding site per generation) (59). In this regime, most beneficial mutations go to fixation before subsequent mutations arise that will go to fixation, such that adaptive evolution effectively becomes an adaptive walk (Methods).

Specifically, we performed 1,000 adaptive walks from each of the 1,000 *lac* operator variants with lowest fitness, i.e., a total of 10^6^ such walks. At each step of such a walk, we introduced a random DNA mutation, and allowed it go to fixation with a _1-e_-2Nsp probability F_l_, given by the well-established expression 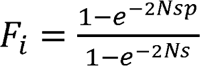 derived by Kimura (44,60,61). This expression depends on both the fitness change s caused by the mutation (the selection coefficient), and on the (effective) population size N (see Methods). Given that *E. co*li populations are very large (>10^8^ individuals) (62), genetic drift is weak and most fixation events will be driven by selection. We continued each walk until it reaches a fitness peak.

In glycerol, 99.85% of adaptive walks reached peaks with fitness higher than the WT (f_l_ > 0), but only 2.32% reached one of the top 100 peaks (Figure 4c). In lactose, only 12.27% of adaptive walks reached peaks with fitness higher than WT. However, the fraction of adaptive walks reaching the top 100 peaks, 2.04%, was similar to that in glycerol (Figure 4d). That is, a similar fraction of populations reaches a top 100 peak in both environments, even though these peaks are accessible from more variants in glycerol (77.7%) than in lactose (71.4%). These evolutionary dynamics analysis highlights the limitations of adaptive evolution on a rugged landscsape.

To better understand the importance of contingency, our next analysis focused only on those random walks that did reach one of the top 100 peaks. Among starting genotypes (n = 938 in glycerol; n=904 in lactose) that accessed at least one top 100 peak (23,175 walks in glycerol and 20,441 walks in lactose), most reached multiple peaks. Specifically, in glycerol, adaptive walks starting from 79% of starting variants reached two or more top 100 peaks, whereas walks starting from only 15.3% and 5.7% starting variants reached exactly one or no top 100 peak, respectively. In lactose, the corresponding values are 71.4%, 19.4%, and 9.2% respectively. On average, adaptive walks that originated from any one variant reached 3.21 ± 1.69 peaks in glycerol (median = 3.0), and 2.72 ± 1.46 peaks in lactose (median = 2.0). While multiple top 100 peaks were accessible from individual genotypes in both environments, we observed no starting genotype from which adaptive walks could reach all top 100 peaks (Figures 5a, 5b).

**Figure 5.**
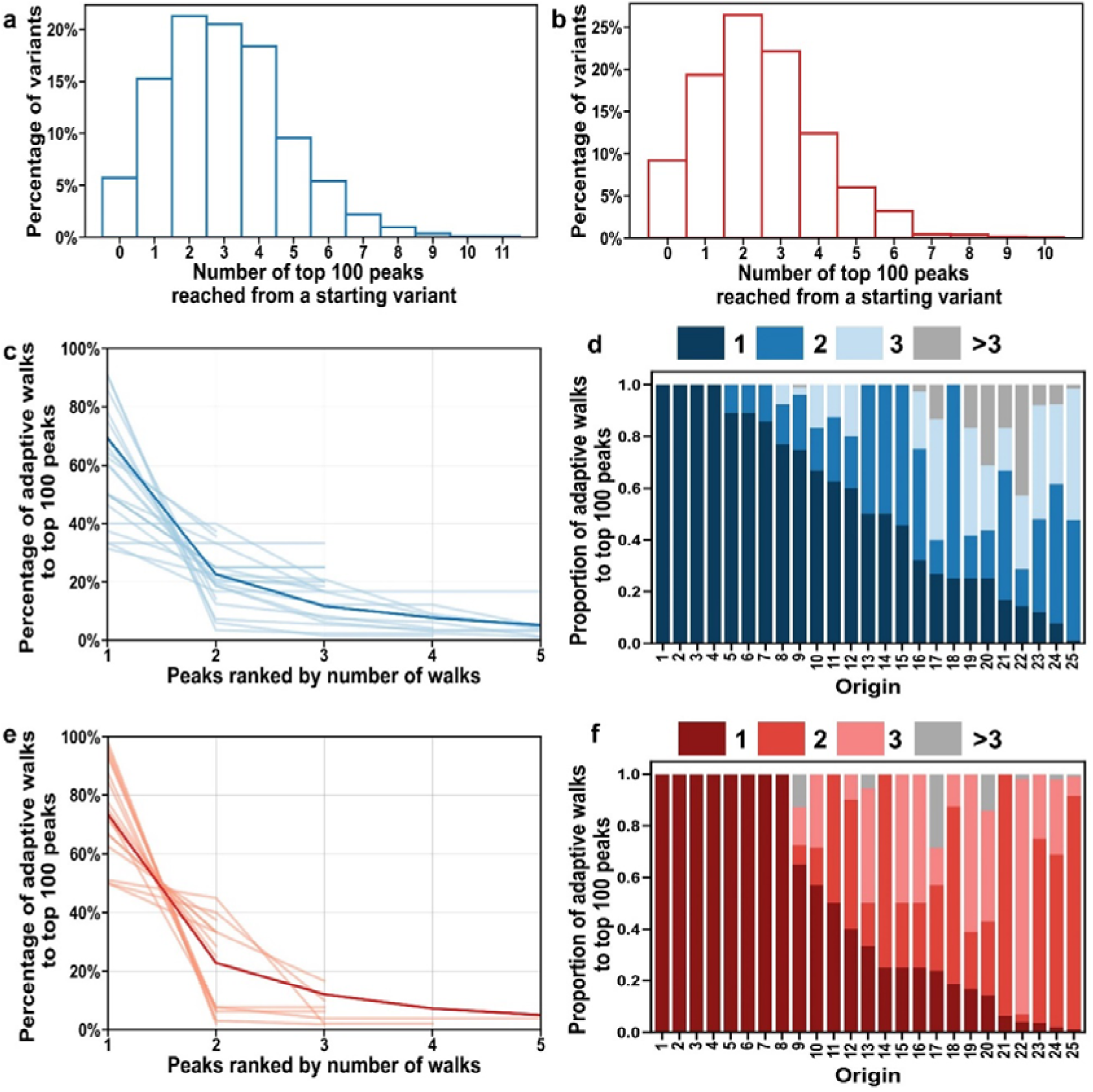
Some top 100 peaks are reached more frequently by adaptive walks than others. (a-b) More than one top 100 peak is reached by adaptive walks starting from most low fitness genotypes. The histogram shows the proportion of low fitness genotypes (vertical axes) from which zero, one, or more than one top 100 peaks are reached (horizontal axes) by adaptive walks starting from these genotypes. Data is based on 10^3^ adaptive walks from each of the 1000 lowest fitness non-peak variants (Methods). In (a) glycerol, 5.7%, 15.3%, and 79% of variants, and in (b) lactose, 9.2%, 19.4%, and 71.4% of variants, accessed no peaks (x = 0), exactly one peak (x = 1), or multiple peaks (x > 1), respectively. (c-f) Adaptive walks tend access some top 100 peaks more frequently than others. (c) The vertical axis represents the percentage of adaptive walks that accessed the first, 2^nd^, 3^rd^, or 4^th^ most frequently attained peak (horizontal axis) among all walks that reached one of the top 100 peaks (starting from 938 starting variants). Each light blue line summarizes data from adaptive walks initiated from the same variant. To minimize visual clutter, data from only 25 randomly selected starting variants are shown in light blue. The dark blue bold line represents the mean percentage across all 938 starting variants. (d) Data from the same 25 randomly selected starting variants as in (c) are represented as bars, with stacks indicating the number of top 100 peaks accessed during 10^3^ adaptive walks. The color legend indicates the total number of peaks accessed. Stack heights represent the proportion of walks (from those reaching a high fitness peak) that accessed each peak. Stacks are arranged by frequency, with the three most frequently accessed top 100 peaks color-coded in different shades of blue, while other top 100 peaks are shown in grey. (e, f), like (c, d) but for the lactose environment and for 904 starting variants from which a high peak is reached.

The likelihood of accessing a given peak varied widely. In glycerol, on average, 69.3 ± 22.4% of adaptive walks (n=16,813) that reached a top 100 peak converged on the most frequently attained one (Figure 5c-d). Similarly, in lactose, 73.4O±O22.0% of adaptive walks (nO=O16,137) that reached a top 100 peak accessed the most frequently attained one (Figure 5e-f). In summary, our observations suggest that the identity of any peak attained during adaptive evolution is highly contingent on stochastic events, with some peaks being more frequently accessed than others.

## Discussion

We used bulk competition and high-throughput sequencing to map and study the fitness landscape of ∼250,000 *lac* operator variants in both glycerol and lactose. We deliberately excluded a glucose environment, because catabolite repression through glucose is unrelated to *lac* operator binding and is thus not relevant to study operator mutations. We also validated our high-throughput fitness measurements via independent growth rate measurements and beta-galactosidase assays in glycerol and lactose (Figures 2b-e, Figure S13). We chose these environments because they convey different regulatory states of the *lac* operon. In glycerol, where *lac* operon expression has costs but no benefits, operator variants with weak affinity to the LacI repressor entail high *lac* operon expression and thus low fitness (Figure 2). In lactose, where *lac* operon expression is beneficial, the repressor LacI should detach from the *lac* operator, such that all our operator variants should have equal fitness.

Our fitness data belies this simple textbook model of *lac* operon regulation. The reason is that *lac* operator mutants do not have identical fitness in lactose. To the contrary, the distribution of fitness in lactose is even broader than this distribution in glycerol. A likely explanation is the demonstrated ability of the *lac* operator to act as a secondary binding site for the CAP/CRP activator (31,38,63–66). Previous studies suggest that CAP binding at the *lac* operator might inhibit promoter function, with little to no effect on CAP-mediated activation, but the CAP-operator interaction may influence fitness in ways not captured by these studies (64–66). Unfortunately, directly studying the role of CAP alone is challenging, as its secondary binding site completely overlaps with the LacI repressor binding site. Dissecting these interactions remains an exciting direction for future work.

Our analysis of landscape topography shows that both landscapes are highly rugged, with ∼9,000 fitness peaks in each environment. An analysis of statistical power to identify peaks shows that these numbers are not artefacts of measurement error (Figure S34-35). Fitness peaks, including the top 100 peaks, are scattered through genotype space, with a distance distribution similar to randomly chosen sequences. Only a minority of evolving populations (approximately 2%) reach the top 100 peaks in both environments, which highlights that landscape ruggedness can be a substantial obstacle for adaptive evolution.

The ruggedness of our landscapes is also reflected by a high frequency of reciprocal sign epistasis, a form of non-additive interaction in which a combination of two individual mutations is beneficial, even though each individual mutation is deleterious (Figure S36a, Table S8). Specifically, 33% of mutant pairs in both glycerol and lactose exhibited this type of epistasis (Table S8). This high incidence of epistasis is consistent with previous work on bacterial TFBSs (67–69), where epistasis is generally more prevalent than in eukaryotic TFBSs (4-5%) (11). We also observed diminishing returns epistasis in both landscapes (70,71), a form of global epistasis where the fitness benefit of a mutation decreases as genotypes approach high fitness (Figure S36b-c).

A notable difference between the two landscapes exists in the evolutionary accessibility of peaks, i.e., the existence of paths that lead from a specific non-peak genotype to a peak, such that no step decreases fitness. Each peak has a basin of attraction, i.e., a subset of genotypes from which accessible paths to the peak exist, In glycerol, the average basin size of the top 100 peaks is 74.8 percent larger than in lactose, meaning that its peaks are more accessible (Figure S29c).

In addition, the basins of attraction for the top 100 peaks share 29.3 percent more variants in glycerol than in lactose, highlighting greater potential for evolutionary contingency-the influence of chance events for the outcome of adaptive evolution. To quantify this potential, one can enumerate the number of peaks that adaptive evolution starting from the same genotype can reach. Indeed, among those evolving populations that originated from the same non-peak genotype and reached at least one top 100 peak, 10.64% reached more such peaks in glycerol than in lactose.

In both environments, some top 100 peaks are preferentially reached by adaptive walks, and this preference is weaker in glycerol. For example, the five most frequently reached top 100 peaks are accessed by 5.43% and 7.61% of adaptive walks in glycerol and lactose, respectively (Figure 5).

The WT operator is suboptimal, because 0.24% of operator variants outperform it in both environments. It is also not a fitness peak, because one of its 1-mutant neighbors shows higher fitness in lactose, and three do so in glycerol. Not surprisingly, lactose must have exerted greater selection pressure than glycerol on the *E. coli* genome during *E. coli’s* natural history, as the WT *lac* operator is much better adapted to lactose than to glycerol. Specifically, among the *lac* operator variants with fitness data in both environments, merely 0.45% show higher fitness than the WT in lactose, compared to 25.4% in glycerol (Figure 1f). Even this moderately high WT fitness in glycerol may overestimate the evolutionary importance of glycerol environments, because fitness correlates positively across environments (Figure 2a). Thus, selection for high fitness in lactose could entail moderately high fitness in glycerol, even if selection had never acted on fitness in glycerol. Also relevant here is that lactose is an ecologically important sugar for *E. coli* (72,73), and previous studies show that *E. coli* has evolved to regulate the *lac* operon in response to lactose (18,74–79). In contrast, glycerol is typically present at much lower concentrations in *E. coli*’s natural habitats, such as the mammalian gut, where it is not a major or stable carbon source, and is mainly produced by microbial lipid degradation (80,81). Although *E. coli* was probably exposed to glycerol in its evolutionary history, because it can grow on glycerol as a sole carbon source, it does so slowly (82–84). Experimental evolution can achieve faster growth on glycerol, but requires extensive genetic and metabolic reorganization, including specific mutations (e.g., in *glpK* and *rpoC*) that enhance glycerol metabolism (85–88). These observations underscore that glycerol utilization by *E. coli* is not a naturally optimized trait that is ecologically as important as lactose utilization.

One limitation of experimental landscape studies like ours is that all fitness measurements are associated with measurement errors. We reduced such errors by requiring multiple sequence reads for individual operator variants and quantified errors by replicating our measurements four-fold (Supplementary Methods 8). Our statistical power analysis shows that these errors are low enough to allow confident characterization of the landscape. For example, we can expect the vast majority of high fitness peaks to be identified correctly (Supplementary Methods 8). Specifically, 100 and 89 percent of our top 100 peaks are true positive peaks in glycerol and lactose, respectively. In addition, 99.9 and 86.8 percent of our peaks with fitness above the wild-type are expected to be true positives in glycerol and lactose.

A second limitation results from our focus on fitness measurements rather than biophysical properties, such as the affinity of *lac* operator variants for the LacI repressor. To identify the biophysical causes for the fitness differences we observed remains an important task for future work. Thirdly, we studied only two environments, but the *lac* operon evolved in multiple environments. Future work will reveal whether expression costs in additional environments can also influence the *lac* operator landscape.

Overall, our study shows that even short regulatory sequences like the *lac* operator can have very rugged fitness landscapes that limit the paths available to evolution. In addition, an environmental change in a single carbon source can change the topography of such a landscape. And the WT *lac* operator, albeit well-adapted to a lactose environment, is not the best possible operator. Our work provides a starting point for future studies to better understand how regulatory sequences evolve.

## Materials and Methods

### Strains and plasmids

We used *E. coli* strain MR^S^, a derivative of K-12 MG1655 with a wild-type mutation rate that carries the amber stop codon suppressor mutation *glnV44* (*supE44*) (89). We edited the *E. coli* genome using the pKDsgRNA-lacO and pCas9-CR4 plasmids from Addgene, with inducible Cas9 expression and λ-Red recombination (Supplementary Methods 1) (39,40). A complete list of strains and vectors is provided in Tables S1 and S2.

### Gene editing and bulk competition experiments

We grew *E. coli* MRS/pCas9-CR4/pKD-gRNA-lacO in SOB medium with antibiotics and 1 mM Isopropyl β-D-1-thiogalactopyranoside (IPTG). We induced lambda-Red expression with 0.2% arabinose for one hour, prepared electrocompetent cells and transformed them with LacI-Regular-Oligo (Supplementary Table S3) as the recombination template for *lac* operator gene editing. We recovered the cells in SOC, plated them on selective agar with IPTG and anhydrotetracycline, and incubated the plates at 30°C. We performed mutagenesis in the presence of Isopropyl β-D-1-thiogalactopyranoside (IPTG) to prevent LacI from interfering with Cas9 or recombination.

To remove residual IPTG from mutagenized *E. coli* cells before bulk competition, we grew the cultures overnight in IPTG-free M9 medium and performed multiple washing steps to ensure complete depletion. After IPTG removal, we grew the cells in glucose-containing M9 medium to repress the *lac* operon and prepare for selection. We split the cultures and resuspended them in either M9 medium with 10% glycerol (for a non-inducing condition) or M9 medium with 1 mM lactose (to induce the *lac* operon). We incubated the cells in these conditions for 16 hours and measured cell densities before and after selection by absorbance (optical density) at λ=600 nm (OD_600_). We collected samples before and after selection for high-throughput sequencing to assess the frequency of *lac* operator variants. For amplicon sequencing, we extracted genomic DNA, amplified the mutagenized region of the *lac* operator using barcoded primers, purified and quantified the PCR products, and sequenced them on an Illumina NovaSeq platform (Supplementary Methods 6).

### Calculating fitness

We used a dual sequence filtering approach for reliable fitness estimations. First, we retained only genotypes with a sum of at least 50 read counts across our eight replicates (four replicates before and four after selection). In addition, we only used genotypes detected in at least one replicate before and after selection for further analysis. For these filtered genotypes we calculated relative fitness using a previously established method rooted in population genetic principles (Supplementary Methods 8) (41,44). The method estimates the difference between the growth rate of each variant and a reference variant, because this difference corresponds directly to a variant’s selection coefficient, which is critical for our simulation of adaptive evolution. Using this approach, we estimated fitness for each *lac* operator variant relative to the wild-type (WT) in both glycerol and lactose environments. We then tested the null-hypothesis that the two fitness values are not different using the Wald test, and adjusted p-values with the Benjamini-Hochberg method for false discovery rate control. (Supplementary Methods 8).

### Validating fitness estimates

We validated the fitness estimates from the bulk competition experiment by measuring the growth rates of individual *lac* operator variants. To this end, we isolated 20 sequence-confirmed variants from each of variant categories H_G+L_, H_G_, and L_G+L_ (Figure 2a) of the oligopool libraries, as well as 23 *lac* operator variants from the main CRISPR-Cas9 mutagenesis library. We confirmed the *lac* operator sequences of all these isolates through Sanger sequencing (Supplementary Methods 10).

To assess the growth rate of each variant, we first prepared overnight cultures in M9 medium with 10% glucose, followed by multiple washes to deplete IPTG. We then allowed the variants to grow overnight in M9 medium with either 10% glycerol or 1 mM lactose. Subsequently, we measured growth in three replicate cultures by monitoring optical density (OD_600_) for 24 hours, with data collected every 5 minutes. We calculated the maximal growth rate (fitness) for each variant by fitting tangent lines to the growth curves, and we used the steepest slope as the estimate of the maximal growth rate (Supplementary Methods 10).

In addition to measuring growth rate, we also assessed β-galactosidase activity for individual variants. These experiments started again from overnight cultures that we centrifuged and washed to remove IPTG. We then grew the variants in M9 medium with either 10% glycerol, or 1 mM lactose. We measured β-galactosidase activity by incubating the cells with 2-nitrophenyl β-D-galactopyranoside (ONPG), and monitored the reaction by measuring OD_420_ to calculate Miller units (90). These measurements allowed us to evaluate the relationship between enzyme activity and fitness for both glycerol and lactose environments (Supplementary Methods 10).

### Analysis of the fitness landscape

We constructed the fitness landscape as a network or graph, where each node represents a genotype (a *lac* operator variant). Directed edges connect genotypes differing by a single nucleotide, pointing from the lower fitness genotype to the higher fitness one. This representation also allowed us to identify fitness peaks, evolutionarily accessible paths, and basins of attraction for each peak (Supplementary Methods 11). We analyzed epistasis using four-genotype motifs and quantified diminishing-returns epistasis (Supplementary Methods 11). We performed all analyses using the `igraph` library (version 0.10.8) in Python 3.10.11.

### Adaptive walk simulations

To model adaptive evolution on our fitness landscape, we assumed the strong selection weak mutation (SSWM) regime (56–58). We calculated fixation probabilities for one-mutant neighbors using Kimura’s equation and normalized the probabilities to generate a probability mass function for the simulations. We performed adaptive walk simulations, starting from the 1,000 non-peak variants with the lowest fitness values, executing 1,000 independent simulations per variant. This resulted in a total of 10^6^ adaptive walks, each continuing for up to 50 steps or stopping earlier if a fitness peak had been reached (Supplementary Methods 13).

## Author contributions

G.C. and A.W. conceived the study and designed the experiments. G.C. carried out the experiments. G.C. and A.W. analyzed data. G.C. and A.P. wrote computer code to carry out bioinformatic work, simulations, and analysis. A.W. and G.C. wrote the paper, which was edited by all authors.

## Data and materials availability

Amplicon sequencing data are available from Sequence Read Archive under BioProject accession no. PRJNA1286118. The data generated in this study have been deposited in the Github private repository (https://github.com/GopinathChattopadhyay/lac-operator-fitness-landscape) and will be publicly accessible following publication. For peer review, the same code is available to reviewers as a zipped file named lac-operator-fitness-landscape.zip. Bacterial strains and mutants will be made available under the terms of the Uniform Biological Material Transfer Agreement (UBMTA).

## Code availability

The computer code generated in this study has been deposited in the Github private repository (https://github.com/GopinathChattopadhyay/lac-operator-fitness-landscape) and will be publicly accessible following publication. For peer review, the same code is available to reviewers as a zipped file named lac-operator-fitness-landscape.zip.

