## Supplementary Information for "The fitness landscape of the *E.coli lac* operator is highly rugged in two different environments"

### SUPPLEMENTARY METHODS, TABLES AND FIGURES

### Index

### Supplementary Methods

1. Strains and plasmids
2. Media and reagents
3. Preparation of electro-competent cells using the mannitol-glycerol method
4. Selection of guide RNA (gRNA) spacers
5. Generating *lac* operator variant library through gene editing
6. Bulk competition experiments
7. NGS processing workflow and variant identification
8. Calculating fitness
9. Generating small libraries of *lac* operator site using oligopools
10. Comparison of relative fitness from bulk competition to the growth rate and β-galactosidase activity in single culture
11. Analysis of the fitness landscape
12. Principal component analysis
13. Simulated adaptive walks
14. Generating sequence logos for the *lac* operator site
15. Sensitivity and specificity of peak identification

### Supplementary Results

1. Variation in fitness and *lac* operon expression in operator mutants across environments
2. Nucleotide preferences at *lac* operator sites across different environments
3. Basin Size Increases with Peak Fitness
4. Robustness of peak identification

### Supplementary Tables

S1. Vectors used in this study
S2. *E.coli* strains used in this study
S3. Oligonucleotides used in this study
S4. Sequence analysis pipeline statistics for the main selection experiment
S5. Sequences of Oligopool1 (First [$H_{G+L}$] category variants)
S6. Sequences of Oligopool2 (Second [$H_G$] and third [$H_L$] category variants)
S7. Sequences of Oligopool3 (Fourth category [$L_{G+L}$] variants)
S8. Frequency of epistasis types from network motifs

### Supplementary Figures

**S1. DNA Sequence logo obtained by a previous study**
**S2. Correlation of read coverage in glucose (pre-selection) between four replicates**
**S3. Correlation of read coverage in glycerol (post selection) between four replicates**
**S4. Correlation of read coverage in lactose (post selection) between four replicates**
**S5. Growth curves of 24 *E. coli* populations harboring 24 different *lac* operator variants in individual cultures grown in M9 medium with 10% glycerol**
**S6. Growth curves of 24 *E. coli* populations harboring 24 different *lac* operator variants in individual cultures grown in M9 medium with 1 mM lactose**
**S7. Growth curves of 20 *E. coli* populations harboring 20 different *lac* operator variants from $H_{G+L}$ category in individual cultures grown in M9 medium with 10% glycerol**
**S8. Growth curves of 20 *E. coli* populations harboring 20 different *lac* operator variants from $H_{G+L}$ category in individual cultures grown in M9 medium with 1 mM lactose**
**S9. Growth curves of 20 *E. coli* populations harboring 20 different *lac* operator variants from $H_G$ category in individual cultures grown in M9 medium with 10% glycerol**
**S10. Growth curves of 20 *E. coli* populations harboring 20 different *lac* operator variants from $H_G$ category in individual cultures grown in M9 medium with 1 mM lactose**
**S11. Growth curves of 20 *E. coli* populations harboring 20 different *lac* operator variants from $L_{G+L}$ category in individual cultures grown in M9 medium with 10% glycerol**
**S12. Growth curves of 20 *E. coli* populations harboring 20 different *lac* operator variants from $L_{G+L}$ category in individual cultures grown in M9 medium with 1 mM lactose**
**S13. Correlation between relative growth rates and relative fitness in glycerol and lactose**
**S14. Relative fitness (individual growth assays) measured in two environments.**
**S15. Fitness distribution of lac operator variants across categories**
**S16. β-Galactosidase activity of cells with the WT *lac* operator**
**S17. β-Galactosidase activity of 20 *E. coli* populations harboring 20 different *lac* operator variants from $H_{G+L}$ category in individual cultures grown in M9 medium with 10% glycerol**
**S18. β-Galactosidase activity of 20 *E. coli* populations harboring 20 different *lac* operator variants from $H_{G+L}$ category in individual cultures grown in M9 medium with 1 mM lactose**

**S19. β-Galactosidase activity of 20 *E. coli* populations harboring 20 different *lac* operator variants from $H_G$ category in individual cultures grown in M9 medium with 10% glycerol**
**S20. β-Galactosidase activity of 20 *E. coli* populations harboring 20 different *lac* operator variants from $H_G$ category in individual cultures grown in M9 medium with 1 mM lactose**
**S21. β-Galactosidase activity of 20 *E. coli* populations harboring 20 different *lac* operator variants from $L_{G+L}$ category in individual cultures grown in M9 medium with 10% glycerol**
**S22. β-Galactosidase activity of 20 *E. coli* populations harboring 20 different *lac* operator variants from $L_{G+L}$ category in individual cultures grown in M9 medium with 1 mM lactose**
**S23. Relationship between relative enzyme activity and fitness measured from individual growth assays in different media**
**S24. Fitness distribution of peak genotypes in lactose and glycerol environments.**
**S25. Genetic distance distribution of all peaks in *lac* operator landscape across different environmental conditions**
**S26. Principal component analysis of fitness peaks in the glycerol landscape's genotype space**
**S27. Principal component analysis of fitness peaks in the lactose landscape's genotype space**
**S28. Sequence logo representing the *lac* operator site**
**S29. Fitness peaks in glycerol have larger basins of attraction than in lactose**
**S30. Basin sizes correlate with peak fitness**
**S31. The top 100 peaks have larger basins of attraction sizes than 100 peaks at intermediate fitness and the bottom 100 peaks**
**S32. The basins of attraction associated with the top, intermediate and bottom 100 peaks show high, moderate to low overlap.**
**S33. The top 100 peaks in glycerol have more shared variants than those in lactose**
**S34. Sensitivity of peak identification in glycerol and lactose**
**S35. Specificity of high fitness non-peaks variants in glycerol and lactose**
**S36. Patterns of epistasis across different environments**

### 1. Strains and plasmids

We used *E. coli* strain MR$^S$, which is a derivative of K-12 MG1655 and harbors the amber stop codon suppression mutation *gln*V44 (also known as *sup*E44) (1). This amber stop codon suppressor mutation enables UAG stop codons to be read through and replaced with glutamine (CAG) (2). The mutation rate of the MR$^S$ strain (U = 0.00034 per genome per generation) is similar to that of a wild-type *E. coli* B strain (U = 0.00041 per genome per generation), indicating that the MR$^S$ strain does not cause an excessive number of mutations that could compromise the reliability of our results (1,3).

For gene editing, we utilized the plasmid vectors pKDsgRNA-ack and pCas9-CR4 from the Addgene plasmid repository (4). The pCas9-CR4 plasmid harbours a p15A replication origin (medium copy number, i.e., 20-30 copies per cell) and a chloramphenicol resistance gene (4,5). The pCas9-CR4 plasmid encodes the Cas9 nuclease under the control of the *pTet* promoter, with an SsrA consensus tag added to its C-terminus and a constitutively expressed TetR repressor (4). The SsrA tag facilitates recognition by the ClpP protease, promoting the rapid degradation of Cas9 produced from leaky transcription (4). Cas9 expression is induced as needed using anhydrotetracycline (Cyaman, cat. no. 10009542) as the inducer. The pKDsgRNA-ack plasmid harbours a pSC101 replication origin (low copy, i.e., 3-4 copies per cell) and a spectinomycin resistance gene (4,5). The pKDsgRNA-ack plasmid also includes an arabinose-inducible λ-Red recombination system, with the three genes of the λ-Red system (*gam*, *beta*, and *exo*) controlled by the arabinose-inducible ParaB promoter, and an anhydrotetracycline-inducible sgRNA expression cassette targeting the pCas9 plasmid (4).

A complete list of all vectors and strains used is provided in Tables S1 and S2. For all cloning steps involving plasmids, we used SIG10 HIGH Electrocompetent cells (Sigma-Aldrich, cat. no. CMC0003).

### 2. Media and reagents

As we reported previously (6), to prepare SOB medium we dissolved 25.5 g of its solid stock (VWR J906) in 960 ml of water and autoclaved the medium before use. To prepare SOC medium, we added 20 ml of 1 M D-glucose (Sigma, cat. no. G8270), and 20 ml of 1 M magnesium sulfate (Sigma, cat. no. 230391) to 960 ml of SOB medium.

We purchased M9 minimal salts from Sigma (cat. no. M6030) and prepared it according to the supplier's instructions. We sterilized the solution by autoclaving and supplemented it with 0.2% casamino acids (Merk Millipore, cat. no. 2240), 2 mM magnesium sulfate (Sigma, cat. no. 230391), and 0.1 mM calcium chloride (Sigma, cat. no. C7902). We refer to this solution as the M9 medium throughout this study. Depending on the experiment, we supplemented the M9 medium with either 10% glucose (Sigma G8270), 10% glycerol (Sigma, cat. no. G7757), or 1 mM lactose (Sigma, cat. no. 17814) as the carbon source.

As we reported previously (6), where necessary, we supplemented growth media with 50 µg/ml spectinomycin (Sigma, cat. no. S4014), 34 µg/ml chloramphenicol (Sigma C1919), as well as with 0.2% L-arabinose (Sigma, cat. no. A3256), and 0.16 µg/ml anhydrotetracycline (Cyaman, cat. no. 10009542).

**3. Preparation of electro-competent cells using the mannitol-glycerol method**

We prepared electrocompetent cells using glycerol/mannitol step centrifugation as described previously (4,6,7). Briefly, we diluted an overnight culture 100-fold in 5 ml of SOB medium and incubated the culture until the cell density had reached an $OD_{600}$ of 0.5. Subsequently, we chilled the cells on ice for 6 minutes and then pelleted them by centrifuging for 10 minutes at 3000g at 4°C. After removing the supernatant, we resuspended the cells in 1 ml of ice-cold water and transferred the suspension to a 2 ml microcentrifuge tube. Next, we gently added 1 ml of glycerol-mannitol solution (20% glycerol [Sigma, cat. no. G7757] and 1.5% mannitol [Sigma, cat. no. M4125]) to the bottom of the tube. We centrifuged the tubes again for 10 minutes at 3000g at 4°C and carefully removed the supernatant. Finally, we resuspended the cell pellet in 100-200 µl of glycerol-mannitol solution, and used the freshly prepared competent cells for electroporation.

**4. Selection of guide RNA (gRNA) spacers**

We identified potential gRNA spacer sequences based on available PAM motifs near the *lac* operator (21 bp) in the *E. coli* genome (GenBank accession number

NC_000913). We used Cas-OFFinder, to evaluate off-target sites for the identified candidate gRNA spacers (8).

**5. Generating *lac* operator variant library through gene editing**

Construction of pKDsgRNA-lacO to edit the lac operator site: Here we describe how we constructed the vector pKDsgRNA-lacO, which expresses a guide RNA targeting the chromosomal *lac* operator site, by replacing the gRNA spacer in pKDsgRNA-ack through “round the horn” cloning, a form of inverse polymerase chain reaction (PCR) which can be used for plasmid mutagenesis by introducing mutations through PCR primers (9). The necessary PCR reaction mix contained 2X Verifi Hot Start Mix (Witec, cat. no. PB10.46-05), 0.1 µM primers (Table S3), as well as 0.1 ng of template (pKDsgRNA-ack) DNA. We added nuclease-free water (BioConcept, cat. no. 12931S) to obtain a total volume of 100 µl. The PCR itself used the following thermocycling protocol: 95°C for 3 min, 35 cycles of 95°C for 30 s, 55°C for 30 s, 72°C for 4 mins, followed by 15 min at 72°C after the last cycle. We then purified the PCR product using the Qiagen QIAquick PCR Purification Kit (cat. no. 28106) as per the manufacturer's protocol. After purification, we performed a 20 µl ligation reaction by using 2 µl of purified PCR product, 10 U of T4 DNA ligase (NEB, cat. no. M0202L), and 2 µl of 10× ligation buffer (NEB, cat. no. M0202L). We incubated the reaction for ~16 h at 20-22°C, and inactivated the ligase at 65°C for 10 min. We then transformed 2 µl of the ligation reactions to 25 µl of electrocompetent SIG10 cells and transferred the mixture into a 0.1 cm-gap Gene Pulser/MicroPulser Electroporation Cuvette (Axon Lab, cat. no. EP-201). We electroporated cells at 18 kV/cm using a Micropulser electroporator (Bio-Rad). After electroporation, we added 1 ml of pre-warmed SOC medium, and incubated the culture for 1.5 h at 30°C with shaking at 220 rpm. We verified the correct construction of pKDsgRNA-lacO in five colonies isolate from the electroporated culture by Sanger sequencing (Microsynth AG, Switzerland) using the universal primer pBAD-rev (Table S3). We purified pKDsgRNA-lacO from one of the sequence-validated clones using a Qiagen Plasmid purification kit (cat. no. 27106). Next, we transformed *E. coli* cells that already carried pCas9-CR4 by electroporating it with 2 µl of the newly designed pKDsgRNA-lacO plasmid (1 mm cuvette, 2.4 kV, BioRad MicroPulser). To recover the transformed cells, we incubated them in SOC medium for 2 hours at 30°C, and then plated them on SOC agar plates containing 50 µg/ml spectinomycin and 34 µg/ml chloramphenicol.

We used three clones of MR$^S$/pCas9-CR4/pKD-gRNA-lacO to verify Cas9 activity using a Cas9-induced mortality assay. For this assay, we resuspended bacterial colonies in Dulbecco's Phosphate Buffered Saline (PBS buffer, Sigma, cat. no. D8537) and serially diluted the suspension up to $10^5$-fold. We then plated samples from these dilutions on SOB plates containing 34 µg/ml chloramphenicol and 50 µg/ml spectinomycin, with or without 0.16 µg/ml anhydrotetracycline.

After incubating for 24 hours at 30°C, we compared colony counts on the anhydrotetracycline-induced and uninduced plates, with the expectation that if cell killing is efficient, no colonies will appear on the induced plate. At the very least, there should be a ten-fold reduction in colony counts on the induced plate compared to the uninduced plate. We designated one confirmed clone, based on a greater than 10-fold difference in colony counts between the induced and uninduced plates, as MR$^S$/pCas9-CR4/pKD-gRNA-lacO and used it later as the parental strain to create mutant libraries.

*Design of recombination template for gene editing*: To design the recombination template for *lac* operator gene editing, we used an oligonucleotide that (i) has a sequence complementary to the genomic region containing the target, (ii) is 80-90 nucleotides long, and (iii) has its center no farther than 10 nucleotides from the target position. We computed the GC content, secondary structure and the minimum free energy (ΔG) of the oligonucleotide using the Mfold web server for nucleic acid folding and hybridization prediction (4,10). To assess the secondary structure, we used Mfold with default settings (10). If the minimum free energy (ΔG) value was below -12.5 kcal/mol, we modified the oligonucleotide to minimize secondary structure.

To be able to mutagenize the *lac* operator at multiple positions, we modified the resulting template by designing an oligonucleotide library with several degenerate positions (N), where all four nucleotide bases—A, C, G, and T—are incorporated with equal frequency during synthesis, generating all possible sequence combinations at those positions. The resulting degenerate oligonucleotide LacI-Regular-Oligo is shown in Table S3. This oligonucleotide includes nine consecutive and degenerate bases (N) that correspond to nucleotide positions 1-9 of the *lac* operator (21 bp) on the *E. coli* chromosome. The oligonucleotide library thus contains $4^9$= 262,144 unique nucleotide sequences. We designed the oligonucleotide with three bases at the 5'-end and two bases at the 3'-end that use phosphorothioate bonds for protection against exonucleases (11). We obtained the oligonucleotide from IDT Technologies (Belgium).

_CRISPR-Cas9 mutagenesis_: We prepared an overnight culture by inoculating a single colony of $MR^S$/pCas9-CR4/pKD-gRNA-lacO into 1 ml of SOB medium containing 34 µg/ml chloramphenicol, 50 µg/ml spectinomycin and 1 mM Isopropyl β-D-1-thiogalactopyranoside (IPTG, Merck 11411446001). After 24 hours of incubation at 30°C with shaking at 225 rpm, we transferred 80 µl of the culture into 5 ml of fresh SOB medium with the same antibiotic concentrations and 1 mM IPTG, and continued incubating at 30°C with shaking at 225 rpm. After four hours, we induced the lambda-Red genes by adding 0.2% arabinose.

We harvested the induced culture after 1 hour to prepare electrocompetent cells using the glycerol-mannitol method. Next, we transformed the competent cells with 1 mM of the oligonucleotide template, LacI-Regular-Oligo, using a 1 mm cuvette at 2.4 kV on a BioRad MicroPulser. After transformation, we allowed the cells to recover in 1 ml of SOC medium at 30°C with shaking for one and a half hours. We then serially diluted the culture $10^{-1}$ to $10^{-3}$-fold and plated it onto petri dishes (Thermo Scientific 168381; 150 mm x 21 mm) containing SOC agar with 34 µg/ml chloramphenicol, 50 µg/ml spectinomycin, 0.16 µg/ml anhydrotetracycline, and 1 mM IPTG. After incubating the plates at 30°C for 24 hours, we scraped the bacterial lawn and resuspended it in PBS. We centrifuged the suspension at 3000 g for 20 minutes and then re-suspended the pellet in 20 ml of PBS containing 30% glycerol. Finally, we snap-froze the resuspended cells in liquid nitrogen and stored them at -80°C.

**6. Bulk competition experiments**

_IPTG depletion_: Cells stored at -80°C after mutagenesis of the *lac* operator contain IPTG inducer molecules. IPTG is essential for library creation, because it causes the LacI repressor to detach from DNA – bound LacI repressor which may otherwise interfere with Cas9 activity or the recombination process that is essential for mutagenesis. However, because the IPTG inducer can also affect the fitness of *lac* operator mutations during a bulk competition experiment, it needs to be removed before bulk competition. We thus depleted IPTG molecules by allowing mutagenized cells to grow without it. To this end, we thawed an aliquot of these cells from -80°C on ice and 50 ml of fresh M9 medium with it (10% glucose, 0.2% casamino acid).

We incubated the resulting culture overnight at 30°C with shaking at 225 rpm. Afterward, we washed the overnight culture three times in 100 ml of M9 medium (10% glucose, 0.2% casamino acid), and then resuspended the final pellet in 50 ml of fresh M9 medium (10% glucose, 0.2% casamino acid), continuing to incubate overnight at 30°C with shaking at 225 rpm. We repeated this washing and resuspension process three times to ensure the complete removal of excess IPTG.

Selection: After having depleted the mutagenized cells of IPTG, we began the bulk competition experiment by first ensuring that the *lac* operon remains unexpressed. To this end, we grew four replicate cultures of *lac* operator cells to saturation in 50 ml Falcon tubes (Greiner, cat. no. 227261) with 15 ml of M9 medium (10% glucose, 0.2% casamino acid). We subjected a sample from each replicate population to high-throughput sequencing, to quantify the frequency of the *lac* operator variants before selection.

Next, we split the growing culture into two equal parts, centrifuged them, and discarded the supernatant of glucose-containing M9 medium. We then resuspended one pellet in M9 medium containing 10% glycerol, centrifuged it, and repeated the resuspension and centrifugation steps two more times, for a total of three washes. Similarly, the other pellet was resuspended in M9 medium containing 1 mM lactose as an inducer of the *lac* operon and washed using the same procedure.

We then transferred the resuspended cell pellets into two different media: M9 with 10% glycerol and M9 with 1 mM lactose. For each medium condition, we established four replicate cultures and incubated them for 16 hours at 30°C with shaking at 225 rpm. We measured cell densities before and after selection by recording absorbance at λ=600 nm in 1 cm cuvettes (ratiolab, cat. no. 2712120) using a Genesys 180 UV-Vis spectrophotometer (ThermoFisher). We then subjected a sample from each replicate population to high-throughput sequencing to quantify the frequency of the *lac* operator variants after selection.

Amplicon Sequencing: We sampled cells from the bulk competition experiment both before and after selection to extract genomic DNA using a DNeasy Blood & Tissue Kit (Qiagen). We then amplified the mutagenized region of the *lac* operator from the genomic DNA using the reverse primer ngs_lacO_LIB_R01 (Table S3) and one of 12 forward primers (4 replicates x 3 conditions) ngs_lacO_LIB_F01 to ngs_lacO_LIB_F12, Table S3). The forward primers included 10-letter barcodes, which allowed us to pool the amplification products. We performed the amplification in a 50 µl PCR containing 2X Verifi Hot Start Mix (PB10.46-05), 0.1 µM primers (Table S3), and 0.1 ng of template DNA (genomic DNA) under the following conditions: 95℃ for 3 minutes, followed by 25 cycles of 95°C for 10 seconds, 60°C for 20 seconds, and 72°C for 30 seconds; ending with 72°C for 10 minutes. After amplification, we treated the PCR products with 10 U of DpnI (NEB , cat. no. R0176S) and ExoI (NEB cat. no. M0293S) enzymes at 37°C for 1 hour. We purified the PCR products using a Monarch PCR & DNA Cleanup Kit, estimated their concentration using Qubit dsDNA Quantification Assay Kits (Invitrogen, cat. no. Q32853) pooled them equimolarly, and sent them for paired-end sequencing on the Illumina NovaSeq 6000 platform (Eurofins Genomics, Germany).

#### 7. NGS processing workflow and variant identification

We first examined the quality of the sequencing reads using FastQC (v0.11.9, www.bioinformatics.babraham.ac.uk/projects/fastqc) with default parameters. We used Merging and Tag Sorting services from Eurofins Genomics Design to perform the initial processing, where paired-end reads were merged, and barcodes were sorted according to the specific replicates and conditions from which the samples originated (see Table S3). Following the sorting step, we developed an in-house Python script to filter out sequences that did not match the expected read length of the wild-type (WT) sequence. We then removed reads containing indels or mutations outside the targeted region, identified unique variants within the targeted stretch, and calculated their frequencies. Finally, we exported the frequency table of variants for further analysis. Table S4 shows pipeline statistics and read numbers for different samples for the main selection experiment.

#### 8. Calculating fitness

<u>Sequence Filtering</u>: To ensure the reliability of our fitness estimations, we implemented a dual sequence filtering approach. First, we only retained genotypes with a combined total of at least 50 read counts across all eight replicates (four before and four after selection). Subsequently, we further filtered the data to include only those genotypes that were detected in at least one replicate both before and after selection. This two-step filtering process ensured that the variants selected for fitness

estimation were present across conditions and reduced errors in our fitness estimations. With this filtered dataset, we proceeded to calculate fitness values as previously described (6), with minor modifications detailed below.

*Justification of the Chosen Method*: In competition experiments, relative fitness is often calculated using the ratio of exponential growth rates between the variant of interest and a reference variant, expressed as $r_{variant}/r_{ref}$, where the fitness value of the reference variant equals one. While this approach is common, it has a notable limitation: it does not directly correlate with the selection coefficient of specific variants (12,13). Instead, using the difference ($r_{variant}$ – $r_{ref}$), is more advantageous, as population genetic theory shows that this difference corresponds to the selection coefficient (14). Under this framework, a variant with a growth rate matching the reference yields a relative fitness of zero. Given that selection coefficients are critical for analyzing adaptive evolution across a fitness landscape, we have opted for the latter method, as described in more detail in the following sections.

*The Model*: To estimate relative fitness, we utilized a previously established model for competition among haploid genotypes (14). This model operates under the assumption of exponential growth, continuous time, overlapping generations, and the absence of density- or frequency-dependent selection. Under these conditions, the number $N$ of individuals of a given variant at time $t$ can be expressed as:

$$N_t = N_0 e^{mt} \qquad (1)$$

where $N_0$ represents the initial population size, and $m$ is the Malthusian parameter or intrinsic growth rate for the variant. According to fundamental population genetics, the relative frequency of a variant $i$ that competes with all other variants changes linearly on a logit (logarithm of the odds ratio) scale:

$$ln\frac{p_t}{1-p_t} = ln\frac{p_0}{1-p_0} + (r_i - r)t \qquad (2)$$

Here, $p$ denotes the frequency of variant $i$ at time $t$, $r_i$ is the intrinsic growth rate of that variant, and $r$ is the average growth rate of all variants. The term $(r_i - r)$ indicates the growth rate of variant $i$ relative to the rest of the population during competition, reflecting its relative fitness (12). When time $t$ is expressed in generations, we can convert $r$ to Wrightian fitness using the equation $w = e^r$ (3) (49). We can then denote the number of sequencing-reads for variant $i$ as $N(i)$ and for all other variants as $N(other)$. By substituting these values into equation 2, we can express the variant frequencies in the final part of our analysis as follows:

$$logit\,(p) = ln\frac{p}{1-p} = ln\frac{N(i)}{N(other)} \qquad (4)$$

or equivalently

$$ln\frac{N(i)_t}{N(other)_t} = ln\frac{N(i)_0}{N(other)_0} + (r_i - r_{other})t \qquad (5)$$

*Statistical Estimation of Relative Fitness*: As we described previously (6), we can use equation (5) to estimate the relative fitness $(r_i - r_{other})$ by fitting a generalized linear model (GLM) to the sequencing read data using logistic regression. Specifically, we define this model as:

$$Y = \beta_0 + \beta_1 t + \epsilon \qquad (6)$$

Here, $Y$ represents the log odds ratio of sequencing read counts for the variant of interest compared to all other competing variants, which can be expressed mathematically as $Y = ln\frac{N(i)}{N(other)}$. The quantity $N(i)$ refers to the number of sequencing reads for the variant $i$ of interest, while $N(other)$ encompasses the number of reads from all other variants and can be expressed as $N(other) = N(Total) - N(i)$, where $N(Total)$ is the total number of reads. The parameter $\beta_0$ is the intercept, estimating $Y$ at time 0 (before selection), while $\beta_1$ represents the slope and corresponds to $(r_i - r_{other})$. In addition, $t$ denotes generation time, and $\epsilon$ is an error term. We utilized a generalized linear model with a binomial family distribution due to its use of the logit transformation (as a link function), which aligns with the logit-transformed sequencing read counts in equation (5).

We fitted this generalized linear model to the sequencing read data for each pair of variants using the `glm` function from the `statsmodels` Python package, with the argument `family=Binomial(link=Logit())` (15).

This approach allowed us to obtain estimates of the model coefficients, standard errors, test statistics, and corresponding p-values through the Python `statsmodels` library.

We calculated the standard errors of the model coefficient $\beta_1$ as part of the logistic regression output. Coefficient $\beta_1$ represents the variability or uncertainty associated with the fitness estimated of variant $i$.

To test the hypothesis that the fitness estimator for any one genotype $i$ differs significantly from zero, we employed the Wald test $(H_0\!:\!\beta_1 = 0)$. We applied the Benjamini-Hochberg false discovery rate method to adjust the p-values (187,484 tests for glycerol and 141,149 tests for lactose).

Our analysis quantifies the fitness of each variant relative to the wild-type $(WT)$. Specifically, we aim to estimate $(r_i - r_{WT})$, the difference in growth rates between a variant of interest $i$ and the $WT$. We do so by first fitting the GLM to the sequencing read data using logistic regression to obtain the fitness of the $WT$ $(r_{WT} - r)$, and then repeat this procedure for each variant to obtain $(r_i - r)$. From this calculation we obtain the fitness of variant $i$ relative to the $WT$ as $(r_i - r) - (r_{WT} - r) = (r_i - r_{WT})$. This difference equals the selection coefficient of variant $(i)$, allowing us to determine whether the variant is beneficial $(r_i > r_{WT})$ or deleterious $(r_i < r_{WT})$ relative to the $WT$.

We apply this method to both environments in which we conducted our experiment, i.e., the environment with glycerol as the sole carbon source, in which the WT *lac* operon is repressed, and the environment with lactose as the sole carbon source, in which the WT operon is de-repressed.

#### 9. Generating small libraries of *lac* operator site using oligopools

When we studied the relative fitness of *lac* operator variants in glycerol against their relative fitness in lactose, it became evident that the variants fall into four categories (Figure 2a). To create individual libraries for each of the four categories, we designed three oligopools (10 pmol each), with each pool containing 50 sequences from categories I, II, and IV. Since category III comprised only two sequences, we combined these sequences from the oligopool of category II. See Tables S5-S7 for the sequences in each oligopool. We designed all these oligopools with a 5'-end phosphate modification for protection against exonucleases (11). We obtained the oligopools from IDT Technologies (Belgium).

We prepared an overnight culture of $MR^S$/pCas9-CR4/pKD-gRNA-lacO in SOB medium containing 34 µg/ml chloramphenicol, 50 µg/ml spectinomycin and 1 mM IPTG. After 24 hours of incubation at 30°C, we transferred the culture into fresh SOB medium with the same antibiotic concentrations as well as 1 mM IPTG, and incubated at 30°C for an additional four hours. We then induced the lambda-Red genes by adding 0.2% arabinose, and incubated the culture for one more hour before harvesting it to prepare electrocompetent cells using the glycerol-mannitol method.

Next, we transformed the competent cells with 1 µM of the oligopools template, using a 1 mm cuvette at 2.4 kV on a BioRad MicroPulser. After transformation, we allowed the cells to recover in 1 ml of SOC medium at 30°C with shaking for one and a half hours. We then serially diluted the culture and plated it onto petri dishes containing SOC agar with 34 µg/ml chloramphenicol, 50 µg/ml spectinomycin, 0.16 µg/ml anhydrotetracycline, and 1 mM IPTG. After 24 hours of incubation at 30°C, we scraped the bacterial lawn, resuspended it in PBS, centrifuged at 3000 g for 20 minutes, and re-suspended the pellet in 20 ml PBS with 30% glycerol. We snap-froze the cells in liquid nitrogen and stored them at -80°C. We also isolated clones with multiple *lac* operator variants from each of the three oligopool libraries.

After isolating these clones, we extracted genomic DNA using the Dneasy Blood & Tissue Kit (Qiagen, cat. No. 69504), and amplified the mutagenized *lac* operator region by a PCR with LacI fwd and LacI rev primers (Table S3). We performed the PCR in a 50 µl reaction with 2X Verifi Hot Start Mix (PB10.46-05), 0.1 µM primers (Table S3), and 0.1 ng of template DNA, under the following conditions: 95°C for 3 minutes, followed by 25 cycles of 95°C for 10 seconds, 65°C for 20 seconds, 72°C for 30 seconds, followed by a 10-minute extension at 72°C. We purified the PCR product using the Monarch PCR Cleanup Kit (NEB T1030), and sent it for Sanger sequencing (Microsynth AG, Switzerland) with the LacI reverse primer. Sequencing revealed that the clones included none of the two variants from category III. Consequently, we selected 20 sequence-confirmed variants from the remaining categories (I, II, and IV) for further analysis.

### 10. Comparison of relative fitness from bulk competition to the growth rate and β-galactosidase activity in single culture

Isolating and genotyping individual variants: We validated the fitness estimates from our bulk competition experiment by measuring the growth of *E. coli* populations expressing individual *lac* operator variants. To this end, we isolated 20 sequence-confirmed variants from each of the three categories (I, II, IV) from the oligopool libraries, as described in the previous section.

Additionally, we isolated 23 *lac* operator variants from the library that we had created by CRISPR-Cas9 mutagenesis. Briefly we plated the library stock stored at -80°C onto fresh SOB agar plates containing chloramphenicol and spectinomycin to isolate individual colonies. The cells retained IPTG from the initial plating condition. We then selected 23 individual colonies for further analysis. After isolating these clones, we determined their *lac* operator sequences using Sanger sequencing. To this end, we extracted genomic DNA using the Dneasy Blood & Tissue Kits (Qiagen, cat. No. 69504) and amplified the mutagenized *lac* operator region with PCR, using the primers LacI fwd and LacI rev (Table S3). The amplification occurred in a 50 µl reaction containing 2X Verifi Hot Start Mix (PB10.46-05), 0.1 µM primers (Table S3), and 0.1 ng of template genomic DNA, under the following conditions: 95°C for 3 minutes, followed by 25 cycles of 95°C for 10 seconds, 65°C for 20 seconds, and 72°C for 30 seconds, ending with 72°C for 10 minutes. After amplification, we purified the PCR product using the Monarch PCR & DNA Cleanup Kit (NEB T1030), and sent the purified fragments for Sanger sequencing (Microsynth AG, Switzerland), using the LacI reverse primer (Table S3).

Measuring growth rate: We determined the growth of individual variants in single cultures. To this end, we prepared an overnight culture of each variant in 10 ml M9 medium with 10% glucose at 30°C with shaking at 225 rpm. Afterward, we washed the overnight culture three times in 50 ml of the same medium, and then resuspended the final pellet in 50 ml of this medium, continuing to incubate overnight at 30°C with shaking at 225 rpm. We repeated this process of washing, resuspending, and overnight incubating at 30°C three times to ensure the complete removal of excess IPTG.

After depleting the culture of IPTG, we began the growth experiment in 15-ml Falcon tubes (Greiner, cat. No. 188271) by growing the *lac* operator variants in three replicate cultures until saturation in 5 ml of M9 medium (10% glucose, 0.2% casamino acid), to ensure that the *lac* operon remained unexpressed. Next, we centrifuged each overnight culture and discarded the M9 medium (0.2% casamino acid, 10% glucose). We washed the remaining pellet thrice with M9 medium containing either 10% glycerol or 1 mM lactose. We then inoculated the resuspended culture into M9 medium with either 10% glycerol or 1 mM lactose. After resuspending, we measured the $OD_{600}$ of each culture. For subsequent growth analysis, we diluted cultures directly to $1.6\times10^{8}$ cells/ml ($OD_{600}$=0.2) with either M9 medium with 10% glycerol or M9 medium with 1 mM lactose in 96-well plates (TPP 92096). We incubated the plates for 24 h at 30°C and 1000 rpm in a Spark plate reader (Tecan), measuring the optical density ($OD_{600}$) at 5 min intervals. In this way, we measured the growth dynamics of each isolate in 3 replicate cultures. We $\log_2$-transformed optical density values and scaled time in units of hours. We fitted multiple tangent lines to the growth data of each culture, using a sliding window of three data points (15 minutes) and a step size of one data point (5 minutes). We used the slope of the steepest line (maximum slope) as an estimate of the maximal growth rate (number of cell divisions per hour), our measure of fitness.

*Measuring β-galactosidase activity*: We began the β-galactosidase activity assay analogously to what we just described, by preparing overnight cultures of each *lac* operator variants in three replicate cultures in M9 medium with 10% glucose, followed by multiple washes to deplete IPTG. We grew the variants in M9 medium with either 10% glycerol, or 1 mM lactose after washing.

We performed β-galactosidase activity assays as described previously (16–18). In each well we used 80 µL of an overnight culture, which we permeabilized with a combination of 20 µL of bacterial protein extraction reagent (B-PER, ThermoFisher, cat. no. 90084) and 2 µL of Lysozyme (50 mg/mL, ThermoFisher, cat. no. 90082). We then mixed the cell suspension with 88 µL of Z-buffer (60 mM $Na_2HPO4.2H_2O$ [Sigma, cat. no. 71643], 40 mM $NaH_2PO4.2H_2O$ [Riedel-de Haën, cat. no. 04269], 10 mM KCl [Sigma, cat. no. P3911], 1 mM MgSO4 [Sigma, cat. no. 63136], and 50 mM β-mercaptoethanol, pH 7 [Amresco, cat. no. 1328-0482). We started the β-galactosidase reaction by adding 10 µL (4 mM) of the substrate 2-nitrophenyl β-D-galactopyranoside (ONPG, ThermoFisher, cat. no. 34055). We incubated the 96-well plates for 1 hour at 30°C in a Spark plate reader (Tecan), taking $OD_{420}$ and $OD_{600}$ measurements every 45 s, and shaking the samples at 275 rpm. We used the resulting $OD_{420}$ readings to calculate Miller Units (19) using the formula

$$Miller\ Units = \frac{1000 \times (\mathrm{OD}_{420}/\mathrm{min})}{\text{Initial culture } \mathrm{OD}_{600} \times \text{volume used (ml)}} \qquad (7)$$

For 80 µL culture in 200 µL solution this is equivalent to

$$Miller\ Units = \frac{5000 \times (\mathrm{OD}_{420}/\mathrm{min})}{\text{Initial culture } \mathrm{OD}_{600}} \qquad (8)$$

We calculated the mean and standard deviation of this quantity from our three independent replicate experiments. Using the resulting β-galactosidase activity measurements, we assessed the relationship between enzyme activity and fitness in both glycerol and lactose environments.

For glycerol, we performed a linear regression analysis to evaluate the correlation between activity and fitness. To this end, we fitted a linear model and computed Pearson's correlation coefficient (*r*) along with the corresponding P-value of the null hypothesis that *r*=0. For lactose, we observed a nonlinear relationship between activity and fitness and therefore employed a quadratic model, fitting the second-degree polynomial

$$y = ax^2 + bx + c, \qquad (9)$$

where $y$ represents fitness, $x$ represents β-galactosidase activity and $a$, $b$, $c$ are the fitted polynomial coefficients. To assess the goodness of fit, we computed the coefficient of determination $(R^2)$, i.e.,

$$R^2 = 1 - \frac{\sum\left(fitness - y_{pred}\right)^2}{\sum\left(fitness - \overline{fitness}\right)^2}, \qquad (10)$$

where $y_{pred}$ represents the predicted fitness values from the quadratic fit and $\overline{fitness}$ is the mean fitness of the observed fitness values across all *lac* operator variants. To compare the quadratic model with a simpler linear model, we tested the null hypothesis ($H_0$) that the quadratic term is not significantly different from zero, i.e., the data can be explained by a linear model. To this end, we performed an F-test and computed the F-statistic and the corresponding P-value (20).

We also compared β-galactosidase activity between glycerol and lactose environments by fitting a logistic (sigmoidal) model to capture the relationship between activity levels in the two environments. For this purpose, we used the logistic function

$$y = \frac{a}{1 + e^{b(x-c)}} + d, \quad (11)$$

where $y$ represents β-galactosidase activity in lactose, $x$ represents activity in glycerol, and $a$, $b$, $c$, and $d$ are fitted parameters. We computed again the $R^2$ value to assess the quality of the fit, and calculated P-values for each parameter to test the null-hypothesis that each parameter is not significantly different from zero (i.e., no effect of that parameter on the relationship between activity in glycerol and lactose).

**11. Analysis of the fitness landscape**

*Constructing a network of TFBS variants*: We represent the fitness landscape we study as a network or graph (21–23). Specifically, using the fitness data obtained from each environment (glycerol and lactose), we built a network in which each node corresponds to a genotype (TFBS variant). Directed edges (arrows) connect genotypes that differ in exactly one nucleotide, and point from the genotype with lower fitness to the genotype with higher fitness. We performed all analyses on the largest (giant) connected component of this network, which comprises 99.999% (187483/187484) of the nodes in glycerol and 99.998% (141147/141149) of the nodes in lactose. We used the python `igraph` (version 0.10.8) library for all network-based analyses, and its function `component` (`mode=weak`) to extract the giant component (24).

*Fitness Peaks*: A fitness peak is a genotype (*lac* operator variant) that has a higher fitness than all its neighbors. To identify fitness peaks, we defined a function that iterates through each node in the graph and compared its fitness to that of its neighbors.

*(Accessible) paths through the network*: An evolutionarily accessible path is a sequence of mutational steps (edges), in which every mutation increases fitness. We identified the evolutionary accessible paths from each genotype to each fitness peak, with the `get_all_shortest_paths` function from the `igraph` library. In addition to enumerating evolutionarily accessible paths from any one variant to a peak, we also enumerated the total number of shortest paths from each variant to each peak.

*Basins of attraction*: The basin of attraction of a fitness peak is the set of all genotypes from which a given peak is accessible, i.e., at least one accessible path exists from the genotype to the peak. The basin's size is the number of genotypes in the basin. To determine the accessibility of fitness peaks from other genotypes in the landscape, we employed the `distances` function from the `igraph` library, which computes the shortest accessible paths from one set of variants to another set of variants. The function also identifies pairs of variants with no accessible paths between them, which enabled us to determine the basin size.

*Overlap between Basins*: The basins of attraction for different fitness peaks may overlap, i.e., they may share some variants. To assess the overlap between two basins $B1$ and $B2$, we calculated the Jaccard similarity coefficient $J$, defined as (6,25,26)

$$J = \left|\frac{B1 \cap B2}{B1 \cup B2}\right| \quad (12)$$

*Detecting sign epistasis*: Epistasis is the non-additive interaction of two or more mutations in their effect on fitness. It can hamper adaptive evolution (27). Epistasis between two mutations can be classified as magnitude, simple sign, or reciprocal sign epistasis, depending on whether mutations individually or together have positive or negative effects on fitness (27–29). In magnitude epistasis, the effect of a mutation on fitness varies depending on the genetic background, but its overall sign (positive/beneficial or negative/deleterious) remains unchanged. Simple sign epistasis occurs when a single mutant has lower fitness than both the WT and the double mutant, while the other single mutant has a fitness that is intermediate between the WT and the double mutant. Reciprocal sign epistasis arises when both mutations individually reduce fitness, but their combination restores or even increases fitness.

Notably, the presence of reciprocal sign epistasis is a necessary condition for the existence of multiple peaks in an adaptive landscape (28,29).

To quantify epistasis, we analyzed motifs comprising four adjacent nodes ("squares") within a genotype network, which correspond to quadruplets of a WT, two single mutants, and a double mutant (see Supplementary Figure S36a for more details on epistasis motifs). To this end we used the `motifs_randesu` function from the `igraph` library in python. We categorized epistasis for each motif along a single axis by selecting the highest-fitness sequence as the double mutant and classifying interactions into three groups: no sign epistasis, simple sign epistasis, and reciprocal sign epistasis. The no sign epistasis category includes both magnitude epistasis and additivity (no epistasis), because neither affects peak accessibility within an adaptive landscape (30–32). We determined the proportion of all identified motifs that falls into each of these three categories (Supplementary Table S8 and Supplementary Figure S36).

*Detecting diminishing-returns epistasis*: Diminishing-returns epistasis is a kind of epistasis in which the effect of beneficial mutations decreases as the fitness of the genetic background increases. It manifests as decelerating fitness improvements during adaptive evolution over time (33–35). This pattern is important to study in the context of adaptive landscapes because it can help to explain declining adaptability in evolving populations (33–37).

To detect diminishing return epistasis, we analyzed the fitness data of genetic interactions using `igraph` (6). Specifically, we analyzed fitness differences between neighboring genotypes from the network's edges, focusing on the relationship between the fitness of the background (fitness of the source genotype) and the resulting fitness gain (the difference in fitness between the target and source genotype).

**12. Principal component analysis**

To explore the proximity of peaks and other variants in sequence space, we applied principal component analysis (PCA) with a one-hot encoding of our DNA sequences. This encoding method converts each nucleotide in a sequence into a binary vector of length 4, resulting in a 4xL binary matrix for sequences of length L. We then applied

PCA to this matrix using the `PCA` function from the `sklearn.decomposition` module.

The sequence space underlying our landscape is high-dimensional, resembling a Hamming graph, in which more than two alleles can exist at each site (38). While PCA is widely used to visualize complex genotype spaces, it has limitations in capturing the full-extent of genetic variation. Specifically, each principal component accounts for only a small fraction of the total variance (39–41). For example, the first principal component (PC1) explained only 4% of the variance in our data (Supplementary Figures S26b and S27b). Projecting peak and non-peak genotypes onto the first five PCA components illustrates that peaks are not highly clustered in genotype space (Supplementary Figure S26-S27).

**13. Simulated adaptive walks**

To model adaptive evolution on our fitness landscape, we assumed that evolution occurs in the strong selection weak mutation (SSWM) regime (42–44), because the likelihood of a mutation occurring in the part of the *lac* operator we study is very small (9 positions × 2.2 × $10^{-10}$ substitutions per site per generation in the *E.coli* genome) (45). This small mutation rate implies that one beneficial mutation is likely to become fixed before the next beneficial mutation arises that will eventually go to fixation, resulting in populations that harbor only one segregating mutation at any given time. Thus, adaptive evolution can be conceptualized as a sequence of adaptive walks, wherein a population starts at a single genotypic position in the landscape and progresses to adjacent positions through successive mutation-fixation events until reaching a fitness peak.

<u>*Adaptive walks using Kimura's fixation probability*</u>: We calculated the fixation probability of each one-mutant neighbor of a given variant using an equation originally derived by Kimura (14,46,47),

$$F_i = \frac{1 - e^{-2Nsp}}{1 - e^{-2Ns}} \qquad (13)$$

Here $F_i$ is the fixation probability of mutant $i$, $s$ represents the selection coefficient of the mutant, i.e., the fitness difference between the mutant and the focal variant, $p$ is the initial frequency of the mutant (assumed to be $p = 1/N$), and $N$ is the population size. We set the population size to $10^8$ to reflect estimates of the *E. coli* population size (48).

After calculating the fixation probabilities of all neighbors of a given variant, we normalized these probabilities to the interval (0,1), i.e.,

$$F_{i,norm} = \frac{F_i}{\sum_j F_j} \qquad (14)$$

This normalization step is important, because it transforms the fixation probabilities into a valid probability mass function, which is essential for our subsequent probabilistic simulations.

With these normalized fixation probabilities in hand, we then defined a new, bidirected network, in which a pair of directed edges pointing in opposite directions connects each pair of neighboring genotypes. We assigned the normalized fixation probabilities as edge weights to these edges.

We then performed adaptive walk simulations on this network. Specifically, we chose the 1,000 non-peak variants with the lowest fitness values as starting points for these random walks. From each starting point, we executed 1,000 independent simulations, resulting in a total of $10^6$ adaptive walks. Each adaptive walk continued for up to 50 steps or until it reached a peak, whichever came first.

**14. Generating sequence logos for the *lac* operator site**

We aligned the sequences of the top 100 genotypes or top 100 fitness peaks, obtained either from glycerol or lactose environments, to construct sequence logos for the *lac* operator site, as described previously (49). To generate the sequence logo, we first calculated the counts of A, T, G, and C bases at each position in the alignment, and normalized these counts to derive a probability matrix. Specifically, we calculated the probability of encountering each base at a given position as

$$p_i = \frac{count_i}{\sum_i count_i} \qquad (15)$$

where $count_i$ represents the number of occurrences of base $i$ at the given position, and the sum is the total count of all bases at that position. To avoid zero probabilities for any one base, we added pseudocounts, small values to prevent zero probabilities when a nucleotide is absent at a specific position. We used a pseudocount of $1 \times 10^{-10}$ in our analysis.

Next, we calculated the Shannon entropy for each position using the equation:

$$H(x) = -\sum_i p_i log_2(p_i) \quad (16)$$

where $p_i$ is the probability of finding nucleotide $i$ at a given position. The Shannon entropy quantifies the amount of information at each position, with higher entropy indicating greater variability.

We then calculated the normalized information content of each base at each position by multiplying the probability of encountering each base at the position by its corresponding entropy value, i.e.,

$$IC_{ij} = p_{ij} \times H(x_j) \quad (17)$$

where $IC_{ij}$ represents the information content for base $i$ at position $j$, $p_{ij}$ is the probability of base $i$ at position $j$, and $H(x_j)$ is the Shannon entropy at position $j$. We used this normalized information content to display the sequence logo, in which the height of each letter reflects the information content at each position.

**15. Sensitivity and specificity of peak identification**

To determine the robustness of our peak-calling procedure (see Method 11: Fitness Peaks), we performed sensitivity and specificity analyses.

<u>Sensitivity:</u> Sensitivity quantifies the ability to correctly identify true peaks, accounting for sampling noise in sequencing data. Peak identification relies on accurately estimating the fitness difference between the focal genotype (the peak) and its neighboring genotypes. We had estimated this fitness difference as a slope in our generalized model (see Method 8), which integrates sequencing read counts from two genotypes across two time points (before and after selection) and from four replicate libraries. Consequently, each coefficient estimate for each genotype-neighbor pair is influenced by sampling noise from 16 data points ($2 \times 2 \times 4 = 16$).

For a fitness peak, the fitness difference relative to all its neighbors must be positive, meaning the peak must have higher fitness than all adjacent genotypes. Therefore, peak determination is affected by multiple sources of sampling error, which depend on sequencing coverage and the specific set of neighboring genotypes.

Given the complexity of our experimental design, we estimated sensitivity using synthetic datasets sampled from the same binomial distribution we had used for estimating slope coefficients in our model. This approach allowed us to assess how frequently our procedure correctly identifies a fitness peak.

We conducted the sensitivity analysis for each of approximately 9,000 putative fitness peaks within the fitness landscape. For each putative peak, we proceeded as follows:

1. Using experimental data for the peak and its neighbors, we parametrized the binomial distribution $B(n_k, p_i)$, where $n_k$ represents the total number of sequencing reads in replicate library $k$, and $p_i$ is the probability of observing genotype $i$ in that library (estimated from the genotype frequency).

2. We generated 50 independent synthetic datasets for each putative peak by sampling from the parameterized binomial distributions $B(n_k, p_i)$. Each synthetic dataset included sampled read counts for the putative peak and its 1-mutational-step neighbors.

3. For each synthetic dataset, we estimated fitness differences between the putative peak and its neighbors using the same procedure as for the real data (see Method 8). Based on these estimates, we classified the peak in each synthetic dataset as either a true positive (TP) or a false negative (FN). Specifically, we considered a peak a FN if at least one neighboring genotype exhibited higher fitness, meaning the putative peak would not be identified as a peak in that dataset. Conversely, we considered a peak a TP if the peak retained the highest fitness among its neighbors. While we cannot determine with absolute certainty whether a peak in the experimental data is truly a peak, in the synthetic datasets we assume all peaks are “true” by definition, because we generated their data from distributions with known parameters.

4. We calculated sensitivity for each putative peak in the landscape as the proportion of synthetic datasets in which this peak was a true positive:

$$\text{Sensitivity} = \frac{TP}{(TP + FN)} = \frac{TP}{50} \quad (18)$$

The higher the sensitivity, the more accurately the peak-calling method identifies true peaks. A sensitivity value of one indicates that the method correctly identified the putative peak in all 50 synthetic datasets. Sensitivity varies across different putative peaks due to differences in sequencing read coverage and the magnitude of fitness differences between each peak and its neighboring genotypes.

<u>*Specificity*</u>: Specificity measures how often genotypes that are not peaks are mistakenly identified as peaks due to sampling noise in the sequencing data. To quantify specificity, we followed a similar approach as just described, i.e., we sampled synthetic datasets from binomial distributions parameterized by our sequence data.

More specifically, we focused in this analysis on non-peak genotypes with fitness exceeding 0.5 (n=1431) in glycerol, and non-peaks with fitness exceeding zero (n=32) in lactose, because we were most interested in high fitness genotypes. By focusing on high fitness non-peaks, we could better assess the specificity of the peak identification process, ensuring that high fitness non-peak genotypes are not incorrectly classified as peaks due to sampling or modeling noise.

For each selected non-peak genotype, we generated 50 synthetic datasets using a binomial distribution $B(n_k, p_i)$ whose parameters are derived from our sequencing data. Each synthetic dataset included four independently sampled read counts at two time points for the non-peak genotype and all its neighbors. We then applied our generalized linear model (see Method 8) to estimate fitness difference between the non-peak genotype and its neighbors.

If a non-peak genotype appeared to be peak in a synthetic dataset (i.e., if its fitness was higher than that of all its neighbors), we considered it a false positive (FP). Otherwise, i.e., if it appeared to be a non-peak, we considered it to be a true negative (TN). We calculated specificity as the fraction of synthetic datasets (out of 50) in which the non-peak genotype appeared was a true negative:

$$\text{Specificity} = \frac{TN}{(TN + FP)} = \frac{TN}{50} \quad (19)$$

A higher specificity indicates greater accuracy of the peak-calling method. A specificity of one means that a particular non-peak genotype was correctly identified as not being a peak in all 50 synthetic datasets.

### Supplementary Results

#### 1. Variation in fitness and *lac* operon expression in operator mutants across environments

Fitness values are positively correlated in glycerol and lactose environments both during bulk competition (Pearson's r= 0.473, P<2.0×$10^{-308}$, Spearman's ρ= 0.433, P<2.0×$10^{-308}$, N= 133888, Figure 2a) and for individual growth assays (Pearson's r= 0.75, P= 2.27×$10^{-12}$, N=61, Figure S14). In the following sections, we compare fitness values across environments for the four categories of variants defined in the main text, beginning with $H_{G+L}$ variants, which have high fitness in both environments.

In the glycerol environment, $H_{G+L}$ variants exhibit higher fitness than the wild-type (WT) in our bulk competition experiment (mean±SD= 0.651±0.257, N=321, Figure S15). They also grow more rapidly in our individual growth assays (mean±SD= 2.605±0.708, N=20, Figure 2b, Figure S7), and express less β-galactosidase (mean±SD= 0.608±0.073, N=20, Figure 2c, Figure S17). Because *lac* operon expression would not benefit a cell in glycerol, these observations suggest that the $H_{G+L}$ variants repress the *lac* operon strongly, which is crucial for their fitness in this environment.

In lactose, where *lac* operon expression is important, $H_{G+L}$ variants not only show higher fitness than the WT during bulk competition (mean±SD= 0.172±0.168, N=321, Figure S15), but also during our individual growth assay (mean±SD= 1.116±0.160, N=20, Figure 2b, Figure S8). And they show higher β-galactosidase activity than WT (mean±SD= 1.802 ± 0.573, N=20, Figure 2c, Figure S18). In sum, our measurements of individual variants support the notion that $H_{G+L}$ variants have high fitness in both glycerol and lactose, because they are highly effective in repressing the *lac* operon in glycerol and activating it in lactose.

For $H_G$ variants, which are fitter than the WT in bulk competition on glycerol (mean±SD=0.174±0.152, N=26,567, Figure S15), a different pattern emerges. Individual growth assays show that they grow more rapidly (mean±SD= 1.163±0.216, N=20, Figure 2b, Figure S9) and express less β-galactosidase (mean±SD= 0.942±0.111, N=20, Figure 2c, Figure S19). These observations suggest that the $H_G$ variants effectively repress the *lac* operon, which is important for their fitness in this environment. However, in lactose, $H_G$ variants show notably smaller fitness than the WT both during bulk competition (mean±SD =-0.970±0.405, N=26,567, Figure S15), and in our individual growth assays (mean±SD= 0.613±0.060, N=20, Figure 2b, Figure S10). They also show lower β-galactosidase activity (mean±SD= 0.769±0.051, N=20, Figure 2c, Figure S20). In sum, our measurements suggest that $H_G$ variants are well-repressed in glycerol but have reduced ability to activate the *lac* operon in lactose, which may explain their lower fitness in lactose.

The few $H_L$ variants had lower fitness in glycerol than the WT in our bulk competition experiment (mean±SD= -0.026±0.029, N=2, Figure S15), indicating that these variants may not effectively repress the *lac* operon. In lactose, the fitness of $H_L$ mutants is slightly higher than the WT during bulk competition (mean±SD= 0.013±0.012, N=2, Figure S15), indicating that these variants are able to activate the *lac* operon to some degree, supporting their growth in this environment. Because there are only two $H_L$ variants we were not able to obtain clones for them by sequencing individual colonies derived from the mixed ($H_G$ and $H_L$) oligopool library. We could therefore not measure their individual growth rate and β-galactosidase activity.

Finally, $L_{G+L}$ mutants in glycerol show lower fitness than the WT during bulk competition (mean±SD= -0.331±0.211, N=106,998, Figure S15). Individual growth assays show that they grow slowly (mean±SD= 0.940±0.096, N=20, Figure 2b, Figure S11) and express higher β-galactosidase (mean±SD= 1.328±0.149, N=20, Figure 2c, Figure S21), suggesting that these variants are unable to effectively repress the *lac* operon. In lactose, $L_{G+L}$ variants also show a decrease in fitness compared to WT both during bulk competition (mean±SD= -1.347±0.356, N=106,998, Figure S15), and in our individual growth assay (mean±SD= 0.481±0.061, N=20, Figure 2b, Figure S12) and express less β-galactosidase (mean±SD= 0.644±0.047, N=20, Figure 2c, Figure S22). Overall, our measurements suggest that $L_{G+L}$ variants have reduced ability both in repressing the *lac* operon in glycerol and activating it in lactose, which may explain their low fitness in both environments.

To summarize, in glycerol, $H_{G+L}$ variants exhibit significantly higher fitness during both bulk competition and individual growth assays, and express lower β-galactosidase compared to both $H_G$ (fitness: U = 8,075,114, P= $2.57\times10^{-167}$; growth rate: t = 8.71, P= $1.15\times10^{-8}$; activity: t = -11.25, P= $8.88\times10^{-13}$) and $L_{G+L}$ (fitness: U = 34,346,358, P= $8.41\times10^{-211}$; growth rate: t = 10.42, P= $1.84\times10^{-9}$; activity: t = -19.39, P= $1.37\times10^{-17}$) variants. This suggests that $H_{G+L}$ variants are better at repressing the *lac* operon compared to $H_G$ and $L_{G+L}$ variants. $H_G$ variants show higher fitness during both bulk competition and individual growth assay and express lower β-galactosidase than $L_{G+L}$ variants (fitness: U = 2,842,615,866, $P < 2.0\times10^{-308}$; growth rate: t = 4.21, $P= 2.65\times10^{-4}$; activity: t = -9.25, $P= 6.05\times10^{-11}$), indicating that they are also better at repressing the *lac* operon in glycerol compared to $L_{G+L}$ variants.

In lactose, $H_{G+L}$ variants also exhibit significantly higher fitness, during both bulk competition and individual growth assay, and express higher β-galactosidase than both $H_G$ (fitness: U = 8,528,007, $P= 6.36\times10^{-209}$; growth rate: t = 13.20, $P= 1.49\times10^{-12}$; activity: t = 8.03, $P= 1.41\times10^{-7}$) and $L_{G+L}$ (fitness: U = 34,346,358, $P= 8.41\times10^{-211}$; growth rate: t = 16.61, $P= 8.12\times10^{-15}$; activity: t = 9.01, $P= 2.42\times10^{-8}$) variants. This shows that $H_{G+L}$ variants are more effective at activating (de-repressing) the *lac* operon in lactose compared to $H_G$ and $L_{G+L}$ variants. $H_G$ variants have significantly higher fitness, during both bulk competition and individual growth assay and express higher β-galactosidase than $L_{G+L}$ variants (fitness: U = 2,169,463,447, $P < 2.0\times10^{-308}$; growth rate: t = 6.92, $P= 3.20\times10^{-8}$; activity: t = 8.04, $P= 1.04\times10^{-9}$), indicating that they are slightly more efficient than $L_{G+L}$ variants at activating (de-repressing) the *lac* operon in lactose.

**2. Nucleotide preferences at** lac **operator sites across different environments**

We examined which nucleotides occur preferentially at different positions of the *lac* operator in the top 100 genotypes (Figure S28a-b) as well as in the top 100 fitness peaks (Figure S28c-d). To this end, we first aligned either the top 100 sequences or the genotypes of the top 100 peaks. We then counted the occurrences of A, T, G, and C, at each nucleotide site, and normalized these counts to obtain frequency values. The resulting frequency matrix can be represented as a heatmap, and it can be transformed into a visual sequence logo representation by scaling nucleotide frequencies according to their contribution to Shannon entropy (Figure S28). This frequency matrix shows conserved nucleotides similar to the WT, along with additional variants that contribute to high fitness. For example, in lactose the preferred nucleotides at positions 1, 6, and 9 are identical to the WT (Figure S28b,d). We emphasize that this logo does not necessarily reflect the DNA binding strength of the LacI repressor to DNA. The reason is that it is based on in vivo fitness measurements rather than DNA-protein binding assays, and fitness can be affected by multiple factors, including an interplay between CAP and LacI binding. It is thus to be expected that the sequence conservation we observe in vivo does not entirely match previous in vitro binding data (Figure S1).

#### 3. Basin Size Increases with Peak Fitness

The association between the fitness of a peak and the size of its basin of attraction becomes evident if we compare the basin sizes of the 100 lowest fitness peaks with those of 100 peaks of intermediate fitness, and with those of the top 100 fitness peaks. In glycerol, the bottom 100 peaks generally have smaller basin sizes (median: 22,150, 12.75±8.83% of all variants) than the intermediate 100 peaks (median: 73,032, 41.56±8.23%) and the top 100 peaks (median: 79,102, 45.51±8.28%, Figure S31a), a significant difference between each pair of these groups (Mann–Whitney U test, $P=2.78\times10^{-32}$ between bottom and intermediate, as well as between bottom and top peaks, $P=6.8\times10^{-32}$ and between top and intermediate peaks, $P=1.37\times10^{-3}$). The same holds in lactose, where the bottom 100 peaks have smaller basin sizes (median: 1,525, 0.39±1.62% of variants) than the intermediate 100 (median: 32,441, 26.54±9.79%) and the top 100 peaks (median: 38,704, 30.11±9.24%, Figure S31b), a significant difference between each pair of these groups (Mann–Whitney U test, $P=6.94\times10^{-32}$ between bottom and intermediate, as well as between bottom and top peaks, $P=3.07\times10^{-34}$ and between top and intermediate peaks, $P=1.09\times10^{-3}$).

#### 4. Robustness of peak identification

To assess how robust our assessment of peak identity was in the *lac* operator regulatory landscape, we performed a sensitivity analysis (see Methods 15: Sensitivity). In this analysis, we evaluated the reliability of peak identification while considering sampling errors in our sequencing data. We determined these sampling errors using a binomial distribution, which we parameterized from our sequencing read counts for each fitness peak and its one-mutation neighbors. By sampling synthetic data from this distribution 50 times and repeating the peak identification process, we found that our procedure for determining peaks is not highly sensitive to sampling noise. Specifically, for the vast majority of peaks (9164 out of 9183) in glycerol, we found a sensitivity (true positive rate) equal to 0.9 (Figure S34a). Furthermore, for the top 100 high-fitness peaks, which are the most important for our analysis, the sensitivity also exceeds 0.9 (Figure S34b). In lactose, 6783 out of a total of 9074 peaks had a sensitivity of 0.9 or higher (Figure S34c), which is lower than in glycerol.

However, 89 of the top 100 highest-fitness peaks in lactose (fitness > 0.389) also had a sensitivity exceeding 0.9 (Figure S34d).

In addition, we estimated the probability of falsely classifying a non-peak genotype as a peak. In this analysis we focused on 1431 non-peak genotypes with fitness values exceeding 0.5 in glycerol, and 32 non-peak genotypes with fitness exceeding the WT fitness of zero in lactose. These genotypes were not originally classified as peaks (Figure S35). We followed the same procedure of parameterizing the binomial sampling distribution with actual sequence read count data, and asked for each of these putative non-peak genotypes, how frequently noise in our data would lead them to be falsely identified as peaks. We found that false positives were rare, despite the high fitness values of the non-peaks (Figure S35). More precisely, the average specificity (mean of true negatives [TN] across 50 replicate synthetic data sets, see Method 15: Specificity) of peak identification was 0.943 in glycerol and 0.896 in lactose. The average false positive rate (mean of false positives [FP] across 50 replicates) across the 1432 glycerol and 32 lactose non-peak genotypes was 0.057 and 0.105 respectively, further highlighting the reliability of our peak identification process.

To conclude, our analysis shows that despite sampling noise in the sequence data, our peak identification is reliable, including the identification of high-fitness peaks.

### Supplementary Tables

**Table S1.**

Vectors used in this study

| Name | Genotype | Selective antibiotics (concentration µg/ml) | Source | T (°C) | Description |
|---|---|---|---|---|---|
| pKDsgRNA-*ack* | exo, beta, gam, sgRNA-ack | Spectinomycin (50) | Addgene, 62654 (4) | 30 | Vector for expressing gRNA targeting ack |
| pCas9-CR4 | cas9-CR4 | Chloramphenicol (34) | Addgene, 62655 (4) | 37 | Vector for expressing cas9 |
| pKDsgRNA-*lacO* | exo, beta, gam, sgRNA (*lac* operator site) | Spectinomycin (50) | This study | 30 | Vector for expressing gRNA targeting *lac* operator site |

**Table S2.**

*E.coli* strains used in this study

| Name | Genotype | Selective antibiotics (concentration µg/ml) | Source | T, °C | Description |
|---|---|---|---|---|---|
| $MR^S$ | F-, fhuA2, lacY1, tsx-1 or tsx-70, glnV44 (AS), gal-6, λ-, xyl-7, mtlA2, cycA30::Tn10 | | Wagner lab stock (1) | 37 | Derivative of K-12 MG1655 with wild-type *mutL*. Amber (UAG) suppressor. |
| $MR^S$/ pCas9-CR4 | cas9-CR4 | Chloramphenicol (34) | This study | 37 | $MR^S$ strain that harbours the pCas9-CR4 plasmid. |
| $MR^S$/pCas9-CR4/pKD sgRNA-*lacO* | cas9-CR4, exo, beta, gam, sgRNA (*lac* operator site) | Spectinomycin (50) Chloramphenicol (34) | This study | 30 | Parental strain for creating mutant libraries in *lac* operator site. λ-red recombination system inducible by 0.2 % arabinose. |
| SIG10 HIGH Electro-competent cells | | | Sigma-Aldrich CMC0003 | 37 | Strain for molecular cloning |

**Table S3.**

Oligonucleotides used in this study. Capital letters in the right-hand column show degenerate nucleotides (N = A, T, G, C). * denotes phosphorothioate bond at the 5' and 3' end of the oligonucleotide.

| Primer | Sequence |
|---|---|
| LacI fwd | ggcctcttcgctattacgccagct |
| LacI Rev | accctggcgcccaatacgcaaacc |
| sgRNA$_{CGG}$ target F | atgttgtgtggaattgtgaggttttagagctagaaatagcaag |
| sgRNA$_{CGG}$ target R | ctcacaattccacacaacatgtgctcagtatctctatcactga |
| PtetR | PO4-gtgctcagtatctctatcactga |
| LacI-Regular-Oligo (Main experiment) | c*a*t*agctgtttcctgtgtgaaattgttatccgNNNNNNNNNccacacaacatacgagccggaagcataaagtgtaaagcct*g*g |
| ngs_lacO_LIB_F01 | gatctcattcaaaacgacggccagtgaatccgta |
| ngs_lacO_LIB_F02 | gtaaggctccaaaacgacggccagtgaatccgta |
| ngs_lacO_LIB_F03 | tatcatgcagaaaacgacggccagtgaatccgta |
| ngs_lacO_LIB_F04 | atctgcgtacaaaacgacggccagtgaatccgta |
| ngs_lacO_LIB_F05 | gattgcacgcaaaacgacggccagtgaatccgta |
| ngs_lacO_LIB_F06 | ttcacggaagaaaacgacggccagtgaatccgta |
| ngs_lacO_LIB_F07 | tgctaacttcaaaacgacggccagtgaatccgta |
| ngs_lacO_LIB_F08 | atagcagtgcaaaacgacggccagtgaatccgta |
| ngs_lacO_LIB_F09 | tcatggaatcaaaacgacggccagtgaatccgta |
| ngs_lacO_LIB_F10 | cgacgtagtcaaaacgacggccagtgaatccgta |
| ngs_lacO_LIB_F11 | tcaatgatcgaaaacgacggccagtgaatccgta |
| ngs_lacO_LIB_F12 | gatatagctcaaaacgacggccagtgaatccgta |
| ngs_lacO_LIB_R01 | tagctcactcattaggcaccccag |
| pBAD Reverse | gatttaatctgtatcagg |

**Table S4.**

Sequence analysis pipeline statistics for the main selection experiment

| Pipeline step | Number of reads | Fraction retained from previous step |
|---|---|---|
| **Glucose environment** | | |
| Merging | 59518391 | 1.000 |
| Demultiplexing | 53902578 | 0.906 |
| Length Trimming | 49123301 | 0.911 |
| **Glycerol environment** | | |
| Merging | 48850754 | 1.000 |
| Demultiplexing | 44160913 | 0.904 |
| Length Trimming | 41141042 | 0.932 |
| **Lactose environment** | | |
| Merging | 54006855 | 1.000 |
| Demultiplexing | 46623230 | 0.863 |
| Length Trimming | 44828702 | 0.962 |

**Table S5.**

Sequences of Oligopool1 (First [$H_{G+L}$] category variants). All sequences have a 5′ phosphate modification, i.e., a phosphate group is attached to the 5′ end of each strand. The 21 bp *lac* operator sequence is capitalized.

| No. | Oligonucleotide sequences |
|---|---|
| 1 | catagctgtttcctgtgtgaAATTGTTATCCGCTTCAGCCCccacacaacatacgagccggaagcataaagtgtaaagcctgg |
| 2 | catagctgtttcctgtgtgaAATTGTTATCCGCGGAGTTCAccacacaacatacgagccggaagcataaagtgtaaagcctgg |
| 3 | catagctgtttcctgtgtgaAATTGTTATCCGGCCGCAGTAccacacaacatacgagccggaagcataaagtgtaaagcctgg |
| 4 | catagctgtttcctgtgtgaAATTGTTATCCGAACTTTACAccacacaacatacgagccggaagcataaagtgtaaagcctgg |
| 5 | catagctgtttcctgtgtgaAATTGTTATCCGCATATGGCGccacacaacatacgagccggaagcataaagtgtaaagcctgg |
| 6 | catagctgtttcctgtgtgaAATTGTTATCCGCCTGAACAGccacacaacatacgagccggaagcataaagtgtaaagcctgg |
| 7 | catagctgtttcctgtgtgaAATTGTTATCCGGTTGAATGAccacacaacatacgagccggaagcataaagtgtaaagcctgg |
| 8 | catagctgtttcctgtgtgaAATTGTTATCCGCTCACAAGTccacacaacatacgagccggaagcataaagtgtaaagcctgg |
| 9 | catagctgtttcctgtgtgaAATTGTTATCCGGGTGACCCAccacacaacatacgagccggaagcataaagtgtaaagcctgg |
| 10 | catagctgtttcctgtgtgaAATTGTTATCCGTTCCCTCTGccacacaacatacgagccggaagcataaagtgtaaagcctgg |
| 11 | catagctgtttcctgtgtgaAATTGTTATCCGGCATTACCCccacacaacatacgagccggaagcataaagtgtaaagcctgg |
| 12 | catagctgtttcctgtgtgaAATTGTTATCCGCACTGGCCGccacacaacatacgagccggaagcataaagtgtaaagcctgg |
| 13 | catagctgtttcctgtgtgaAATTGTTATCCGTCCCCTCGGccacacaacatacgagccggaagcataaagtgtaaagcctgg |
| 14 | catagctgtttcctgtgtgaAATTGTTATCCGCAAGAGCCGccacacaacatacgagccggaagcataaagtgtaaagcctgg |
| 15 | catagctgtttcctgtgtgaAATTGTTATCCGGAGGTTGCGccacacaacatacgagccggaagcataaagtgtaaagcctgg |
| 16 | catagctgtttcctgtgtgaAATTGTTATCCGTATCCGTACccacacaacatacgagccggaagcataaagtgtaaagcctgg |
| 17 | catagctgtttcctgtgtgaAATTGTTATCCGCGAGAAGGGccacacaacatacgagccggaagcataaagtgtaaagcctgg |
| 18 | catagctgtttcctgtgtgaAATTGTTATCCGCAAACCGAAccacacaacatacgagccggaagcataaagtgtaaagcctgg |
| 19 | catagctgtttcctgtgtgaAATTGTTATCCGACATCCCCTccacacaacatacgagccggaagcataaagtgtaaagcctgg |
| 20 | catagctgtttcctgtgtgaAATTGTTATCCGAACCACGCAccacacaacatacgagccggaagcataaagtgtaaagcctgg |
| 21 | catagctgtttcctgtgtgaAATTGTTATCCGGCTTAATCTccacacaacatacgagccggaagcataaagtgtaaagcctgg |
| 22 | catagctgtttcctgtgtgaAATTGTTATCCGAGGGGATCAccacacaacatacgagccggaagcataaagtgtaaagcctgg |
| 23 | catagctgtttcctgtgtgaAATTGTTATCCGAGAGAACAAccacacaacatacgagccggaagcataaagtgtaaagcctgg |
| 24 | catagctgtttcctgtgtgaAATTGTTATCCGTGTTCCCGTccacacaacatacgagccggaagcataaagtgtaaagcctgg |
| 25 | catagctgtttcctgtgtgaAATTGTTATCCGTGGGCATCAccacacaacatacgagccggaagcataaagtgtaaagcctgg |
| 26 | catagctgtttcctgtgtgaAATTGTTATCCGCGGCTAGGTccacacaacatacgagccggaagcataaagtgtaaagcctgg |

| | |
|---|---|
| 27 | catagctgtttcctgtgtgaAATTGTTATCCGTCCAGTCCTccacacaacatacgagccggaagcataaagtgtaaagcctgg |
| 28 | catagctgtttcctgtgtgaAATTGTTATCCGCGTTTCATAccacacaacatacgagccggaagcataaagtgtaaagcctgg |
| 29 | catagctgtttcctgtgtgaAATTGTTATCCGACGCGATCGccacacaacatacgagccggaagcataaagtgtaaagcctgg |
| 30 | catagctgtttcctgtgtgaAATTGTTATCCGTTCGACGGGccacacaacatacgagccggaagcataaagtgtaaagcctgg |
| 31 | catagctgtttcctgtgtgaAATTGTTATCCGCAGTGAACAccacacaacatacgagccggaagcataaagtgtaaagcctgg |
| 32 | catagctgtttcctgtgtgaAATTGTTATCCGCGAAGAACGccacacaacatacgagccggaagcataaagtgtaaagcctgg |
| 33 | catagctgtttcctgtgtgaAATTGTTATCCGGCACTGCGCccacacaacatacgagccggaagcataaagtgtaaagcctgg |
| 34 | catagctgtttcctgtgtgaAATTGTTATCCGTTGGCATACccacacaacatacgagccggaagcataaagtgtaaagcctgg |
| 35 | catagctgtttcctgtgtgaAATTGTTATCCGCGGCAGGTTccacacaacatacgagccggaagcataaagtgtaaagcctgg |
| 36 | catagctgtttcctgtgtgaAATTGTTATCCGCATGCTCAGccacacaacatacgagccggaagcataaagtgtaaagcctgg |
| 37 | catagctgtttcctgtgtgaAATTGTTATCCGCGGTCTTCGccacacaacatacgagccggaagcataaagtgtaaagcctgg |
| 38 | catagctgtttcctgtgtgaAATTGTTATCCGTACCAACCGccacacaacatacgagccggaagcataaagtgtaaagcctgg |
| 39 | catagctgtttcctgtgtgaAATTGTTATCCGTGCGGTATAccacacaacatacgagccggaagcataaagtgtaaagcctgg |
| 40 | catagctgtttcctgtgtgaAATTGTTATCCGGAAAGACACccacacaacatacgagccggaagcataaagtgtaaagcctgg |
| 41 | catagctgtttcctgtgtgaAATTGTTATCCGATCATTGACccacacaacatacgagccggaagcataaagtgtaaagcctgg |
| 42 | catagctgtttcctgtgtgaAATTGTTATCCGACCAAGGACccacacaacatacgagccggaagcataaagtgtaaagcctgg |
| 43 | catagctgtttcctgtgtgaAATTGTTATCCGACCAGGATGccacacaacatacgagccggaagcataaagtgtaaagcctgg |
| 44 | catagctgtttcctgtgtgaAATTGTTATCCGGGCGTGCGCccacacaacatacgagccggaagcataaagtgtaaagcctgg |
| 45 | catagctgtttcctgtgtgaAATTGTTATCCGGGGGAAGCAccacacaacatacgagccggaagcataaagtgtaaagcctgg |
| 46 | catagctgtttcctgtgtgaAATTGTTATCCGCACAGCAATccacacaacatacgagccggaagcataaagtgtaaagcctgg |
| 47 | catagctgtttcctgtgtgaAATTGTTATCCGCTCACCGGCccacacaacatacgagccggaagcataaagtgtaaagcctgg |
| 48 | catagctgtttcctgtgtgaAATTGTTATCCGAAGCTCACGccacacaacatacgagccggaagcataaagtgtaaagcctgg |
| 49 | catagctgtttcctgtgtgaAATTGTTATCCGATCGATCTCccacacaacatacgagccggaagcataaagtgtaaagcctgg |
| 50 | catagctgtttcctgtgtgaAATTGTTATCCGGTCTCCTGGccacacaacatacgagccggaagcataaagtgtaaagcctgg |

**Table S6.**

Sequences of Oligopool2 (Second [$H_G$] and third [$H_L$] category variants). All sequences have a 5′ phosphate modification, i.e., a phosphate group is attached to the 5′ end of each strand. The 21 bp *lac* operator sequence is capitalized.

| No. | Oligonucleotide sequences |
|---|---|
| 1 | catagctgtttcctgtgtgaAATTGTTATCCGCCTGGGGTGccacacaacatacgagccggaagcataaagtgtaaagcctgg |
| 2 | catagctgtttcctgtgtgaAATTGTTATCCGGTCTAAGCCccacacaacatacgagccggaagcataaagtgtaaagcctgg |
| 3 | catagctgtttcctgtgtgaAATTGTTATCCGAGTTAGTTAccacacaacatacgagccggaagcataaagtgtaaagcctgg |
| 4 | catagctgtttcctgtgtgaAATTGTTATCCGTCGAGACGCccacacaacatacgagccggaagcataaagtgtaaagcctgg |
| 5 | catagctgtttcctgtgtgaAATTGTTATCCGCCGGGGTGTccacacaacatacgagccggaagcataaagtgtaaagcctgg |
| 6 | catagctgtttcctgtgtgaAATTGTTATCCGGAATGACCCccacacaacatacgagccggaagcataaagtgtaaagcctgg |
| 7 | catagctgtttcctgtgtgaAATTGTTATCCGGCATTTGGGccacacaacatacgagccggaagcataaagtgtaaagcctgg |
| 8 | catagctgtttcctgtgtgaAATTGTTATCCGGGATATCTCccacacaacatacgagccggaagcataaagtgtaaagcctgg |
| 9 | catagctgtttcctgtgtgaAATTGTTATCCGGTTTCTTTAccacacaacatacgagccggaagcataaagtgtaaagcctgg |
| 10 | catagctgtttcctgtgtgaAATTGTTATCCGGACTCTGGTccacacaacatacgagccggaagcataaagtgtaaagcctgg |
| 11 | catagctgtttcctgtgtgaAATTGTTATCCGGGGACTCTGccacacaacatacgagccggaagcataaagtgtaaagcctgg |
| 12 | catagctgtttcctgtgtgaAATTGTTATCCGCACATATGCccacacaacatacgagccggaagcataaagtgtaaagcctgg |
| 13 | catagctgtttcctgtgtgaAATTGTTATCCGACGTAGCGGccacacaacatacgagccggaagcataaagtgtaaagcctgg |
| 14 | catagctgtttcctgtgtgaAATTGTTATCCGCCGAGGTCCccacacaacatacgagccggaagcataaagtgtaaagcctgg |
| 15 | catagctgtttcctgtgtgaAATTGTTATCCGTGACGAGCCccacacaacatacgagccggaagcataaagtgtaaagcctgg |
| 16 | catagctgtttcctgtgtgaAATTGTTATCCGGGCACCTGCccacacaacatacgagccggaagcataaagtgtaaagcctgg |
| 17 | catagctgtttcctgtgtgaAATTGTTATCCGGATAAAAAGccacacaacatacgagccggaagcataaagtgtaaagcctgg |
| 18 | catagctgtttcctgtgtgaAATTGTTATCCGAACCAACCCccacacaacatacgagccggaagcataaagtgtaaagcctgg |
| 19 | catagctgtttcctgtgtgaAATTGTTATCCGTACATTATGccacacaacatacgagccggaagcataaagtgtaaagcctgg |
| 20 | catagctgtttcctgtgtgaAATTGTTATCCGACAAATGCAccacacaacatacgagccggaagcataaagtgtaaagcctgg |
| 21 | catagctgtttcctgtgtgaAATTGTTATCCGGTAAACCCCccacacaacatacgagccggaagcataaagtgtaaagcctgg |
| 22 | catagctgtttcctgtgtgaAATTGTTATCCGCTTAGGCCCccacacaacatacgagccggaagcataaagtgtaaagcctgg |
| 23 | catagctgtttcctgtgtgaAATTGTTATCCGCGCCCTGCCccacacaacatacgagccggaagcataaagtgtaaagcctgg |
| 24 | catagctgtttcctgtgtgaAATTGTTATCCGCGCCACATGccacacaacatacgagccggaagcataaagtgtaaagcctgg |
| 25 | catagctgtttcctgtgtgaAATTGTTATCCGCAGGAGCCGccacacaacatacgagccggaagcataaagtgtaaagcctgg |
| 26 | catagctgtttcctgtgtgaAATTGTTATCCGAACTAGCACccacacaacatacgagccggaagcataaagtgtaaagcctgg |

| | |
|---|---|
| 27 | catagctgtttcctgtgtgaAATTGTTATCCGTCTGATCTAccacacaacatacgagccggaagcataaagtgtaaagcctgg |
| 28 | catagctgtttcctgtgtgaAATTGTTATCCGAAGTTTCGTccacacaacatacgagccggaagcataaagtgtaaagcctgg |
| 29 | catagctgtttcctgtgtgaAATTGTTATCCGAATACCATAccacacaacatacgagccggaagcataaagtgtaaagcctgg |
| 30 | catagctgtttcctgtgtgaAATTGTTATCCGGGAGTGCGAccacacaacatacgagccggaagcataaagtgtaaagcctgg |
| 31 | catagctgtttcctgtgtgaAATTGTTATCCGACGATCAGTccacacaacatacgagccggaagcataaagtgtaaagcctgg |
| 32 | catagctgtttcctgtgtgaAATTGTTATCCGTGGCACCCGccacacaacatacgagccggaagcataaagtgtaaagcctgg |
| 33 | catagctgtttcctgtgtgaAATTGTTATCCGCTGATTCTTccacacaacatacgagccggaagcataaagtgtaaagcctgg |
| 34 | catagctgtttcctgtgtgaAATTGTTATCCGGATCACCCGccacacaacatacgagccggaagcataaagtgtaaagcctgg |
| 35 | catagctgtttcctgtgtgaAATTGTTATCCGAGGGACGGGccacacaacatacgagccggaagcataaagtgtaaagcctgg |
| 36 | catagctgtttcctgtgtgaAATTGTTATCCGTGCAGTACCccacacaacatacgagccggaagcataaagtgtaaagcctgg |
| 37 | catagctgtttcctgtgtgaAATTGTTATCCGCAAATGGTGccacacaacatacgagccggaagcataaagtgtaaagcctgg |
| 38 | catagctgtttcctgtgtgaAATTGTTATCCGAATCATCAGccacacaacatacgagccggaagcataaagtgtaaagcctgg |
| 39 | catagctgtttcctgtgtgaAATTGTTATCCGGCCGGCTATccacacaacatacgagccggaagcataaagtgtaaagcctgg |
| 40 | catagctgtttcctgtgtgaAATTGTTATCCGCCCCGTAGCccacacaacatacgagccggaagcataaagtgtaaagcctgg |
| 41 | catagctgtttcctgtgtgaAATTGTTATCCGGATCTCTGAccacacaacatacgagccggaagcataaagtgtaaagcctgg |
| 42 | catagctgtttcctgtgtgaAATTGTTATCCGCTCAGCGTGccacacaacatacgagccggaagcataaagtgtaaagcctgg |
| 43 | catagctgtttcctgtgtgaAATTGTTATCCGCGACCTCCTccacacaacatacgagccggaagcataaagtgtaaagcctgg |
| 44 | catagctgtttcctgtgtgaAATTGTTATCCGAAGTGGGCAccacacaacatacgagccggaagcataaagtgtaaagcctgg |
| 45 | catagctgtttcctgtgtgaAATTGTTATCCGATGTAGATTccacacaacatacgagccggaagcataaagtgtaaagcctgg |
| 46 | catagctgtttcctgtgtgaAATTGTTATCCGCGTACCTGAccacacaacatacgagccggaagcataaagtgtaaagcctgg |
| 47 | catagctgtttcctgtgtgaAATTGTTATCCGCCTACACCCccacacaacatacgagccggaagcataaagtgtaaagcctgg |
| 48 | catagctgtttcctgtgtgaAATTGTTATCCGAACCCTGCAccacacaacatacgagccggaagcataaagtgtaaagcctgg |
| 49 | catagctgtttcctgtgtgaAATTGTTATCCGGCCTTAGGCccacacaacatacgagccggaagcataaagtgtaaagcctgg |
| 50 | catagctgtttcctgtgtgaAATTGTTATCCGTCCGGGCGTccacacaacatacgagccggaagcataaagtgtaaagcctgg |
| 51 | catagctgtttcctgtgtgaAATTGTTATCCGGTAGGCCAAccacacaacatacgagccggaagcataaagtgtaaagcctgg |
| 52 | catagctgtttcctgtgtgaAATTGTTATCCGTCCCGGGTAccacacaacatacgagccggaagcataaagtgtaaagcctgg |

**Table S7.**

Sequences of Oligopool3 (Fourth category [$L_{G+L}$] variants). All sequences have a 5′ phosphate modification, i.e., a phosphate group is attached to the 5′ end of each strand. The 21 bp *lac* operator sequence is capitalized.

| No. | Oligonucleotide sequences |
|---|---|
| 1 | catagctgtttcctgtgtgaAATTGTTATCCGTATAAGCCTccacacaacatacgagccggaagcataaagtgtaaagcctgg |
| 2 | catagctgtttcctgtgtgaAATTGTTATCCGGTGGGCCAGccacacaacatacgagccggaagcataaagtgtaaagcctgg |
| 3 | catagctgtttcctgtgtgaAATTGTTATCCGTGTACCGTAccacacaacatacgagccggaagcataaagtgtaaagcctgg |
| 4 | catagctgtttcctgtgtgaAATTGTTATCCGGTGCAGGAGccacacaacatacgagccggaagcataaagtgtaaagcctgg |
| 5 | catagctgtttcctgtgtgaAATTGTTATCCGCAACACGAGccacacaacatacgagccggaagcataaagtgtaaagcctgg |
| 6 | catagctgtttcctgtgtgaAATTGTTATCCGACGTCTTCCccacacaacatacgagccggaagcataaagtgtaaagcctgg |
| 7 | catagctgtttcctgtgtgaAATTGTTATCCGGCTATGGTCccacacaacatacgagccggaagcataaagtgtaaagcctgg |
| 8 | catagctgtttcctgtgtgaAATTGTTATCCGACAAACCTCccacacaacatacgagccggaagcataaagtgtaaagcctgg |
| 9 | catagctgtttcctgtgtgaAATTGTTATCCGTTCGGGCGGccacacaacatacgagccggaagcataaagtgtaaagcctgg |
| 10 | catagctgtttcctgtgtgaAATTGTTATCCGAATCTATTCccacacaacatacgagccggaagcataaagtgtaaagcctgg |
| 11 | catagctgtttcctgtgtgaAATTGTTATCCGCTGCGGGTTccacacaacatacgagccggaagcataaagtgtaaagcctgg |
| 12 | catagctgtttcctgtgtgaAATTGTTATCCGCCCATGCCCccacacaacatacgagccggaagcataaagtgtaaagcctgg |
| 13 | catagctgtttcctgtgtgaAATTGTTATCCGGTCAGCGCAccacacaacatacgagccggaagcataaagtgtaaagcctgg |
| 14 | catagctgtttcctgtgtgaAATTGTTATCCGGGAAATTGCccacacaacatacgagccggaagcataaagtgtaaagcctgg |
| 15 | catagctgtttcctgtgtgaAATTGTTATCCGCGTGTATGAccacacaacatacgagccggaagcataaagtgtaaagcctgg |
| 16 | catagctgtttcctgtgtgaAATTGTTATCCGTTGTTTTAGccacacaacatacgagccggaagcataaagtgtaaagcctgg |
| 17 | catagctgtttcctgtgtgaAATTGTTATCCGCGAGTGGCAccacacaacatacgagccggaagcataaagtgtaaagcctgg |
| 18 | catagctgtttcctgtgtgaAATTGTTATCCGGGGGAGAGCccacacaacatacgagccggaagcataaagtgtaaagcctgg |
| 19 | catagctgtttcctgtgtgaAATTGTTATCCGCGTCCGACGccacacaacatacgagccggaagcataaagtgtaaagcctgg |
| 20 | catagctgtttcctgtgtgaAATTGTTATCCGATCGGTATGccacacaacatacgagccggaagcataaagtgtaaagcctgg |
| 21 | catagctgtttcctgtgtgaAATTGTTATCCGATCGTGGAGccacacaacatacgagccggaagcataaagtgtaaagcctgg |
| 22 | catagctgtttcctgtgtgaAATTGTTATCCGCATCGCGTGccacacaacatacgagccggaagcataaagtgtaaagcctgg |
| 23 | catagctgtttcctgtgtgaAATTGTTATCCGGCCACGCCGccacacaacatacgagccggaagcataaagtgtaaagcctgg |
| 24 | catagctgtttcctgtgtgaAATTGTTATCCGGAAGAGGCAccacacaacatacgagccggaagcataaagtgtaaagcctgg |
| 25 | catagctgtttcctgtgtgaAATTGTTATCCGAGCCTCATAccacacaacatacgagccggaagcataaagtgtaaagcctgg |
| 26 | catagctgtttcctgtgtgaAATTGTTATCCGATTCTCGGAccacacaacatacgagccggaagcataaagtgtaaagcctgg |

| | |
|---|---|
| 27 | catagctgtttcctgtgtgaAATTGTTATCCGATTCAGAAGccacacaacatacgagccggaagcataaagtgtaaagcctgg |
| 28 | catagctgtttcctgtgtgaAATTGTTATCCGACGTCGCATccacacaacatacgagccggaagcataaagtgtaaagcctgg |
| 29 | catagctgtttcctgtgtgaAATTGTTATCCGATGAAATTAccacacaacatacgagccggaagcataaagtgtaaagcctgg |
| 30 | catagctgtttcctgtgtgaAATTGTTATCCGGTTGAGGGTccacacaacatacgagccggaagcataaagtgtaaagcctgg |
| 31 | catagctgtttcctgtgtgaAATTGTTATCCGATGGGAAATccacacaacatacgagccggaagcataaagtgtaaagcctgg |
| 32 | catagctgtttcctgtgtgaAATTGTTATCCGGCTCTTGTAccacacaacatacgagccggaagcataaagtgtaaagcctgg |
| 33 | catagctgtttcctgtgtgaAATTGTTATCCGGACTTGATAccacacaacatacgagccggaagcataaagtgtaaagcctgg |
| 34 | catagctgtttcctgtgtgaAATTGTTATCCGAGTTCTATAccacacaacatacgagccggaagcataaagtgtaaagcctgg |
| 35 | catagctgtttcctgtgtgaAATTGTTATCCGCGTAGAGGAccacacaacatacgagccggaagcataaagtgtaaagcctgg |
| 36 | catagctgtttcctgtgtgaAATTGTTATCCGGTCAGAGTGccacacaacatacgagccggaagcataaagtgtaaagcctgg |
| 37 | catagctgtttcctgtgtgaAATTGTTATCCGATGACAGCCccacacaacatacgagccggaagcataaagtgtaaagcctgg |
| 38 | catagctgtttcctgtgtgaAATTGTTATCCGTACATTAAGccacacaacatacgagccggaagcataaagtgtaaagcctgg |
| 39 | catagctgtttcctgtgtgaAATTGTTATCCGGCGCTTGGTccacacaacatacgagccggaagcataaagtgtaaagcctgg |
| 40 | catagctgtttcctgtgtgaAATTGTTATCCGTCAGTGTCCccacacaacatacgagccggaagcataaagtgtaaagcctgg |
| 41 | catagctgtttcctgtgtgaAATTGTTATCCGGTAGTAAAGccacacaacatacgagccggaagcataaagtgtaaagcctgg |
| 42 | catagctgtttcctgtgtgaAATTGTTATCCGAGAACTTCGccacacaacatacgagccggaagcataaagtgtaaagcctgg |
| 43 | catagctgtttcctgtgtgaAATTGTTATCCGTCTATCAATccacacaacatacgagccggaagcataaagtgtaaagcctgg |
| 44 | catagctgtttcctgtgtgaAATTGTTATCCGAACTGTTTTccacacaacatacgagccggaagcataaagtgtaaagcctgg |
| 45 | catagctgtttcctgtgtgaAATTGTTATCCGCTCAGACTAccacacaacatacgagccggaagcataaagtgtaaagcctgg |
| 46 | catagctgtttcctgtgtgaAATTGTTATCCGTTGTTTCAGccacacaacatacgagccggaagcataaagtgtaaagcctgg |
| 47 | catagctgtttcctgtgtgaAATTGTTATCCGCTGTAATTAccacacaacatacgagccggaagcataaagtgtaaagcctgg |
| 48 | catagctgtttcctgtgtgaAATTGTTATCCGGTTCGTTATccacacaacatacgagccggaagcataaagtgtaaagcctgg |
| 49 | catagctgtttcctgtgtgaAATTGTTATCCGGGTGAGAAAccacacaacatacgagccggaagcataaagtgtaaagcctgg |
| 50 | catagctgtttcctgtgtgaAATTGTTATCCGTGTCTTCGAccacacaacatacgagccggaagcataaagtgtaaagcctgg |

**Table S8.**

Frequency of epistasis types from network motifs

| Motifs and Types | Frequency |
|---|---|
| **Glycerol environment** | |
| No of motifs[1] | 7040163 |
| Magnitude epistasis or additivity | 2341099 (33.2%) |
| Simple sign epistasis | 2355806 (33.5%) |
| Reciprocal sign epistasis | 2343258 (33.3%) |
| **Lactose environment** | |
| No of motifs[1] | 2837859 |
| Magnitude epistasis or additivity | 944666 (33.3%) |
| Simple sign epistasis | 944557 (33.3%) |
| Reciprocal sign epistasis | 948636 (33.4%) |

[1]A motif is a quadruplet of genotypes connected by edges, i.e., a focal genotype (ab) and a double mutant (AB) connected via two single mutants (Ab and aB)

### Supplementary figures

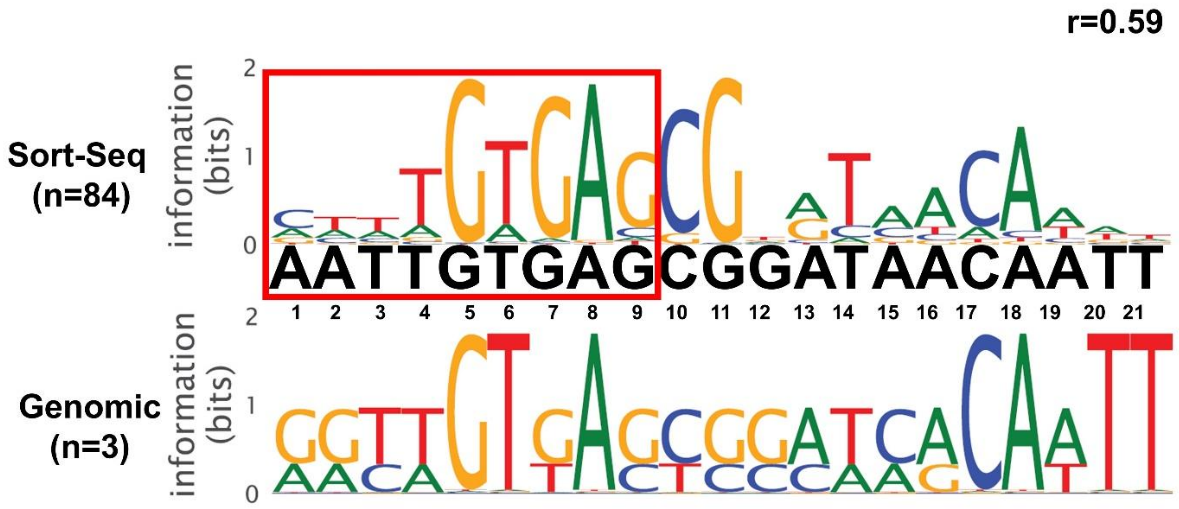

**Figure S1. DNA Sequence logo obtained by a previous study.** Belliveau et al. used the Sort-Seq technique to study the *lac* promoter's regulatory landscape by generating a library of double-stranded DNA sequences with all possible single-base substitutions within two LacI repressor and CRP binding sites (49). Sort-Seq experiments were performed with cells grown in M9 minimal media with 0.5% glucose. Regulatory binding sites were identified by calculating the average expression shift caused by mutations at each position. The Sort-Seq data was used to generate energy matrices by quantifying these shifts, which were then used to calculate the probabilities of each nucleotide at a position based on their contribution to binding strength. These probabilities were used to build the position weight matrix (PWM). From the PWM, the average information content at each position along the binding site was calculated, which was then used to generate the sequence logo. The height of each letter in the logo represents the nucleotide frequency at each position, scaled by its information content, calculated using Shannon entropy. The resulting logo reflects binding strength contributions inferred from the Sort-Seq data. The figure shows the corresponding DNA sequence logo of the *lac* operator obtained from the study (49) with comparisons made between LacI Sort-Seq data and binding site sequences from RegulonDB. *n* denotes the number of mutants and genomic binding sites used to construct the sequence logo. Pearson correlation coefficients were calculated using the position weight matrices from the Sort-Seq and genomic matrices. The red box highlights the positions of the *lac* operator used for mutagenesis in the current study.

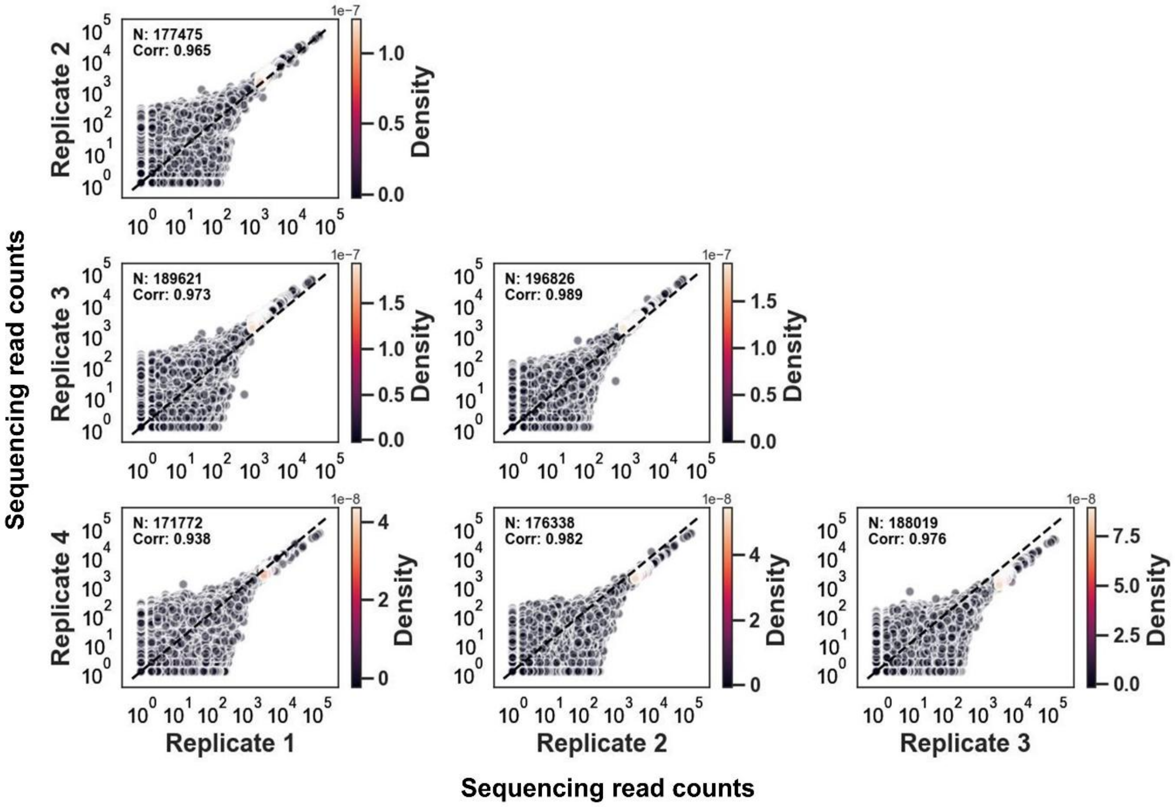

**Figure S2. Correlation of read coverage in glucose (pre-selection) between four replicates** of the *lac* operator mutant library. Note the logarithmic scale in all panels. N: Number of data points. Corr: correlation coefficient between the respective replicates.

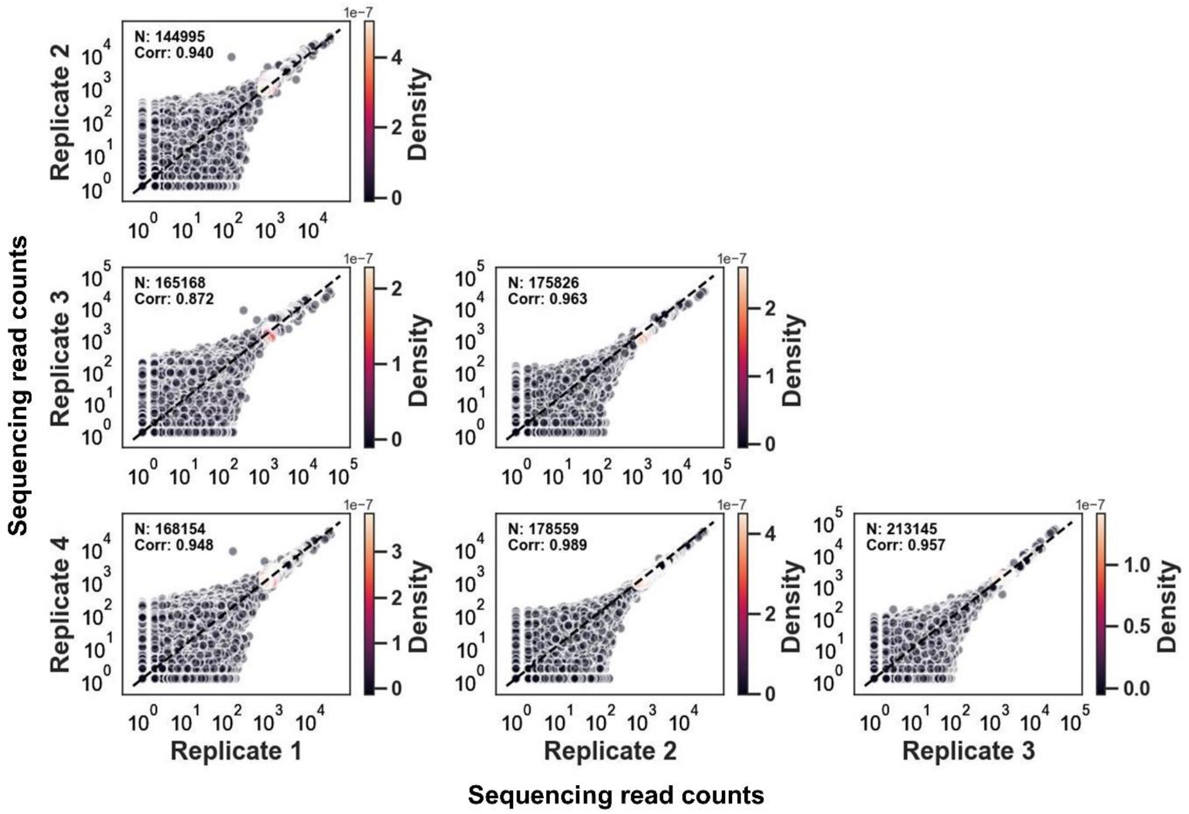

**Figure S3. Correlation of read coverage in glycerol (post selection) between four replicates** of the *lac* operator mutant library. Note the logarithmic scale in all panels. N: Number of data points. Corr: correlation coefficient between the respective replicates.

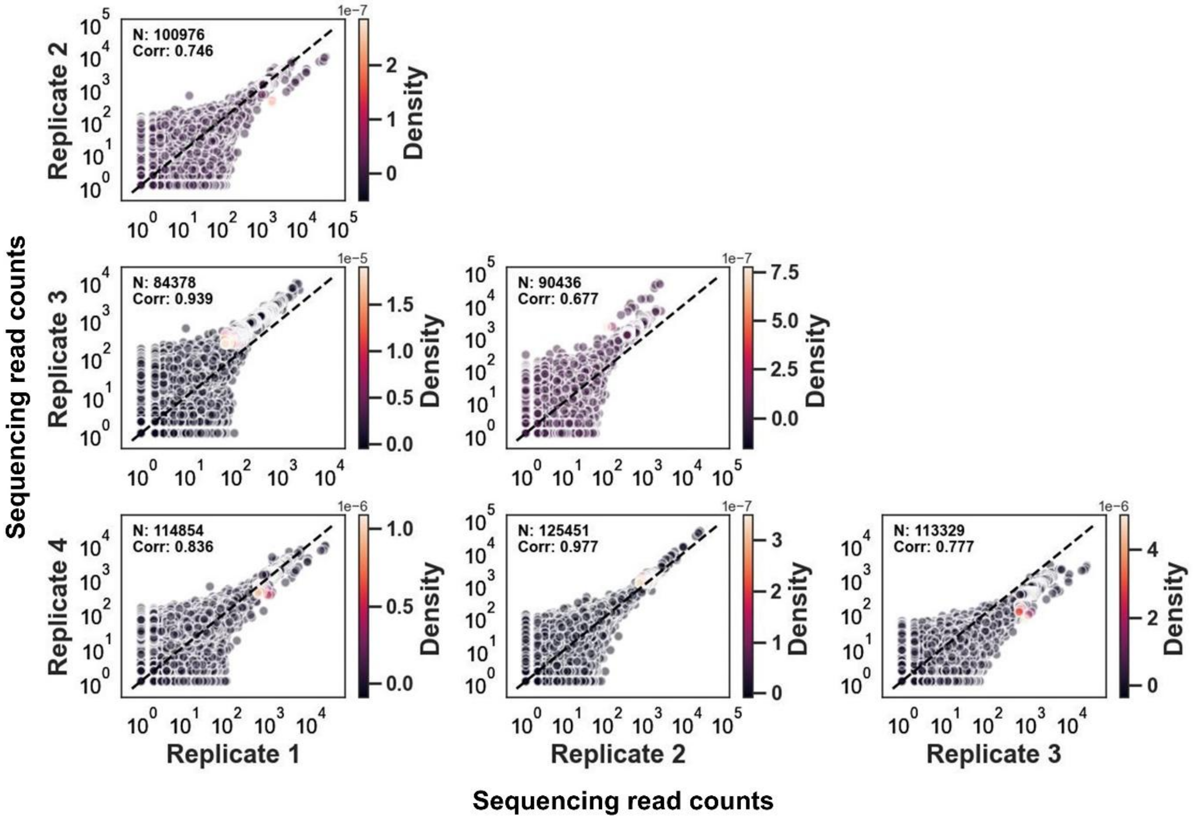

**Figure S4. Correlation of read coverage in lactose (post selection) between four replicates** of the *lac* operator mutant library. Note the logarithmic scale in all panels. N: Number of data points. Corr: correlation coefficient between the respective replicates.

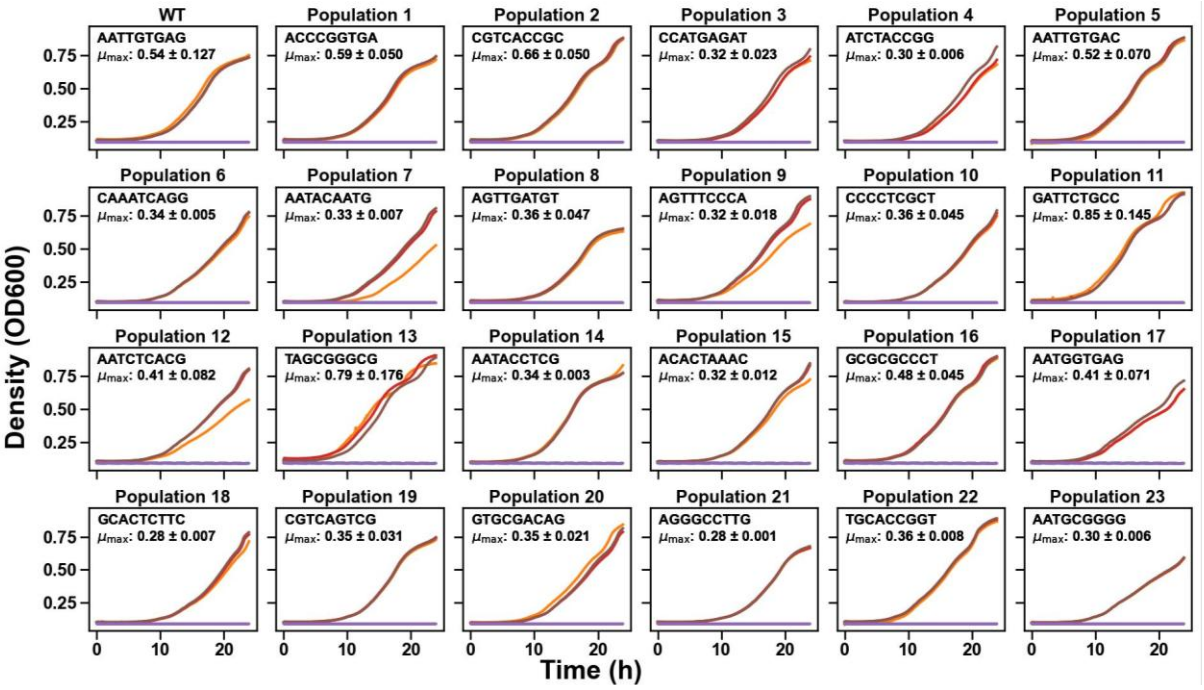

**Figure S5. Growth curves of 24 *E. coli* populations harboring 24 different *lac* operator variants in individual cultures grown in M9 medium with 10% glycerol**. Each panel shows the increase in optical density ($OD_{600}$, vertical axis) of one *lac* operator variant in M9 medium containing 10% glycerol (and 0.2% casamino acid) over a 24-hour period (horizontal axis). Each panel displays the growth curves for the WT and one of the 23 isolated clones (the *lac* operator sequence of the respective clone is shown within each panel), with different colors representing $OD_{600}$ measurements of three replicate cultures for each variant. The dashed purple line in each panel represents a blank media control without cells. We used these growth curves to estimate the maximum growth rate, which is indicated in the upper left of each panel as the mean± one standard error (Methods).

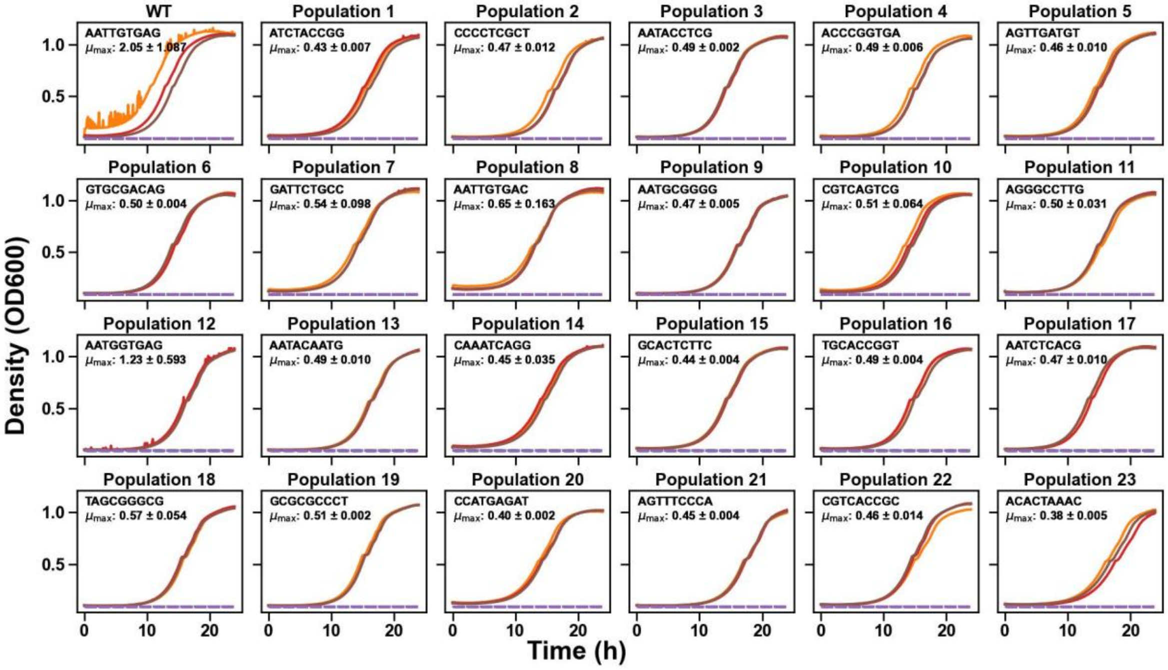

**Figure S6. Growth curves of 24 *E. coli* populations harboring 24 different *lac* operator variants in individual cultures grown in M9 medium with 1 mM lactose**. Each panel shows the increase in optical density ($OD_{600}$, vertical axis) of one *lac* operator variant in M9 medium containing 1 mM lactose (and 0.2% casamino acid) over a 24-hour period (horizontal axis). Each panel displays the growth curves for the WT and one of the 23 isolated clones (the *lac* operator sequence of the respective clone is shown within each panel), with different colors representing $OD_{600}$ measurements of three replicate cultures for each variant. The dashed purple line in each panel represents a blank media control without cells. We used these growth curves to estimate the maximum growth rate, which is indicated in the upper left of each panel as the mean±one standard error (Methods).

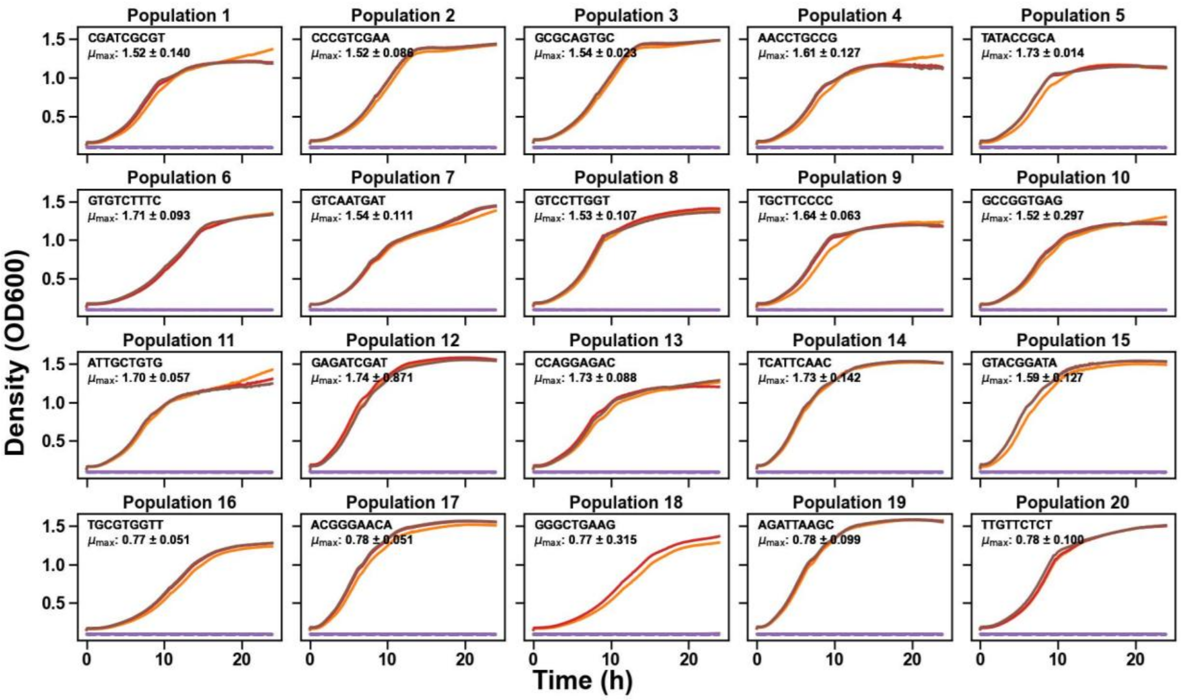

**Figure S7. Growth curves of 20 *E. coli* populations harboring 20 different *lac* operator variants from $H_{G+L}$ category in individual cultures grown in M9 medium with 10% glycerol**. Each panel shows the increase in optical density ($OD_{600}$, vertical axis) of one *lac* operator variant in M9 medium containing 10% glycerol (and 0.2% casamino acid) over a 24-hour period (horizontal axis). Each panel displays the growth curves for one of the 20 isolated clones (the *lac* operator sequence of the respective clone is shown within each panel) belonging to $H_{G+L}$ category (high fitness in both glycerol and lactose), with different colors representing $OD_{600}$ measurements of three replicate cultures for each variant. The dashed purple line in each panel represents a blank media control without cells. We used these growth curves to estimate the maximum growth rate, which is indicated in the upper left of each panel as the mean±one standard error (Methods).

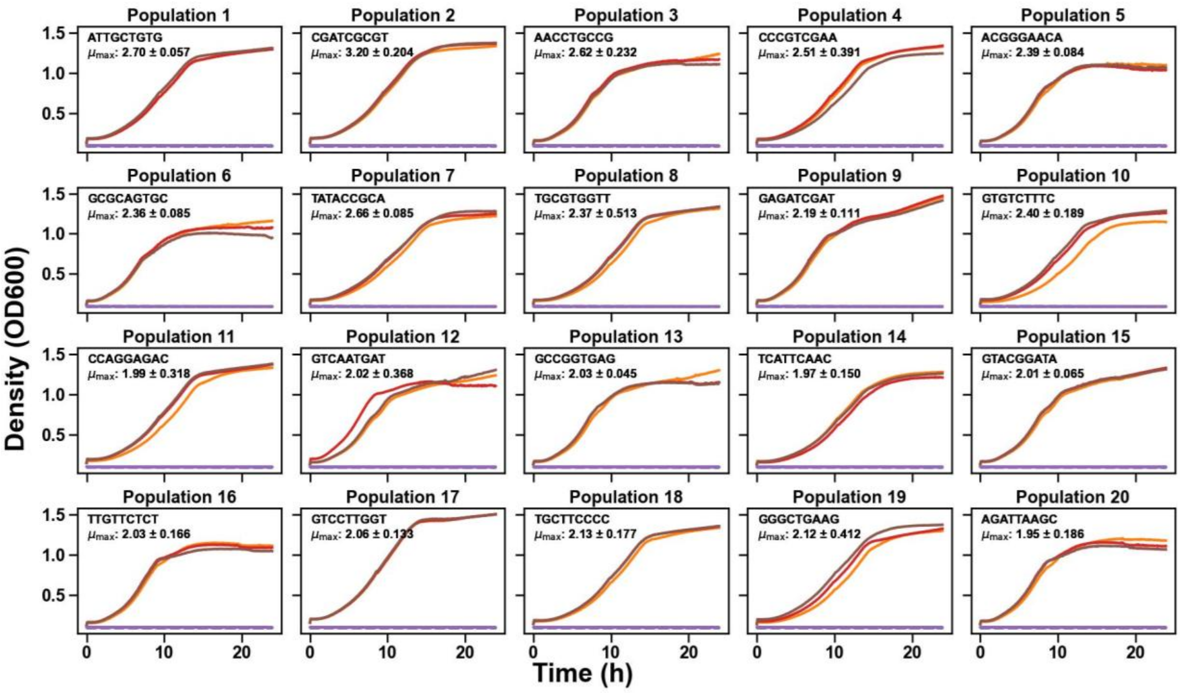

**Figure S8. Growth curves of 20 *E. coli* populations harboring 20 different *lac* operator variants from $H_{G+L}$ category in individual cultures grown in M9 medium with 1 mM lactose**. Each panel shows the increase in optical density ($OD_{600}$, vertical axis) of one *lac* operator variant in M9 medium containing 1 mM lactose (and 0.2% casamino acid) over a 24-hour period (horizontal axis). Each panel displays the growth curves for one of the 20 isolated clones (the *lac* operator sequence of the respective clone is shown within each panel) belonging to $H_{G+L}$ category (high fitness in both glycerol and lactose), with different colors representing $OD_{600}$ measurements of three replicate cultures for each variant. The dashed purple line in each panel represents a blank media control without cells. We used these growth curves to estimate the maximum growth rate, which is indicated in the upper left of each panel as the mean±one standard error (Methods).

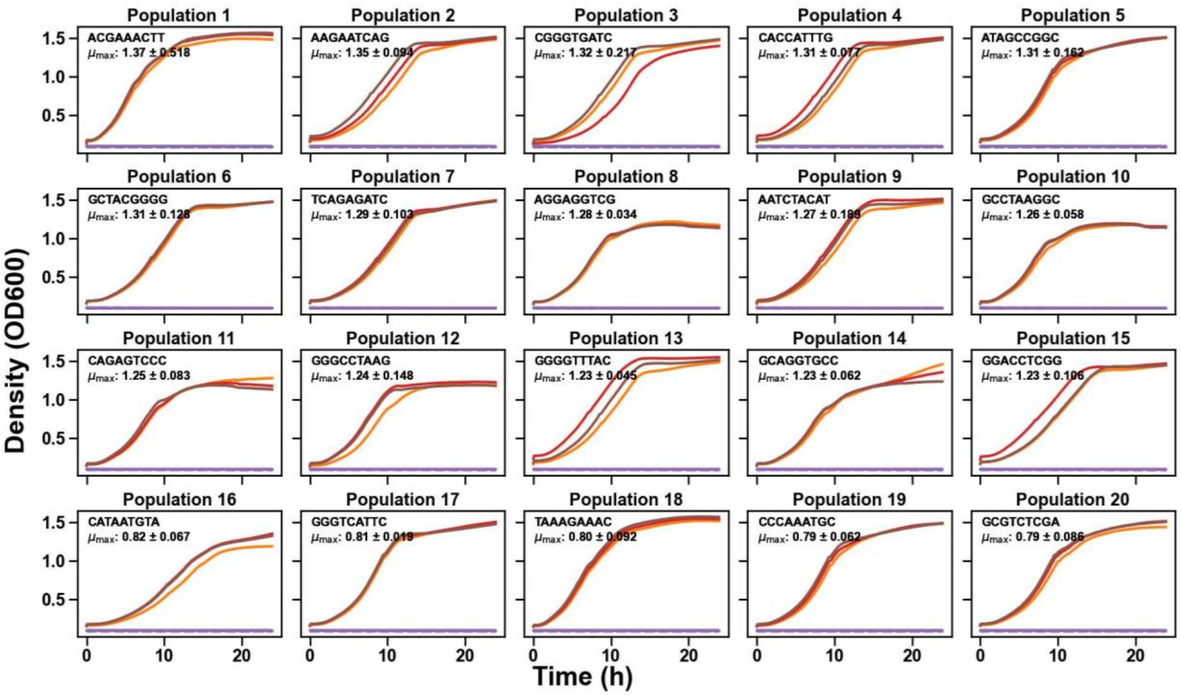

**Figure S9. Growth curves of 20 *E. coli* populations harboring 20 different *lac* operator variants from $H_G$ category in individual cultures grown in M9 medium with 10% glycerol.** Each panel shows the increase in optical density ($OD_{600}$, vertical axis) of one *lac* operator variant in M9 medium containing 10% glycerol (and 0.2% casamino acid) over a 24-hour period (horizontal axis). Each panel displays the growth curves for one of the 20 isolated clones (the *lac* operator sequence of the respective clone is shown within each panel) belonging to $H_G$ category (high fitness in glycerol and low fitness in lactose), with different colors representing $OD_{600}$ measurements of three replicate cultures for each variant. The dashed purple line in each panel represents a blank media control without cells. We used these growth curves to estimate the maximum growth rate, which is indicated in the upper left of each panel as the mean±one standard error (Methods).

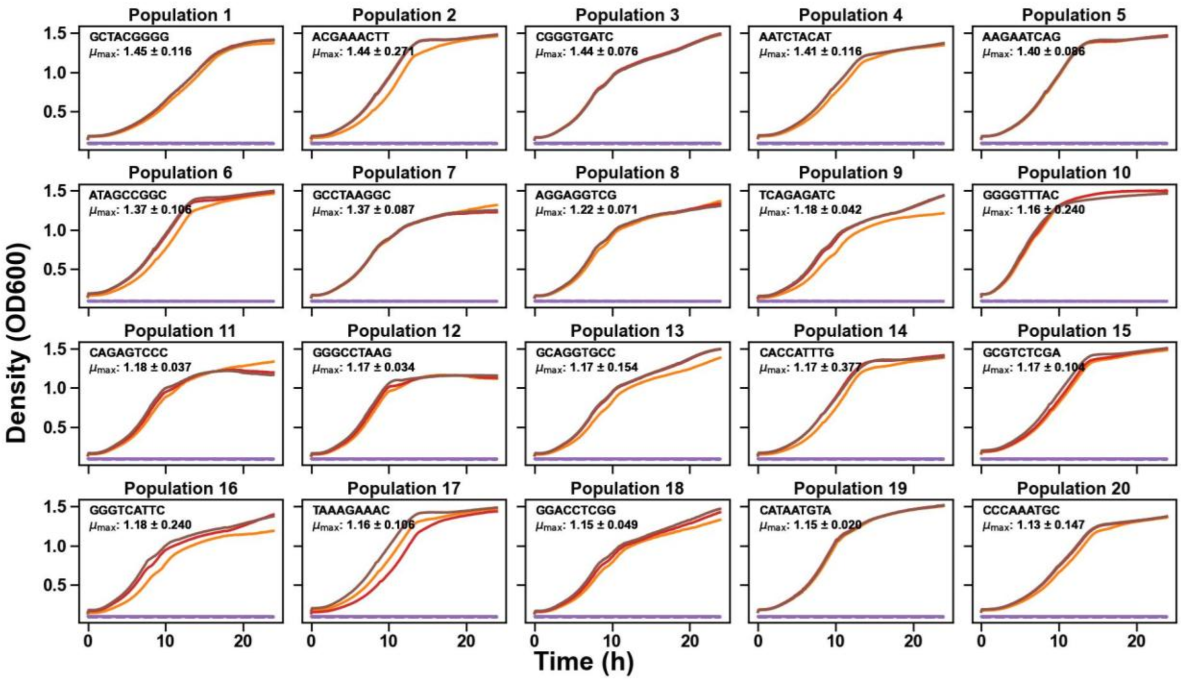

**Figure S10. Growth curves of 20 *E. coli* populations harboring 20 different *lac* operator variants from $H_G$ category in individual cultures grown in M9 medium with 1 mM lactose**. Each panel shows the increase in optical density ($OD_{600}$, vertical axis) of one *lac* operator variant in M9 medium containing 1 mM lactose (and 0.2% casamino acid) over a 24-hour period (horizontal axis). Each panel displays the growth curves for one of the 20 isolated clones (the *lac* operator sequence of the respective clone is shown within each panel) belonging to $H_G$ category (high fitness in glycerol and low fitness in lactose), with different colors representing $OD_{600}$ measurements of three replicate cultures for each variant. The dashed purple line in each panel represents a blank media control without cells. We used these growth curves to estimate the maximum growth rate, which is indicated in the upper left of each panel as the mean±one standard error (Methods).

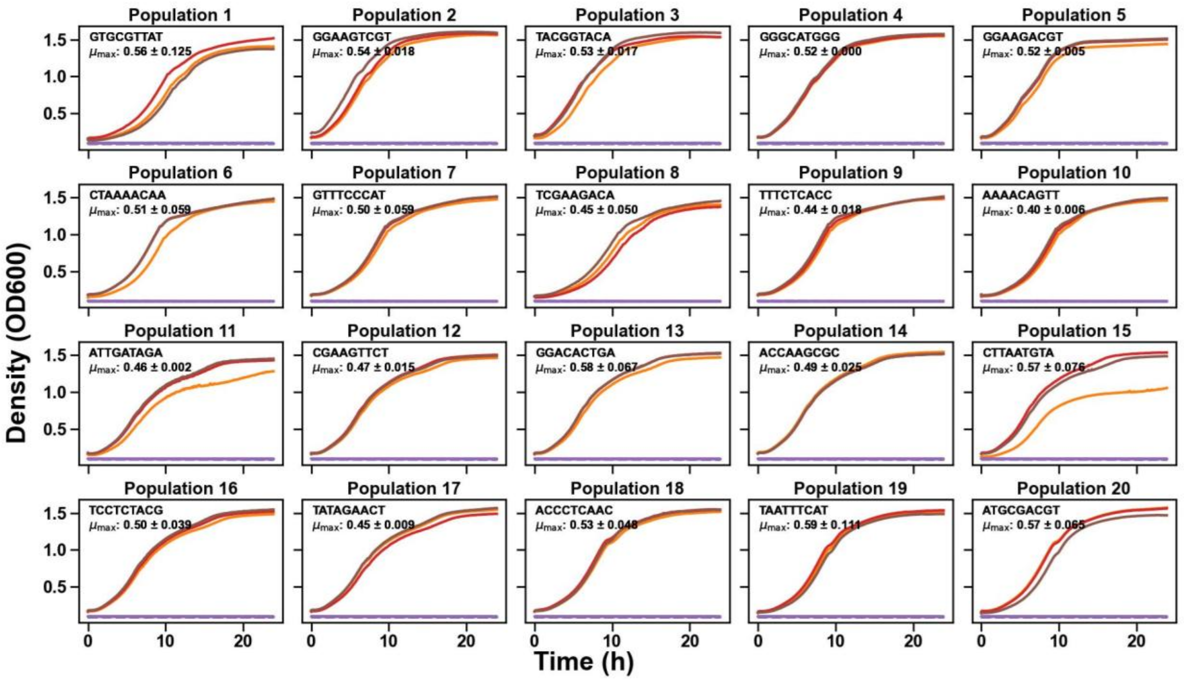

**Figure S11. Growth curves of 20 *E. coli* populations harboring 20 different *lac* operator variants from $L_{G+L}$ category in individual cultures grown in M9 medium with 10% glycerol**. Each panel shows the increase in optical density ($OD_{600}$, vertical axis) of one *lac* operator variant in M9 medium containing 10% glycerol (and 0.2% casamino acid) over a 24-hour period (horizontal axis). Each panel displays the growth curves for one of the 20 isolated clones (the *lac* operator sequence of the respective clone is shown within each panel) belonging to $L_{G+L}$ category (low fitness in both glycerol and lactose), with different colors representing $OD_{600}$ measurements of three replicate cultures for each variant. The dashed purple line in each panel represents a blank media control without cells. We used these growth curves were used to estimate the maximum growth rate, which is indicated in the upper left of each panel as the mean±one standard error (Methods).

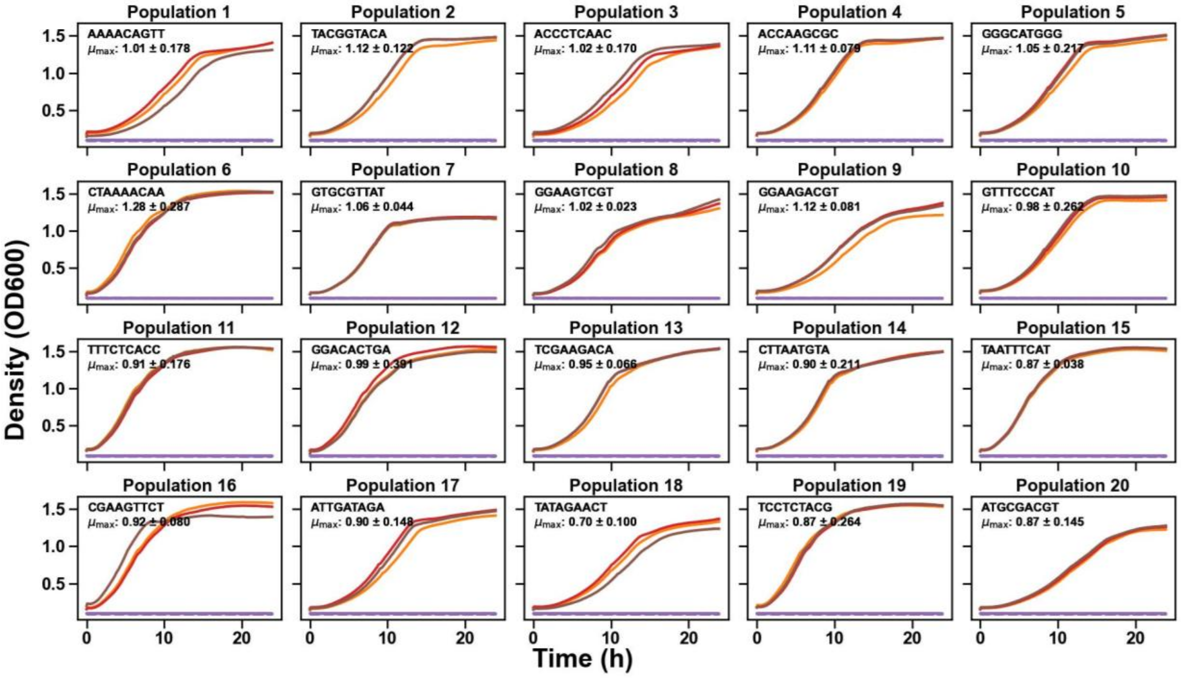

**Figure S12. Growth curves of 20 *E. coli* populations harboring 20 different *lac* operator variants from $L_{G+L}$ category in individual cultures grown in M9 medium with 1 mM lactose**. Each panel shows the increase in optical density ($OD_{600}$, vertical axis) of one *lac* operator variant in M9 medium containing 1 mM lactose (and 0.2% casamino acid) over a 24-hour period (horizontal axis). Each panel displays the growth curves for one of the 20 isolated clones (the *lac* operator sequence of the respective clone is shown within each panel) belonging to $L_{G+L}$ category (low fitness in both glycerol and lactose), with different colors representing $OD_{600}$ measurements of three replicate cultures for each variant. The dashed purple line in each panel represents a blank media control without cells. We used these growth curves were used to estimate the maximum growth rate, which is indicated in the upper left of each panel as the mean±one standard error (Methods).

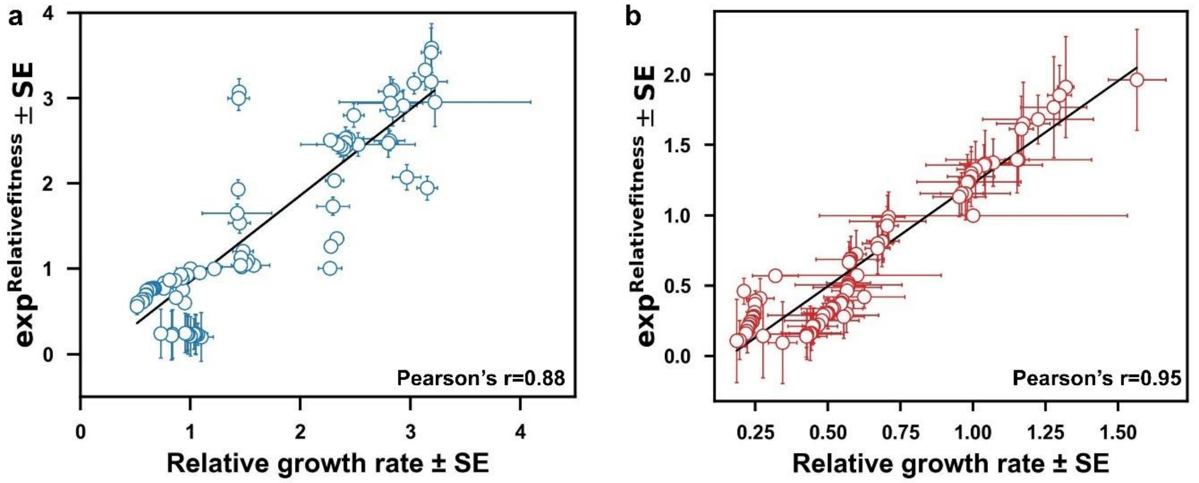

**Figure S13. Correlation between relative growth rates and relative fitness in glycerol and lactose.** We examined the relationship between fitness measured in the main selection experiment relative to the WT (vertical axis) and the maximum growth rate observed in single cultures of individual genotypes relative to the WT (horizontal axis). For these experiments We isolated 84 clones from the library and grew each clone in three replicate cultures to measure growth curves in either glycerol or lactose (see Fig. S13-20). The maximum growth rate during the exponential phase was calculated as the number of doublings per hour. Horizontal error bars represent the standard deviation of growth rates across the four replicate cultures used for fitness estimation by high-throughput sequencing. While vertical error bars indicate the standard errors of fitness estimates for individual genotypes (see Methods).

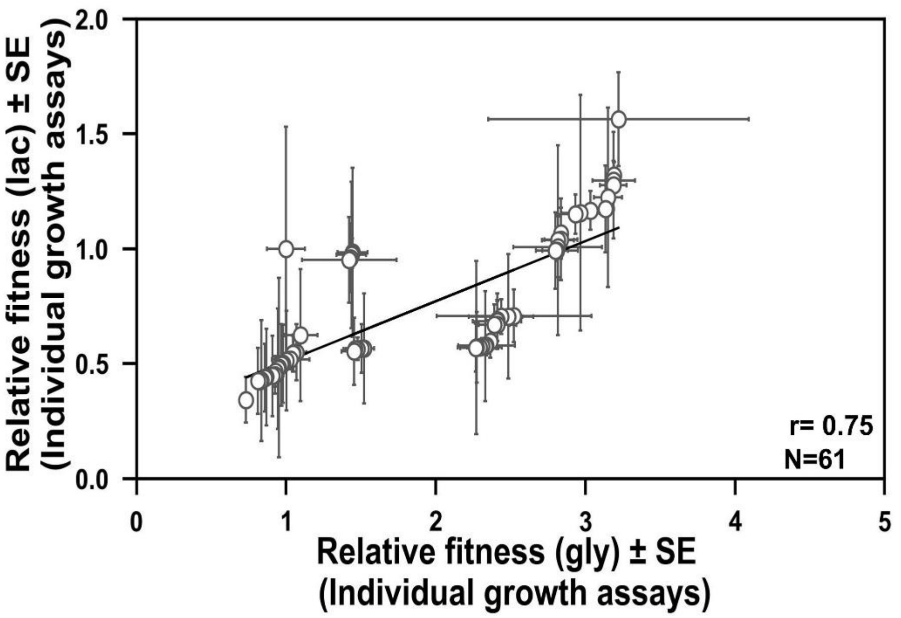

**Figure 14. Relative fitness (individual growth assays) measured in two environments.** Correlation between relative fitness estimated from individual growth assays of the same variants between lactose (vertical axis) and glycerol environments (horizontal axis). (β-gal enzyme activity: $R^2$=0.897, F-statistic = 124.09, F-test P= $1.11\times10^{-16}$, N=61; Individual growth assay: r= -0.75, P= P= $2.27\times10^{-12}$, N=61).

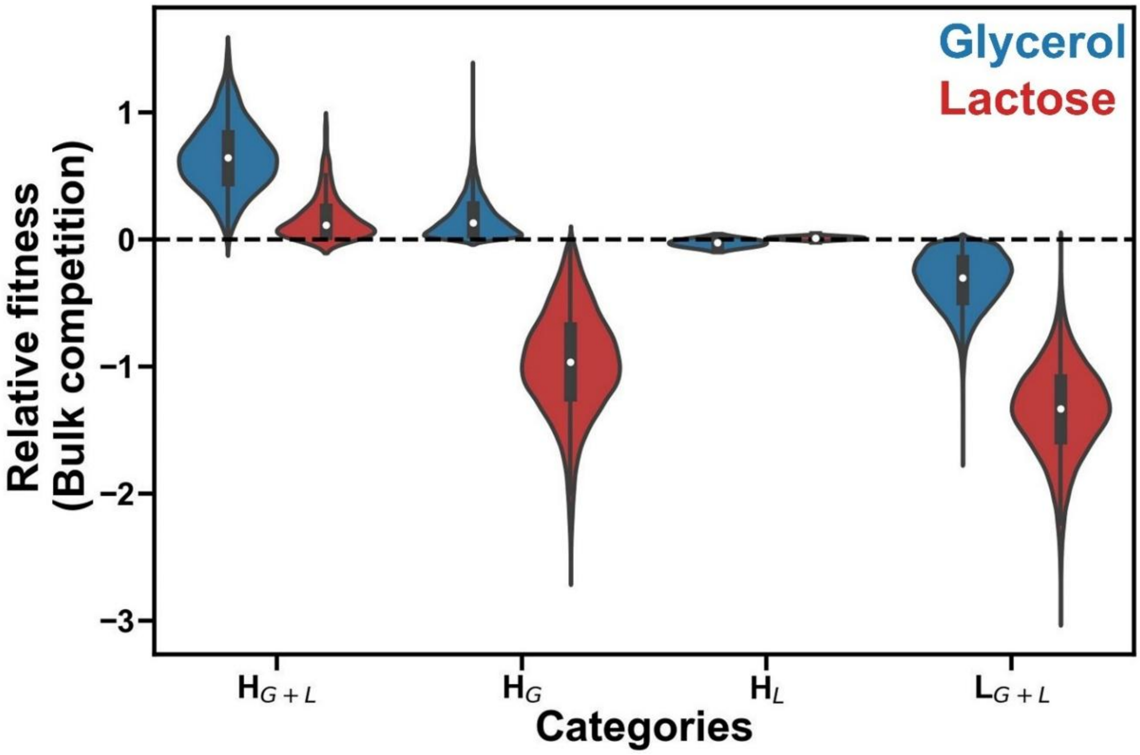

**Figure S15. Fitness distribution of *lac* operator variants across categories.** Violin plots show the fitness distribution of *lac* operator variants in glycerol (blue) and lactose (red) across four distinct fitness categories. Fitness is expressed relative to the WT (fitness zero, dashed black line). First category $H_{G+L}$ includes variants with higher fitness than WT in both glycerol and lactose (mean ± SD: 0.651 ± 0.257 in glycerol, 0.172 ± 0.168 in lactose). The second category $H_G$ comprises variants with high fitness in glycerol but low fitness in lactose (mean ± SD: 0.174 ± 0.152 in glycerol, -0.970 ± 0.405 in lactose). Third category $H_L$ consists of variants with low fitness in glycerol but high fitness in lactose (mean ± SD: -0.026 ± 0.029 in glycerol, 0.013 ± 0.012 in lactose). Fourth category $L_{G+L}$ comprises variants with low fitness in both environments (mean ± SD: -0.331 ± 0.211 in glycerol, -1.347 ± 0.356 in lactose). Mann-Whitney U tests revealed significant fitness differences between categories in both glycerol and lactose. In glycerol, genotypes in $H_{G+L}$ category had higher fitness than those in $H_G$ category ($U = 8.08 \times 10^6$, $P = 2.57 \times 10^{-167}$) and in $L_{G+L}$ category ($U = 3.43 \times 10^7$, $P = 8.41 \times 10^{-211}$), while genotypes in $H_G$ category had higher fitness than those in $L_{G+L}$ category ($U = 2.84 \times 10^9$, $P < 2\times10^{-308}$). In lactose, genotypes in $H_{G+L}$ category had higher fitness than genotypes in $H_G$ category ($U = 8.53 \times 10^6$, $P = 6.36 \times 10^{-209}$) and $L_{G+L}$ category ($U = 3.43 \times 10^7$, $P = 8.41 \times 10^{-211}$), and genotypes in $H_G$ category had higher fitness than genotypes in $L_{G+L}$ category ($U = 2.17 \times 10^9$, $P < 2\times10^{-308}$).

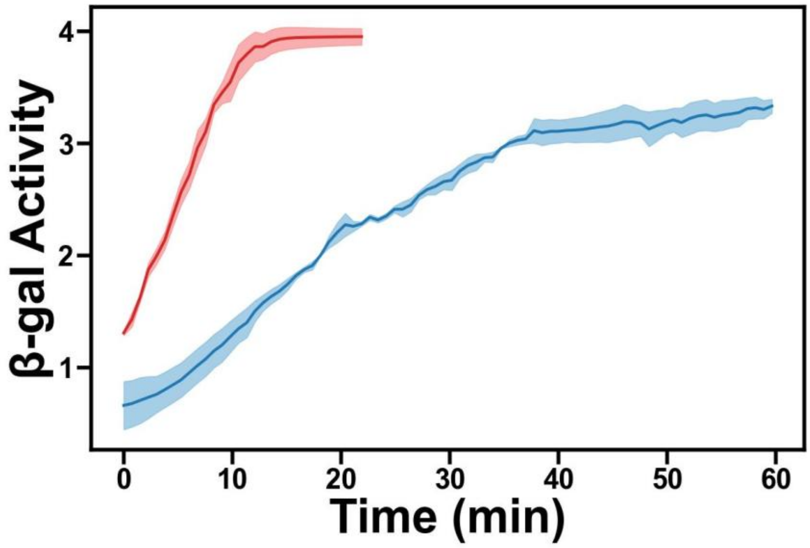

**Figure S16. β-Galactosidase activity of cells with the WT *lac* operator**. The panel shows the increase in optical density ($OD_{420}$, vertical axis) of these cells in M9 medium containing 10% glycerol (blue) and in M9 medium containing 1 mM lactose (red) over a 1-hour period (horizontal axis). The panel displays the mean β-galactosidase activity measured from three replicate cultures. We used these activity curves to calculate the slope ($OD_{420}$/min), which we then used to determine the β-galactosidase activity in Miller units (Methods).

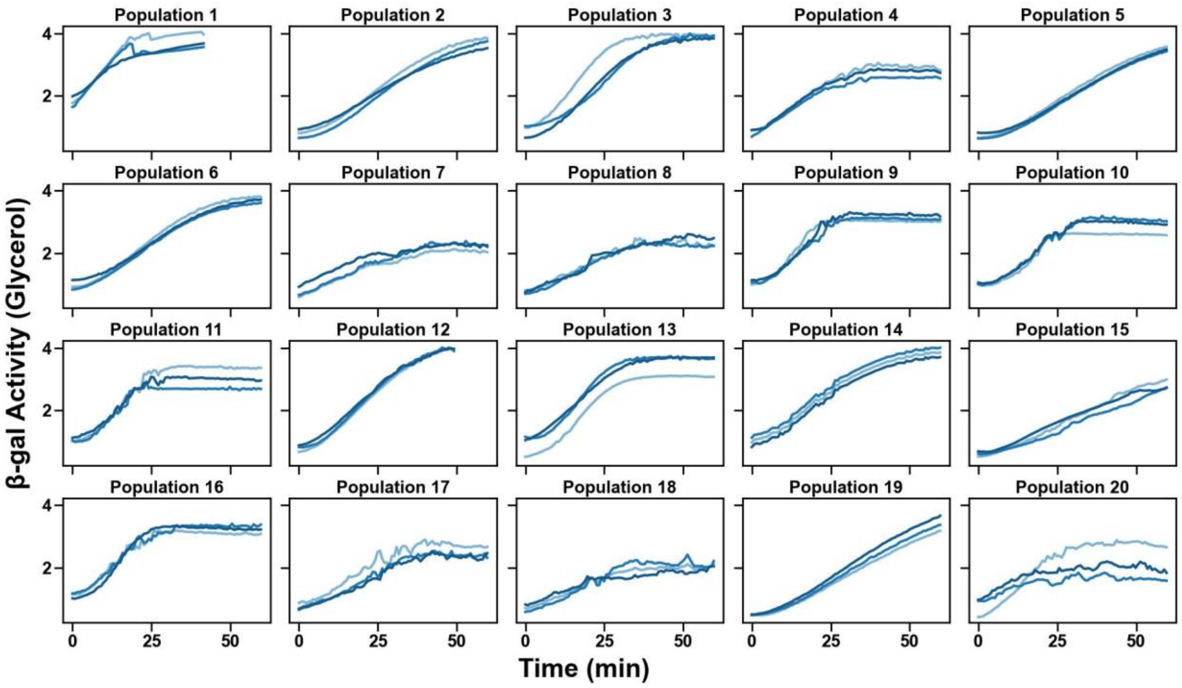

**Figure S17. β-Galactosidase activity of 20 *E. coli* populations harboring 20 different *lac* operator variants from $H_{G+L}$ category in individual cultures grown in M9 medium with 10% glycerol**. Each panel shows the increase in optical density ($OD_{420}$, vertical axis) of one *lac* operator variant in M9 medium containing 10% glycerol (and 0.2% casamino acid) over a 1-hour period (horizontal axis). Different colors in each panel represent $OD_{420}$ measurements of three replicate cultures for each variant. Data in the panels comes from $H_{G+L}$ category variants (high fitness in both glycerol and lactose). We used the activity curves to calculate the slope ($OD_{420}$/min), which allowed us to determine β-galactosidase activity in Miller units (Methods).

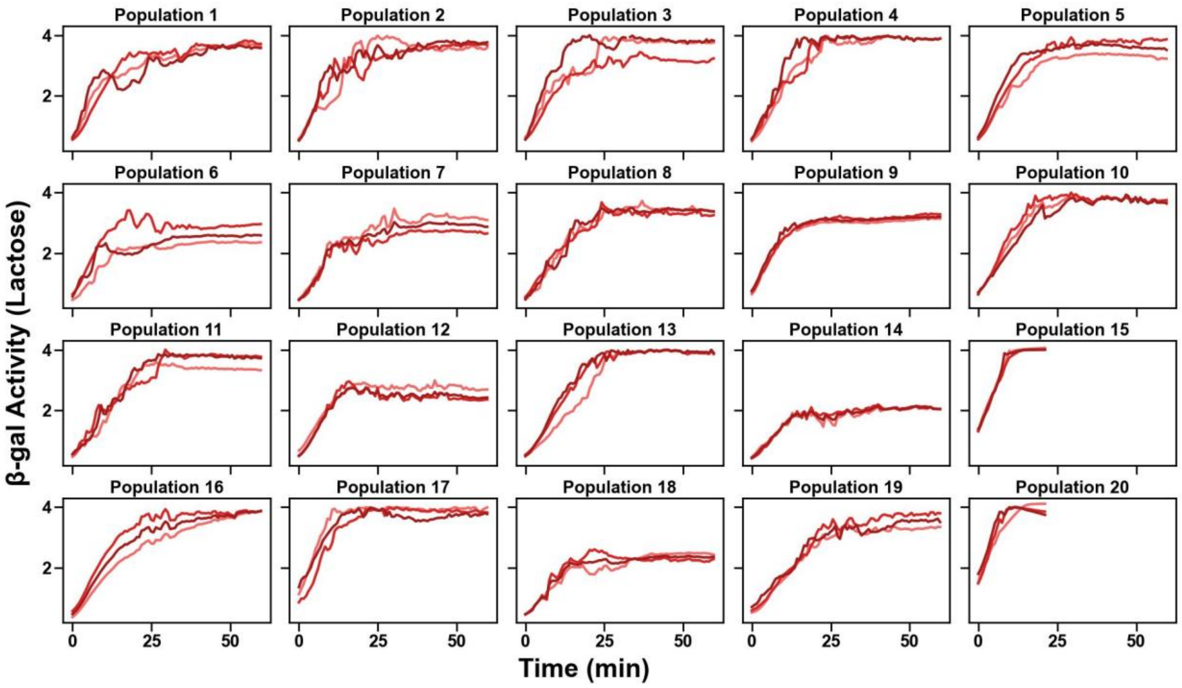

**Figure S18. β-Galactosidase activity of 20 *E. coli* populations harboring 20 different *lac* operator variants from $H_{G+L}$ category in individual cultures grown in M9 medium with 1 mM lactose**. Each panel shows the increase in optical density ($OD_{420}$, vertical axis) of one *lac* operator variant in M9 medium containing 1 mM lactose (and 0.2% casamino acid) over a 1-hour period (horizontal axis). Differently colored trajectories in each panel represent $OD_{420}$ measurements of three replicate cultures for each variant. Data in the panels comes from $H_{G+L}$ variants (high fitness in both glycerol and lactose). We used the activity curves to calculate the slope ($OD_{420}$/min), which allowed us to determine β-galactosidase activity in Miller units (Methods).

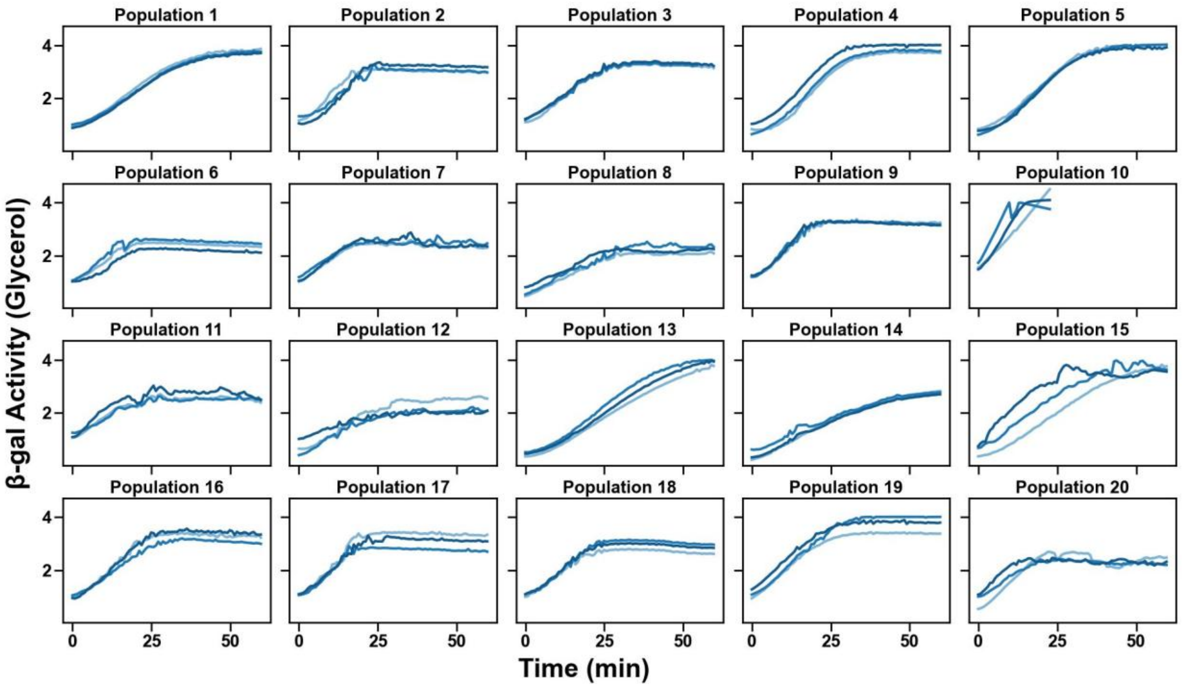

**Figure S19. β-Galactosidase activity of 20 *E. coli* populations harboring 20 different *lac* operator variants from $H_G$ category in individual cultures grown in M9 medium with 10% glycerol**. Each panel shows the increase in optical density ($OD_{420}$, vertical axis) of one *lac* operator variant in M9 medium containing 10% glycerol (and 0.2% casamino acid) over a 1-hour period (horizontal axis). Differently colored trajectories in each panel represent $OD_{420}$ measurements of three replicate cultures for each variant. Data in the panels comes from $H_G$ variants (high fitness in glycerol and low fitness in lactose). We used the activity curves to calculate the slope ($OD_{420}$/min), which allowed us to determine β-galactosidase activity in Miller units (Methods).

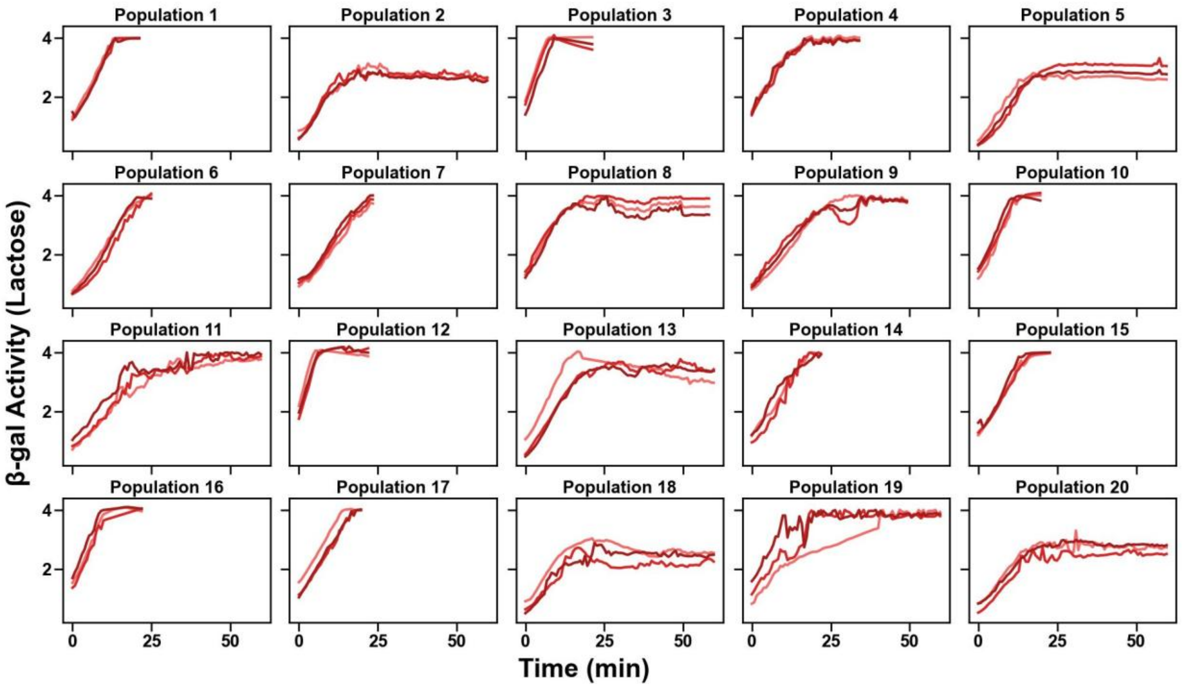

**Figure S20. β-Galactosidase activity of 20 *E. coli* populations harboring 20 different *lac* operator variants from $H_G$ category in individual cultures grown in M9 medium with 1 mM lactose**. Each panel shows the increase in optical density ($OD_{420}$, vertical axis) of one *lac* operator variant in M9 medium containing 1 mM lactose (and 0.2% casamino acid) over a 1-hour period (horizontal axis). Differently colored trajectories in each panel represent $OD_{420}$ measurements of three replicate cultures for each variant. Data in the panels comes from $H_G$ variants (high fitness in glycerol and low fitness in lactose). We used the activity curves to calculate the slope ($OD_{420}$/min), which allowed us to determine β-galactosidase activity in Miller units (Methods).

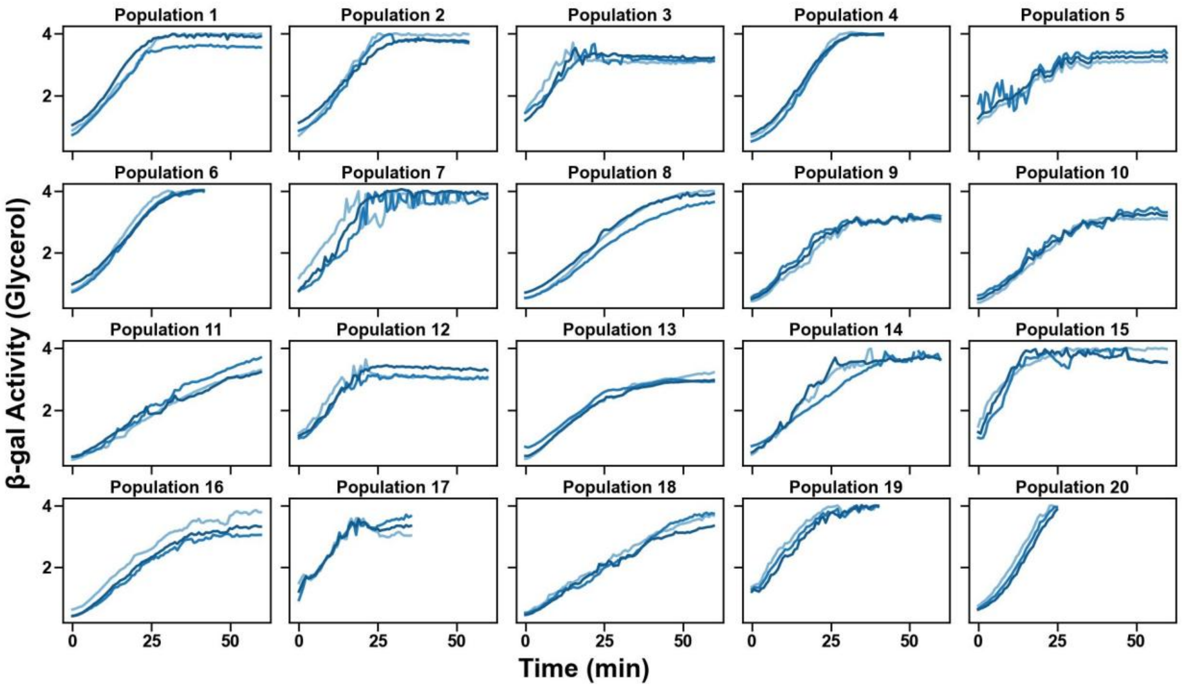

**Figure S21. β-Galactosidase activity of 20 *E. coli* populations harboring 20 different *lac* operator variants from $L_{G+L}$ category in individual cultures grown in M9 medium with 10% glycerol**. Each panel shows the increase in optical density ($OD_{420}$, vertical axis) of one *lac* operator variant in M9 medium containing 10% glycerol (and 0.2% casamino acid) over a 1-hour period (horizontal axis). Differently colored trajectories in each panel represent $OD_{420}$ measurements of three replicate cultures for each variant. Data in the panels comes from $L_{G+L}$ variants (low fitness in both glycerol and lactose). We used the activity curves to calculate the slope ($OD_{420}$/min), which allowed us to determine β-galactosidase activity in Miller units (Methods).

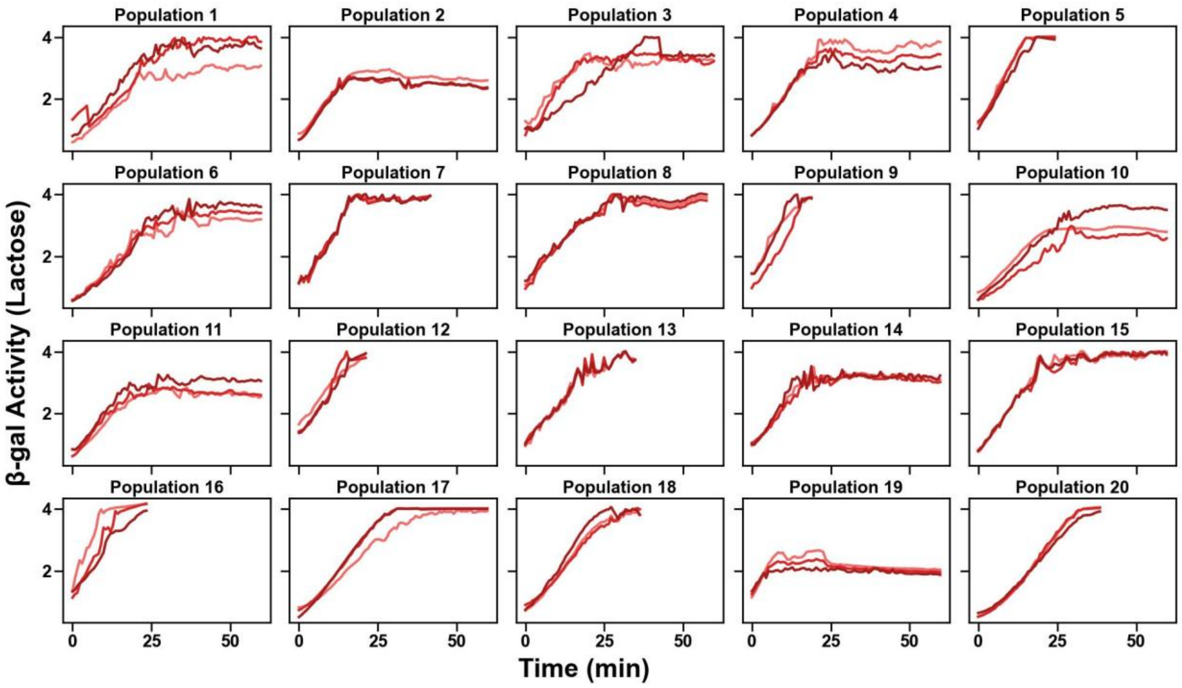

**Figure S22. β-Galactosidase activity of 20 *E. coli* populations harboring 20 different *lac* operator variants from $L_{G+L}$ category in individual cultures grown in M9 medium with 1 mM lactose**. Each panel shows the increase in optical density ($OD_{420}$, vertical axis) of one *lac* operator variant in M9 medium containing 1 mM lactose (and 0.2% casamino acid) over a 1-hour period (horizontal axis). Differently colored trajectories in each panel represent $OD_{420}$ measurements of three replicate cultures for each variant. Data in the panels comes from $L_{G+L}$ variants (low fitness in both glycerol and lactose). We used the activity curves to calculate the slope ($OD_{420}$/min), which allowed us to determine β-galactosidase activity in Miller units (Methods).

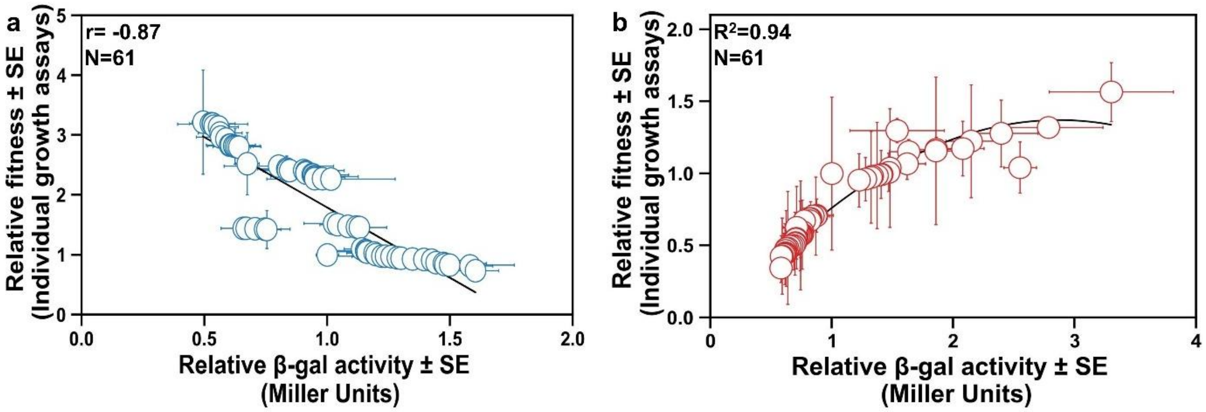

**Figure S23. Relationship between relative enzyme activity and fitness measured from individual growth assays in different media.** (a) Relative β-gal enzyme (horizontal axis) and its relationship to relative fitness estimated from individual growth assay (vertical axis) in glycerol media. (b) like (a), but in lactose media (Glycerol: r= -0.87, P= $4.37\times10^{-20}$, N=61; Lactose: $R^2$=0.935, F-statistic = 67.00, F-test P= $2.99\times10^{-11}$, N=61).

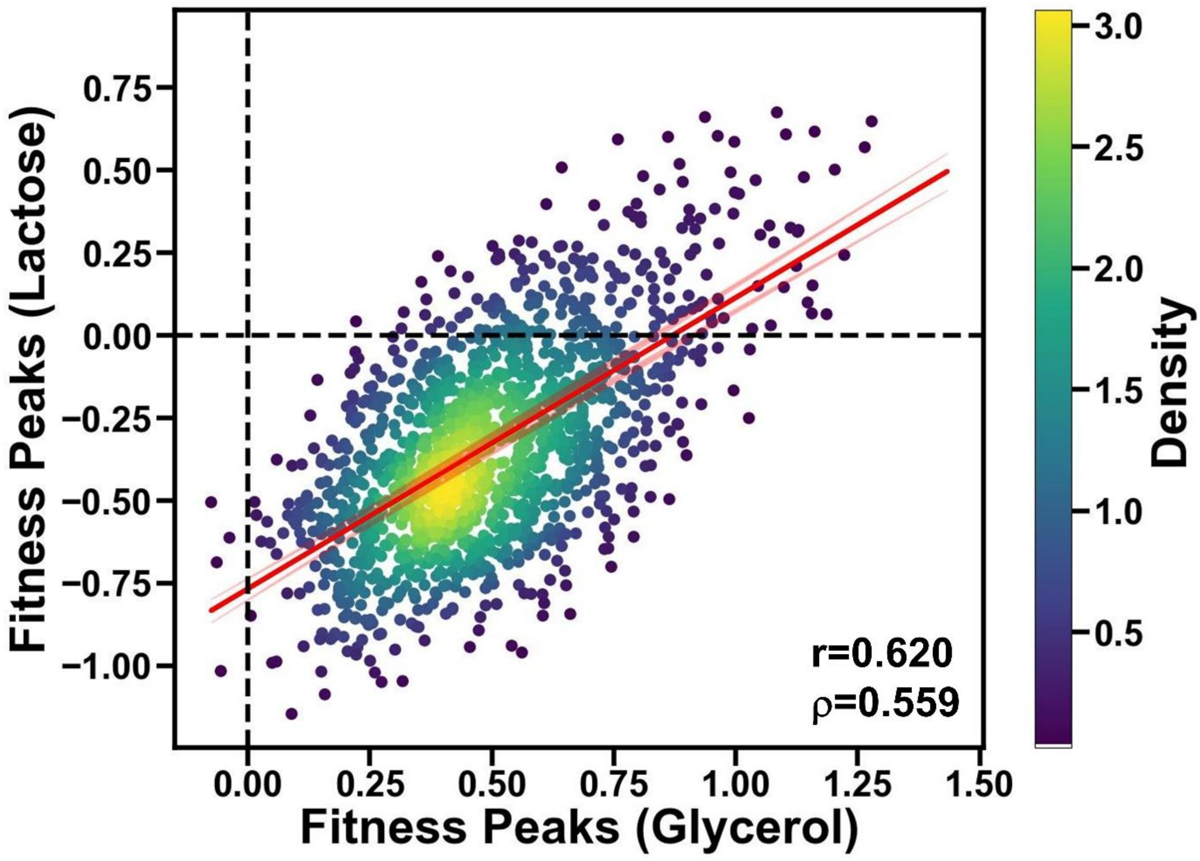

**Figure S24. Fitness distribution of peak genotypes in lactose and glycerol environments.** The density of points is represented by a color gradient, with lighter areas indicating higher density. Each circle represents fitness data from a genotype that is a peak in both environments. A Gaussian GLM model is fitted to the data, shown by the red regression line, with 95% confidence intervals shaded in red. Dashed black lines represent zero (wild-type) fitness values in each environment. Pearson's r = 0.620, $P=1.022\times10^{-151}$, Spearman's $\rho$ = 0.559, $P=1.577\times10^{-117}$, n=1423.

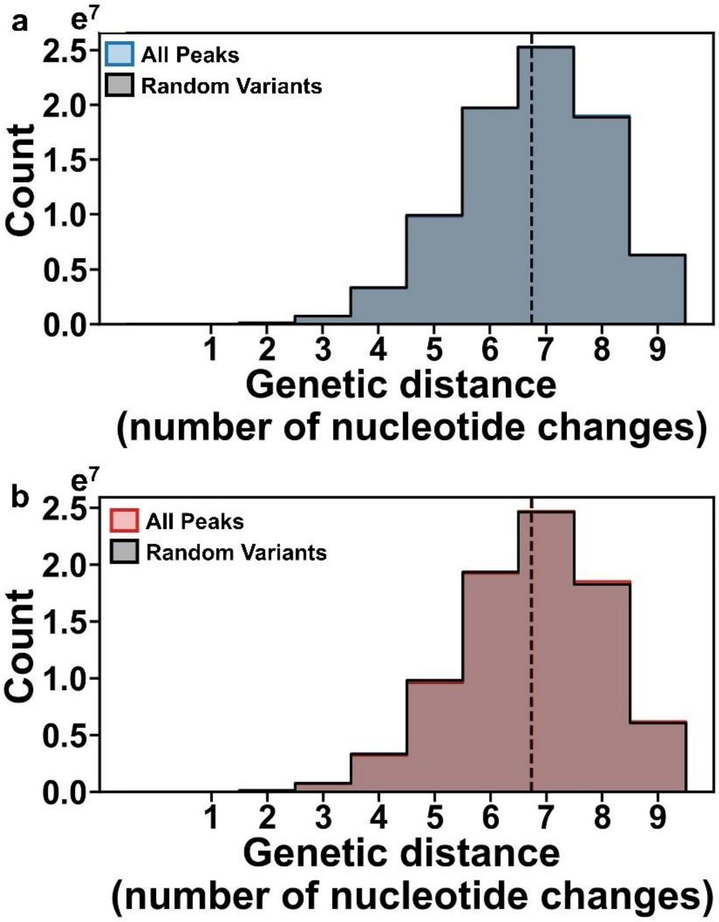

**Figure S25. Genetic distance distribution of all peaks in *lac* operator landscape across different environmental conditions.** (a) Distribution of pairwise genetic distances between nucleotide sequences of 9183 peaks (blue) in glycerol and 9183 randomly chosen genotypes (black), with vertical lines indicating the mean values (d=6.75 for peaks [in blue] and d=6.74 for random variants [in black]). (b) Distribution of pairwise genetic distances between nucleotide sequences of 9074 peaks (red) in lactose and between sequences of 9074 randomly chosen variants (black), with vertical lines indicating the mean values (d=6.75 for peaks [in red] and d=6.73 for random variants [in black]).

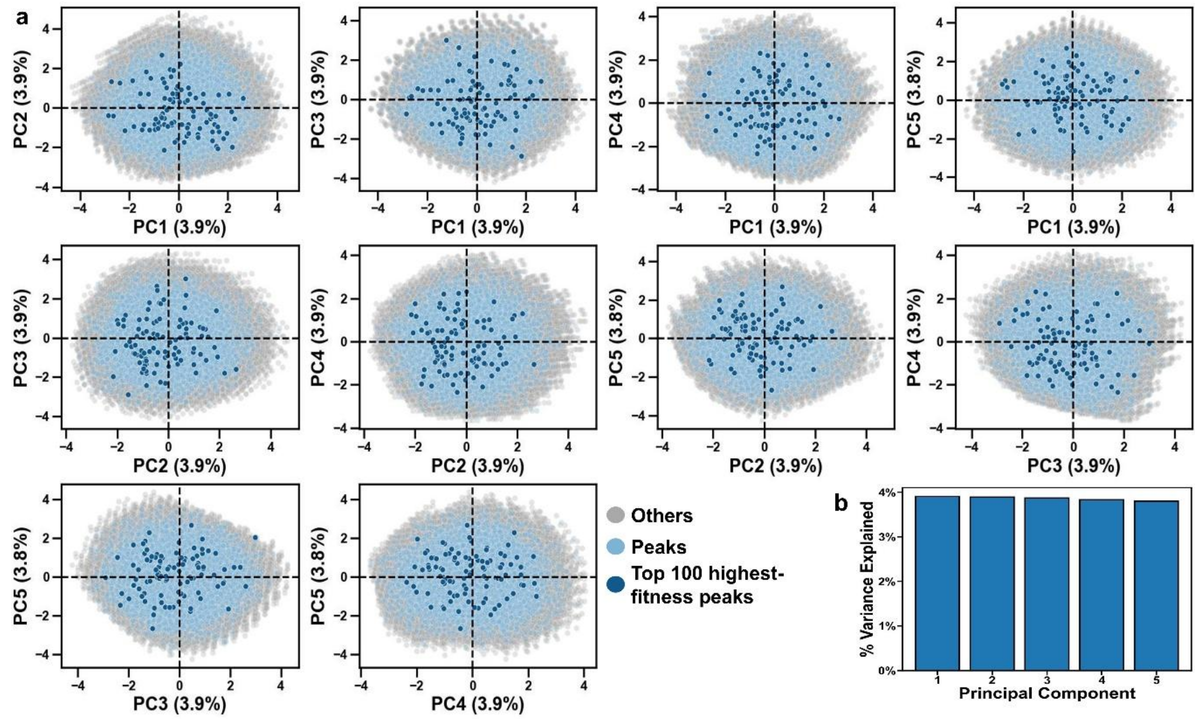

**Figure S26. Principal component analysis of fitness peaks in the glycerol landscape's genotype space.** (a) We performed PCA using a one-hot encoded representation of *lac* operator genotypes for all variants in the landscape. The panels present the projections of various pairs of the first five principal components (PCs). Each circle represents one of the 187,484 genotypes. Dark blue indicates one of the top 100 fitness peaks, light blue indicates any other fitness peak, and grey indicates non-peak variants. (b) The proportion of variance explained by each principal component. Due to the combinatorial complexity and high-dimensional nature of the genotype space, each principal component captures only a small fraction of the overall genetic variation.

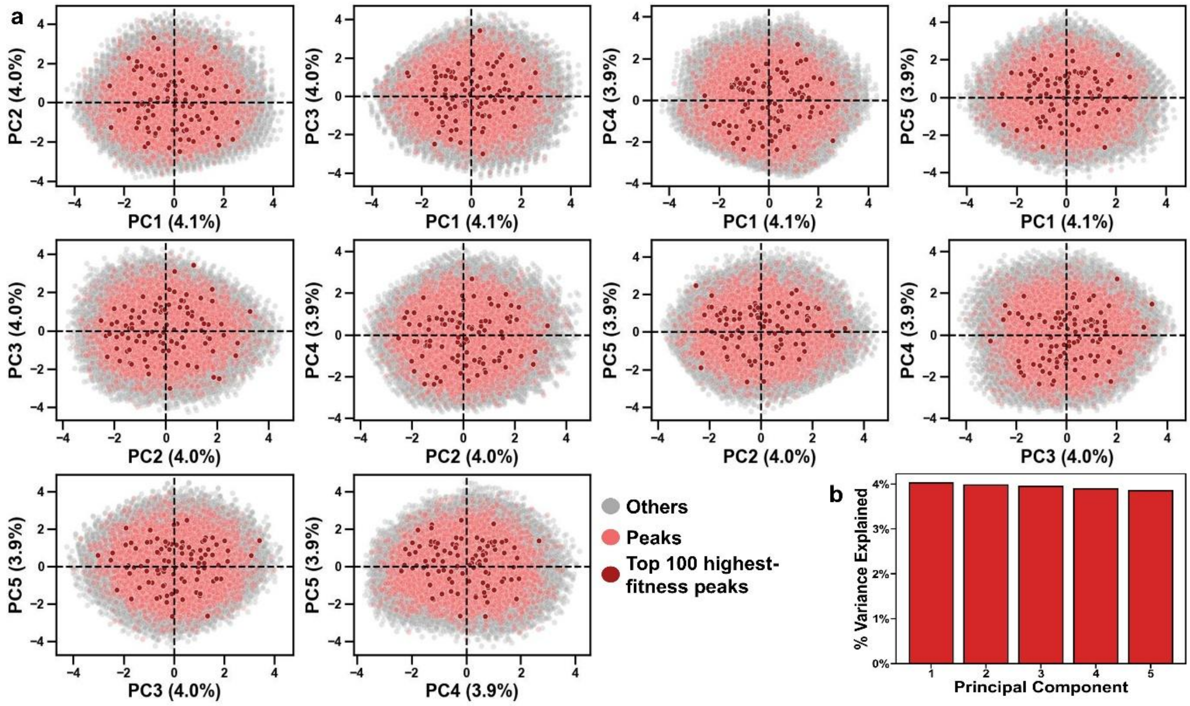

**Figure S27. Principal component analysis of fitness peaks in the lactose landscape's genotype space.** (a) We performed PCA using a one-hot encoded representation of *lac* operator genotypes for all variants in the landscape. The panels present the projections of various pairs of the first five principal components (PCs). Each circle represents one of the 141,147 genotypes. Dark red indicates one of the top 100 fitness peaks, light red indicates any other fitness peak, and grey indicates non-peak variants. (b) The proportion of variance explained by each principal component. Due to the combinatorial complexity and high-dimensional nature of the genotype space, each principal component captures only a small fraction of the overall genetic variation.

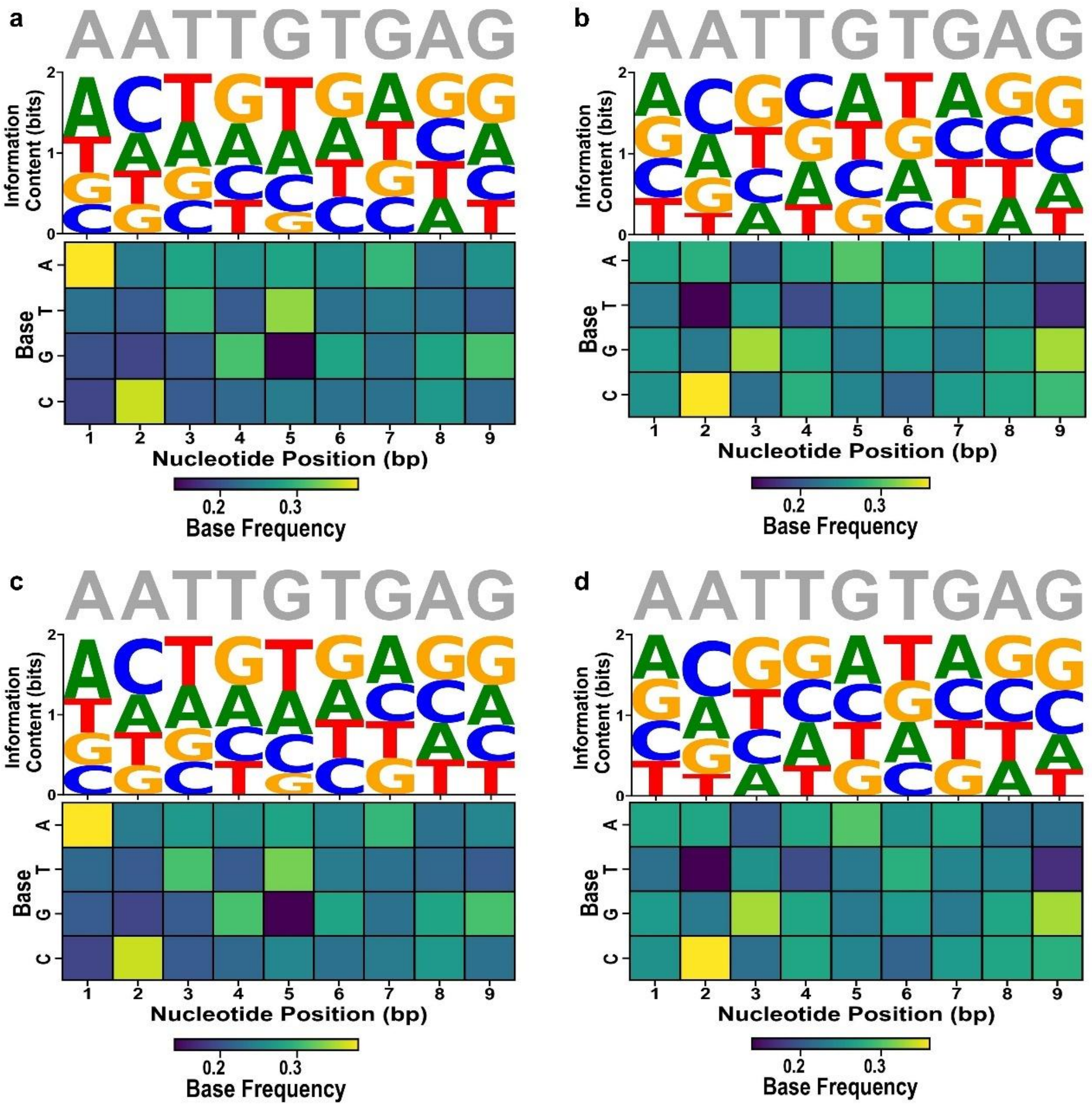

**Figure S28. Sequence logo representing the *lac* operator site.** Grey letters on top indicate the WT (*lac* operator) nucleotides. (a-b) We aligned the top 100 sequences to construct a DNA sequence logo for the *lac* operator site obtained from (a) the glycerol and (b) the lactose environments. (c-d) Same as (a-b), but from the sequences of the top 100 fitness peaks in the (c) glycerol and (d) lactose environments. We generated the sequence logo by scaling nucleotide frequencies according to their contribution to Shannon entropy (Methods), providing a visual representation of conserved and variable positions. Below each logo, a heatmap displays the frequency matrix for each variable nucleotide position, with the vertical axis representing nucleotides (A, T, G, or C) and the horizontal axis representing each position in the sequence. The heatmap's color gradient reflects the frequency of each nucleotide at each position, with the color legend indicating the frequency distribution.

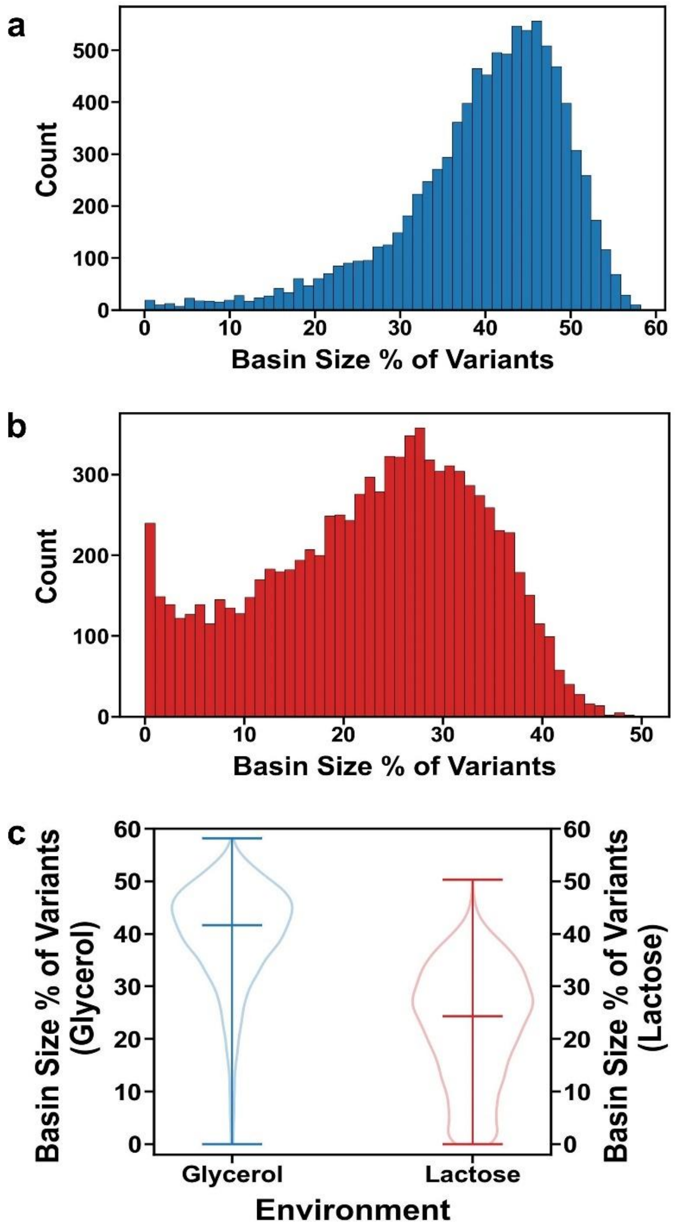

**Figure S29. Fitness peaks in glycerol have larger basins of attraction than in lactose.** (a-b) Basin sizes of all fitness peaks in (a) glycerol and (b) lactose. Basin size is indicated as the percentage of the total number of the variants in the landscape. (c) Distribution of basin sizes as a percentage of variants in both glycerol and lactose environments. The left vertical axis represents the basin size as a percentage of variants in glycerol (red), while the right vertical axis represents the basin size as a percentage of variants in lactose (green). The middle horizontal line within each violin plots marks the median, and the upper and lower horizontal lines mark the range of the data. Average basin size of peaks is 39.80%±9.74% (mean±SD; 70,969/178,301 variants) in glycerol and 22.77%±10.96% (mean±SD; 30,071/132,073 variants) in lactose (Mann-Whitney U statistic: 9,621,941.000, $P<2.0\times10^{-308}$), where average and standard deviations are calculated for the distribution of basin sizes across all peaks.

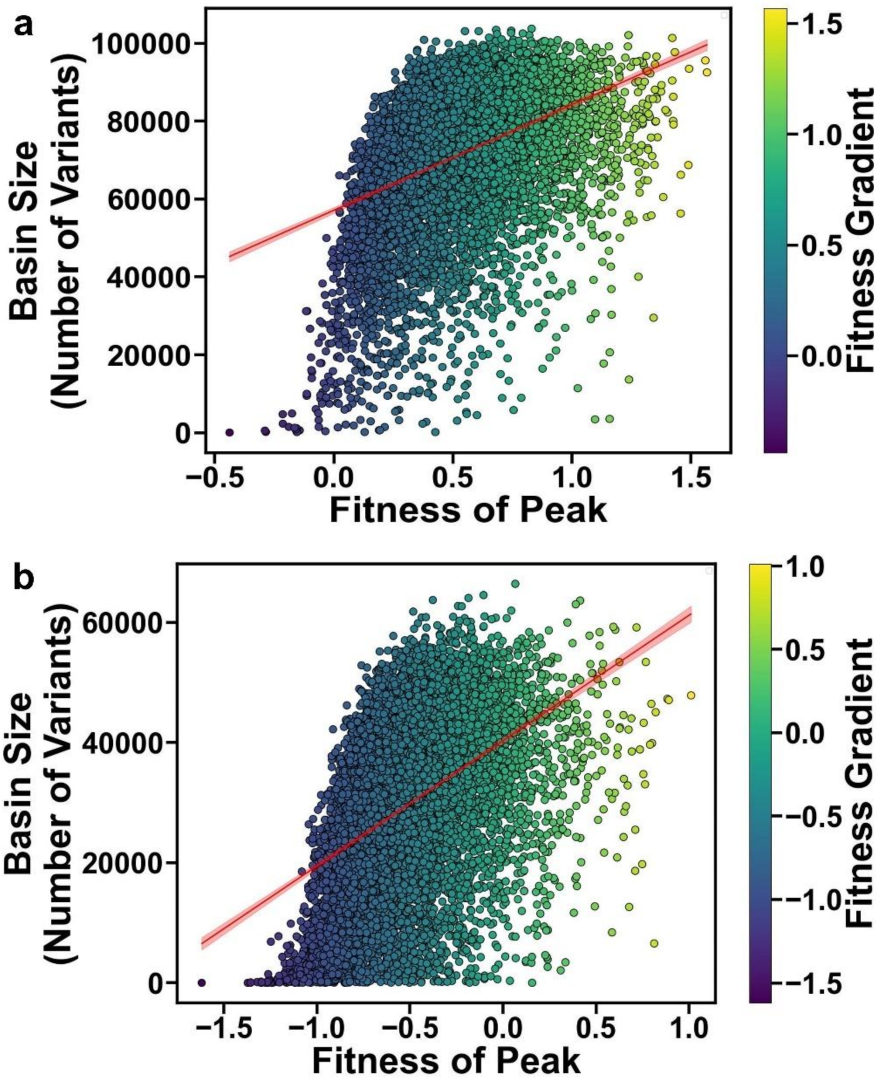

**Figure S30. Basin sizes correlate with peak fitness.** (a-b) The plots show the relationship between peak fitness (x-axis) and basin size (y-axis in absolute numbers) in (a) glycerol and (b) lactose. Each circle corresponds to one peak and is color-coded by fitness (see legend). The red line is a linear regression line and the shaded area its 95% confidence interval.

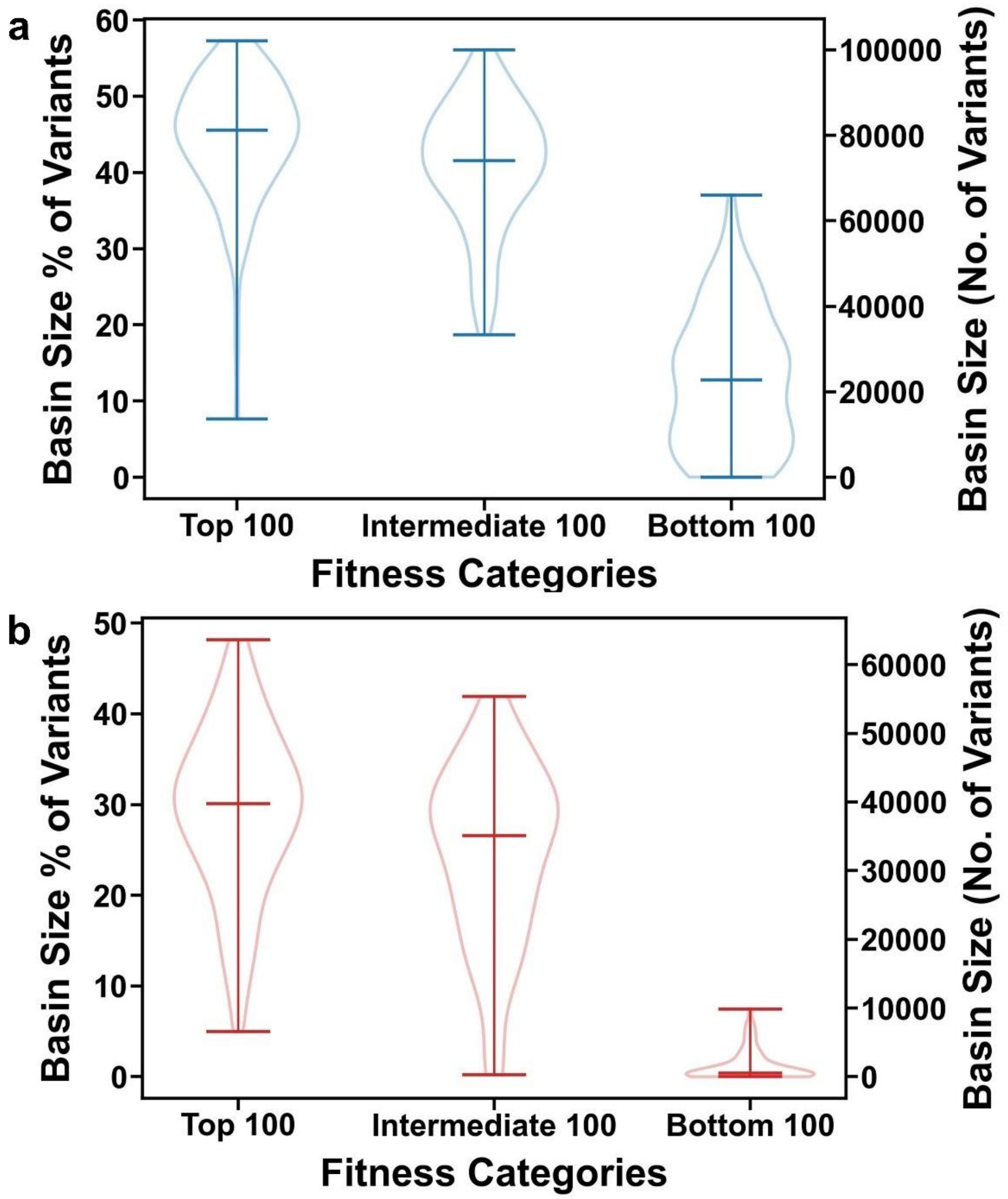

**Figure S31. The top 100 fitness peaks have larger basins of attraction sizes than 100 peaks at intermediate fitness and the bottom 100 peaks.** The vertical axes indicate the basin sizes of peak genotypes as a percentage of all genotypes (left vertical axes) and as the number of genotypes (right vertical axis) in (a) glycerol and (b) lactose environments. The horizontal axis differentiates between the top 100 high fitness peaks, 100 peaks at intermediate fitness, and the bottom 100 peaks. We defined intermediate peaks as the 100 peaks in the middle of the fitness distribution. The middle horizontal line within each violin plots marks the median, and the upper and lower horizontal lines mark the range of the data.

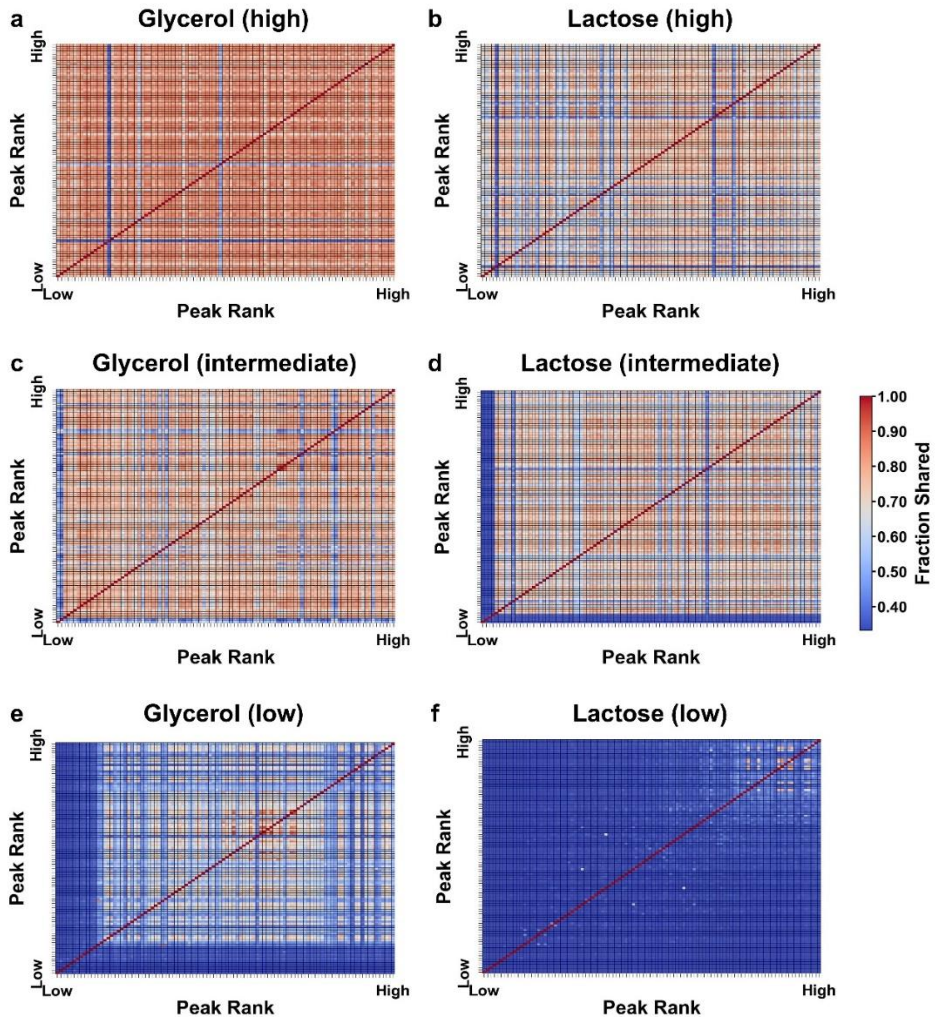

**Figure S32. The basins of attraction associated with the top, intermediate and bottom 100 peaks show high, moderate to low overlap.** The heatmap display the fraction of *lac* operator variants shared (color legend) between the basins of attraction of landscape peaks in glycerol (a, c, e) and lactose (b, d, f) environments, for all pairs of the top 100 fitness peaks (a, b), 100 intermediate fitness peaks (c, d), and for all pairs of the 100 lowest fitness peaks (e, f) (Methods). We defined intermediate peaks as the 100 peaks in the middle of the fitness distribution. Each row and column correspond to a specific peak, and each entry in the matrix represents the overlap (Jaccard index, Methods) between the basins of attraction respective pairs of peaks. A value of one (red) signifies that the basins of attraction for the compared peaks are identical, i.e., they share every single *lac* operator variant, while a value of zero (blue) indicates that they share no variants. In all panels the peaks are ranked by fitness, from high to low on the y-axis and from low to high on the x-axis. On average, the basins of 100 intermediate peaks in glycerol shared 71.7% ± 10.6% of their variants (61086.0 ± 12545.8 variants, mean±SD, N = 4950), while the bottom 100 peaks shared 28.0% ± 20.3% (10797.1 ± 9897.4 variants, mean±SD, N = 4950). In lactose, the intermediate 100 peaks overlapped on average by 49.7% ± 19.6% (22094.8 ± 10633.0 variants, mean±SD, N = 4950) of variants, whereas the bottom 100 peaks overlapped by only 2.5% ± 5.7% (142.1 ± 461.8 variants, mean±SD, N = 4950).

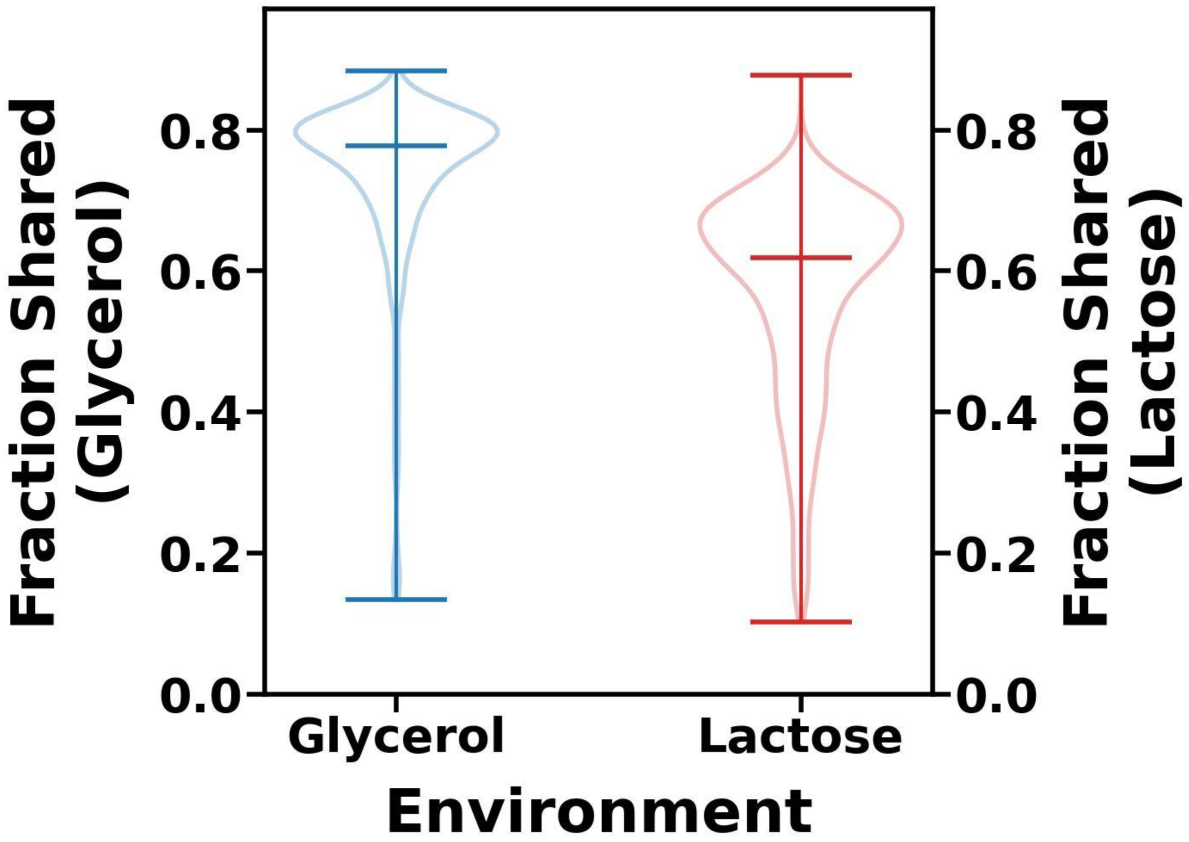

**Figure S33. The top 100 fitness peaks in glycerol have more shared variants than those in lactose.** The vertical axes indicate the fraction of shared variants of the top 100 fitness peaks in two environments, glycerol and lactose. For each environment, we calculated how many genotypes are shared between each pair of peaks. The left vertical axis represents the fraction of shared variants in glycerol (blue), while the right vertical axis represents the fraction of shared variants in lactose (red). The horizontal lines within the violin plots mark the median values, and the vertical extent of the plots represents the range of the data. Average basin size of the top 100 peaks is 73.8% ± 12.2% (67,347.2 ± 13,820.6 variants, N=4950) in glycerol and 57.1% ± 14.4% (28,429.1 ± 9,732.0 variants, N=4950) in lactose (two-sided Mann–Whitney U = 2,925,859, $P < 2.0\times10^{-308}$).

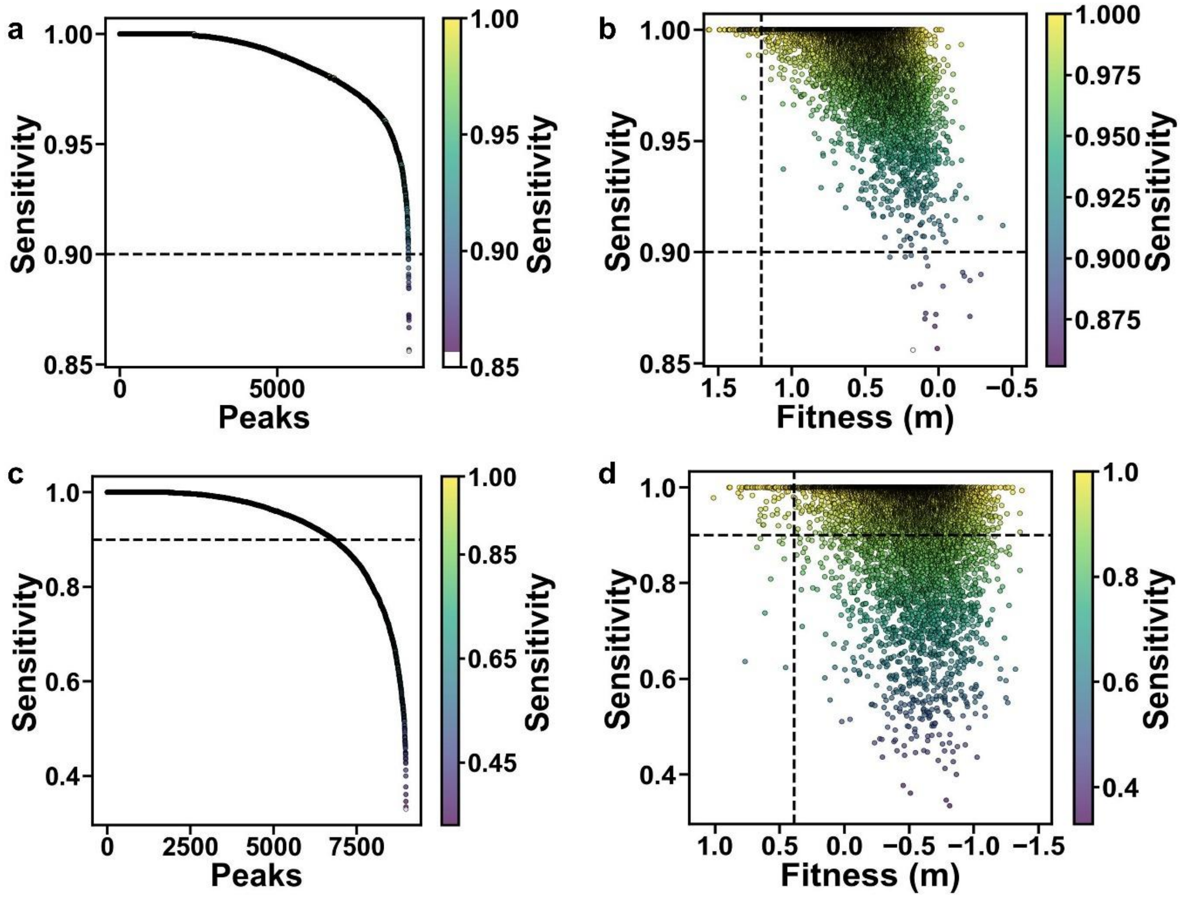

**Figure S34. Sensitivity of peak identification in glycerol and lactose.** Sensitivity was estimated for each putative peak by generating synthetic datasets based on binomial sampling of sequencing read counts (see Supplementary Methods 15). The sensitivity of the peak identification procedure, defined as the fraction of datasets where the peak was correctly identified, is shown for 50 iterations in (a) glycerol and (c) lactose, with peaks ordered by sensitivity. Scatter plots show the relationship between peak fitness and sensitivity of peak identification in (b) glycerol and (d) lactose. The dotted vertical lines at x = 1.207 and x = 0.389 mark the fitness of the lowest among the top 100 peaks in glycerol and lactose, respectively. In all panels circles represent data for individual peaks. Each peak is also colored according to the color legend (side bar). The dotted horizontal line at y = 0.9 in each panel indicates the threshold for high sensitivity. Most high-fitness peaks are identified with high sensitivity, confirming the reliability of peak calling in both environments.

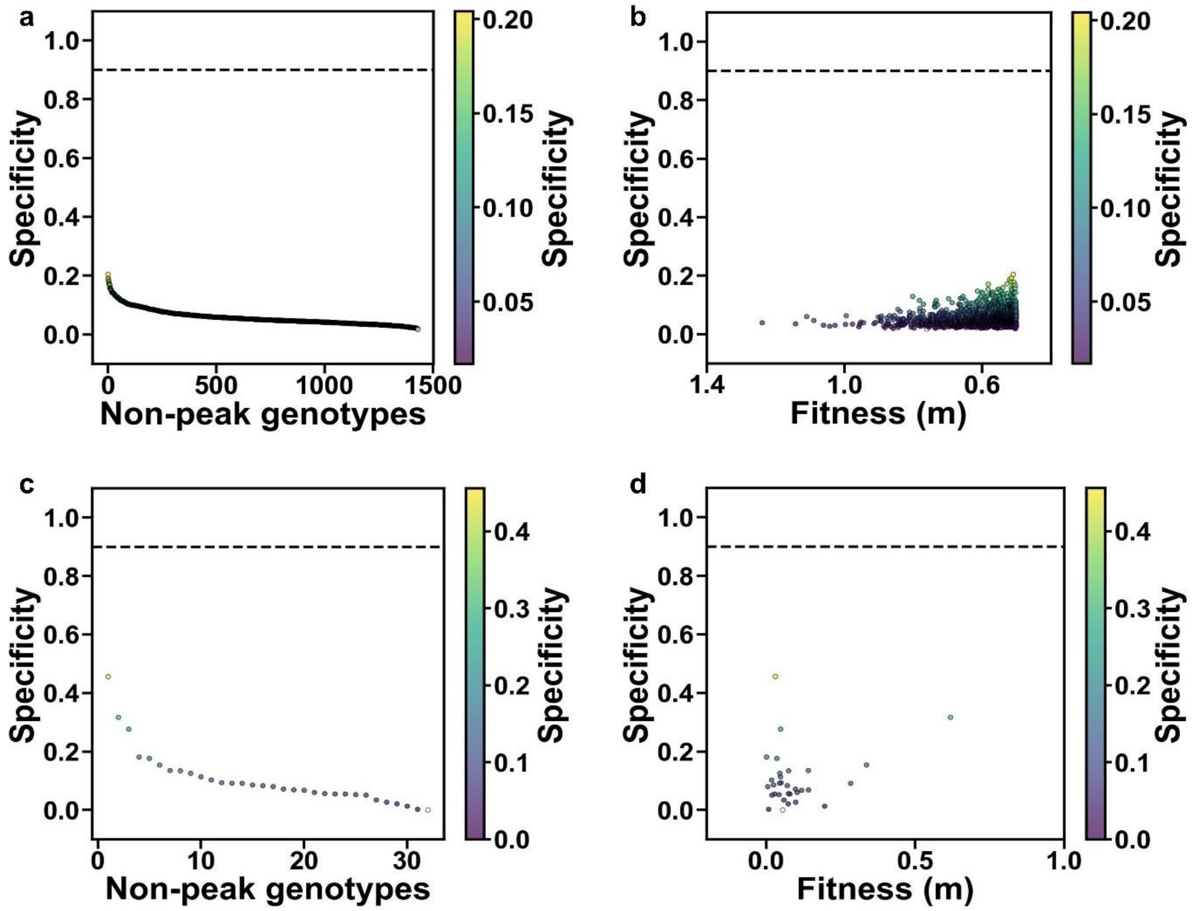

**Figure S35. Specificity of high fitness non-peaks variants in glycerol and lactose.** Specificity was estimated for each high-fitness non-peak genotype by generating synthetic datasets based on binomial sampling of sequencing read counts (see Supplementary Methods 15). The specificity of the peak identification procedure, defined as the fraction of datasets where the non-peak was correctly classified as not a peak, is shown for 50 iterations in (a) glycerol and (c) lactose, with variants ordered by specificity. Scatter plots show the relationship between variant fitness and specificity of peak identification in (b) glycerol and (d) lactose. In all panels circles represent data for individual non-peak genotypes. Each non-peak genotype is also colored according to the color legend (side bar). The dotted horizontal line at y = 0.9 in each panel represents the threshold for high specificity. High-fitness non-peaks are generally classified correctly, demonstrating the robustness of the peak-calling method against false positives.

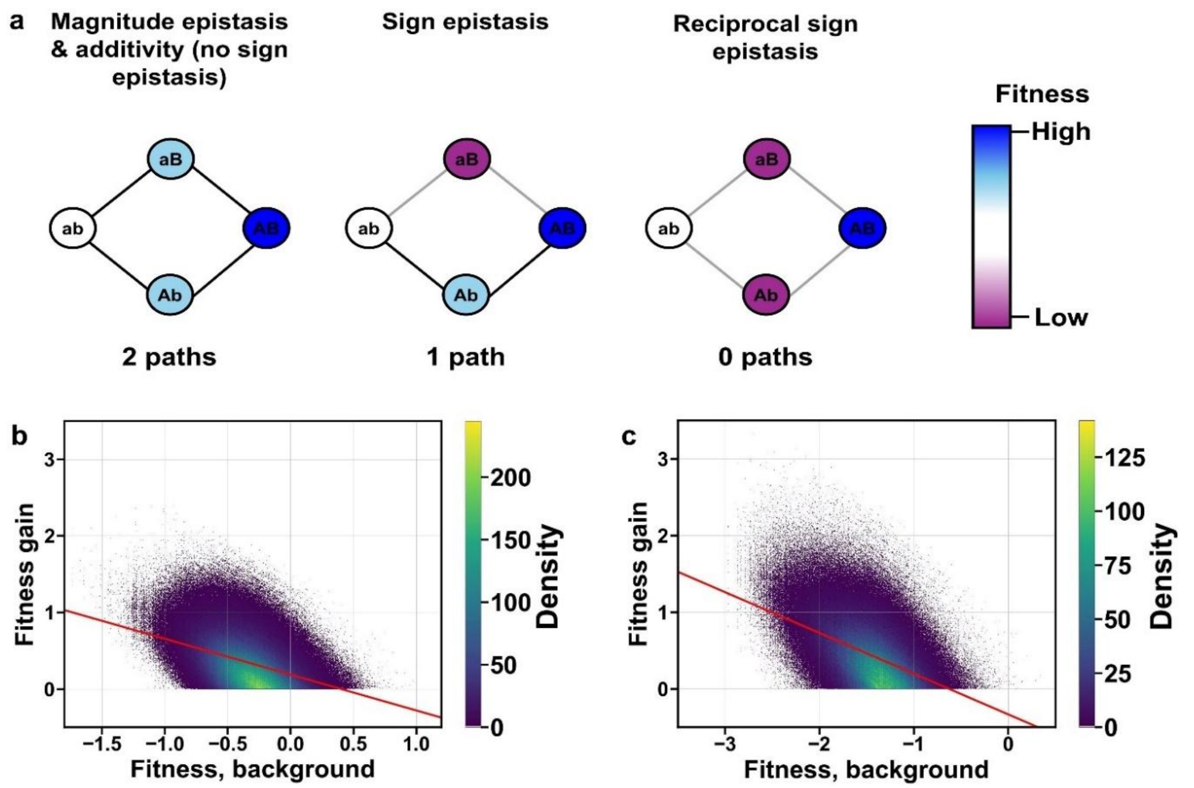

**Figure S36. Patterns of epistasis across different environments.** (a) Schematic representation of three types of epistasis using graph motifs as described previously (6). Each motif includes four genetic variants (circles), a double mutant (AB), the corresponding wild-type (ab), and two single mutants (Ab and aB), which are separated by a genetic distance of two nucleotides. Circle colors indicate fitness (see color legend). Lines between genotypes represent possible evolutionary paths of single mutations, where black lines indicate accessible paths from ab to AB, while gray lines denote inaccessible paths due to low-fitness intermediates (Ab and/or aB). The three types of epistasis differ based on the number of accessible paths between ab and AB. Frequency of the three types of epistasis in glycerol and lactose environments is listed in Table S8. (b-c) Evidence of diminishing returns epistasis in both environments. The density of points is represented by a color gradient, with lighter areas indicating higher density. The plots show that the fitness advantage of beneficial mutations (y-axis) decreases as the fitness of the genetic background in which a mutation occurs (x-axis) increases. A linear regression model (red line) illustrates this negative relationship. Panel (b) represents data from the glycerol environment (N=1,906,692 mutations), while panel (c) shows the same trend in lactose (N=1,131,786 mutations). The vertical axes do not admit values smaller than zero, because the analysis considers only mutations that increase fitness.
